# Hotspot *ESR1* mutations are multimodal and contextual drivers of breast cancer metastasis

**DOI:** 10.1101/2021.02.10.430701

**Authors:** Zheqi Li, Yang Wu, Megan E. Yates, Nilgun Tasdemir, Amir Bahreini, Jian Chen, Kevin M. Levine, Nolan M. Priedigkeit, Simak Ali, Laki Buluwela, Spencer Arnesen, Jason Gertz, Jennifer K. Richer, Benjamin Troness, Dorraya El-Ashry, Qiang Zhang, Lorenzo Gerratana, Youbin Zhang, Massimo Cristofanilli, Maritza A. Montanez, Prithu Sundd, Callen T. Wallace, Simon C. Watkins, Li Zhu, George C. Tseng, Nikhil Wagle, Jason S. Carroll, Paul Jank, Carsten Denkert, Maria M Karsten, Jens-Uwe Blohmer, Ben H. Park, Peter C. Lucas, Jennifer M. Atkinson, Adrian V. Lee, Steffi Oesterreich

## Abstract

Constitutively active estrogen receptor-α (ER/*ESR1*) mutations have been identified in approximately one third of ER+ metastatic breast cancer. Although these mutations are known mediators of endocrine resistance, their potential role in promoting metastatic disease has not yet been mechanistically addressed. In this study, we show the presence of *ESR1* mutations exclusively in distant, but not local recurrences. In concordance with transcriptomic profiling of *ESR1* mutant tumors, genome-edited Y537S and D538G cell models have a reprogrammed cell adhesive gene network via alterations in desmosome/gap junction genes and the *TIMP3/MMP* axis, which functionally confers enhanced cell-cell contacts while decreased cell-ECM adhesion. Context-dependent migratory phenotypes revealed co-targeting of Wnt and ER as vulnerability. Mutant ESR1 exhibits non-canonical regulation of several metastatic pathways including secondary transactivation and *de novo* FOXA1-driven chromatin remodeling. Collectively, our data supports evidence for *ESR1* mutation-driven metastases and provides insight for future preclinical therapeutic strategies.

**Significance:** Context and allele-dependent transcriptome and cistrome reprogramming in genome-edited *ESR1* mutation cell models elicit diverse metastatic phenotypes, including but not limited to alterations in cell adhesion and migration. The gain-of-function mutations can be pharmacologically targeted, and thus may be key components of novel therapeutic treatment strategies for ER-mutant metastatic breast cancer.

## Introduction

More than 70% of breast cancers express estrogen receptor-α (ER/*ESR1*). Antiestrogen therapies, including depletion of estradiol (E2) by aromatase inhibitors (AIs) or antagonizing ER activity by Selective Estrogen Receptor Modulators/Degraders (SERMs/SERDs), are conventional treatments for ER+ breast cancer. Development of resistance to these endocrine therapies, however, remains a clinical and socioeconomic challenge (1,2).

30–40% of endocrine-resistant metastatic breast cancer (MBC) is enriched in *ESR1* somatic base pair missense mutations (3–5), that can be detected in the blood of patients with advanced disease (6,7). Clinically, ligand binding domain (LBD) *ESR1* mutations correlate with poor outcomes in patients with advanced disease (6,8,9). Recent work from our group and others has uncovered a crucial role for these *ESR1* hotspot mutations in driving constitutive ER activity and decreased sensitivity towards ER antagonists (10–12). Moreover, structural investigation of the two most frequent mutations, variants Y537S and D538G, has demonstrated that *ESR1* mutations stabilize helix 12 (H12) in an agonist conformation, thereby providing a mechanistic explanation for constitutive ER activity (13).

The identification of *ESR1* mutations in endocrine resistant MBC suggests that mutant ER may not only mediate endocrine resistance but also have an unappreciated role in enabling metastasis. Indeed, recent *in vivo* studies showed that mutant ER can promote metastasis (14,15), and *in vitro* studies showed a gain of cell motility (15,16) and growth in 3D culture (17). Although epithelial-mesenchymal transition (EMT) has been described as one potential explanation for the Y537S mutant (18), overall mechanisms remain largely unclear. In order to identify personalized therapeutic vulnerabilities in patients harboring *ESR1* hotspot mutations, there is an urgent need to decipher the mechanistic underpinnings and precise roles of mutant ER in the metastatic progression using comprehensive approaches and model systems.

Previous transcriptomic profiling performed by us and others has revealed a context-dependence of *ESR1* mutation effects, as well as significant differences between the two most frequent hotspot mutations, Y537S and D538G (11,12,14,15,19). Differentially expressed genes vary widely following expression of the mutations in their respective cell line model, however, both Y537S and D538G maintain distinction from the E2-dependent wild-type (WT) ER transcriptome. Similarly, comparison of the WT and mutant ER cistromes has also revealed context-dependent and allele-specific effects on ER recruitment (11,14). Furthermore, we recently showed that *ESR1*-mutant transcriptomic reprogramming is associated with epigenetic remodeling (19). While these findings imply that in the setting of high molecular diversity in tumors and patients, somatic *ESR1* mutations have the potential to trigger different metastatic phenotypes, this phenomenon has yet to be investigated.

In this study, we explore metastatic gain-of-function phenotypes in genome-edited *ESR1* mutant models under the guidance of transcriptomic changes detected in clinical samples. We identify mechanisms underlying context and allele-specific metastatic phenotypes, and subsequently confirm alterations in a number of potential therapeutic targets in metastatic tumors. We believe that our systematic bedside-to-bench approach will ultimately lead to improved metastasis-free outcomes and prognosis for patients with ER+ tumors.

## Results

### Significant enrichment of *ESR1* mutations in distant metastases compared to local recurrences

To establish clinical evidence for potential metastasis-conferring roles of *ESR1* LBD mutations, we compared the *ESR1* mutation frequencies between distant metastatic and locally recurrent tumors. A combination of four publicly available clinical cohorts (MSKCC, METAMORPH, POG570 and IEO) showed that while 156/867 distant metastases (18%) harbored *ESR1* mutations, none were found in the 38 local recurrence samples (Table 1 and Supplementary Table S1) (20–23). To expand upon this observation, we additionally screened 75 ER+ recurrent tumors from the Women’s Cancer Research Center (WCRC) and Charite Hospital for *ESR1* hotspot (Y537S/C/N and D538G) mutations using highly sensitive droplet digital PCR (ddPCR). We identified 12 *ESR1* mutation-positive cases among the distant metastases (25%), whereas none of the local recurrences were *ESR1* mutation-positive (Table 1 and Supplementary Table S2). Notably, there was no significant difference in time to recurrence for patients with distant vs local recurrences in four of the cohorts (Supplementary Fig. S1 & Table S3, data is not available for IEO cohort), excluding the possibility that the observed differences could simply be due to duration of time to recurrence, as was previously suggested (6).

**Table 1.** Significant enrichment of *ESR1* mutations in distant compared to local recurrences. Upper panel: Data from 867 distant metastatic and 38 local recurrence cases were merged from three cohorts (METAMORPH, 39 distant/9 local; POG570, 86 distant/14 local; MSKCC, 716 distant/8 local; IEO, 26 distant/7 local). *ESR1* mutation status was previously identified by whole exome sequencing (METAMORPH), whole genome sequencing (POG570) or target panel DNA sequencing (MSKCC, IEO). Lower panel: 48 distant ER positive metastases and 27 local ER positive recurrences were obtained from the WCRC and Charite cohorts. Genomic DNA (gDNA) was isolated from either FFPE or frozen tumor tissues, and subjected to droplet digital PCR (ddPCR) detection with specific probes against Y537S, Y537C, Y537N and D538G hotspot point mutations (cDNA rather than gDNA was used for 3 of the local recurrent samples). Hotspot *ESR1* mutation incidences between distant metastatic and local recurrent samples in both panels were compared using a Fisher’s exact test.

| Cohorts | Site of Recurrence | Total Number | *ESR1* WT | *ESR1* Mutant | Fisher's Exact p |
| --- | --- | --- | --- | --- | --- |
| METAMORPH/POG570/MSKCC/IEO Merged | Distant | 867 | 711 (82%) | 158 (18%) | 0.0014 |
|  | Local | 38 | 38 (100%) | 0 (0%) |  |
| WCRC/Charite | Distant | 48 | 36 (75%) | 12 (25%) | 0.0031 |
|  | Local | 27 | 27 (100%) | 0 (0%) |  |

### *ESR1* mutant tumors show a unique transcriptome associated with multiple metastatic pathways

To identify candidate functional pathways mediating the metastatic properties of *ESR1* mutant cells, we compared WT and *ESR1* mutant tumor transcriptomes from four cohorts of ER+ metastatic tumors: our local WCRC cohort (46 *ESR1* WT and 8 mutant tumors) (24–26) and three previously reported cohorts - MET500 (34 *ESR1* WT and 12 mutants tumors), POG570 (68 *ESR1* WT and 18 mutants tumors) and DFCI (98 *ESR1* WT and 32 mutants tumors) (14,22,27) (Fig. 1A & Supplementary Table S4).

**Figure 1.**
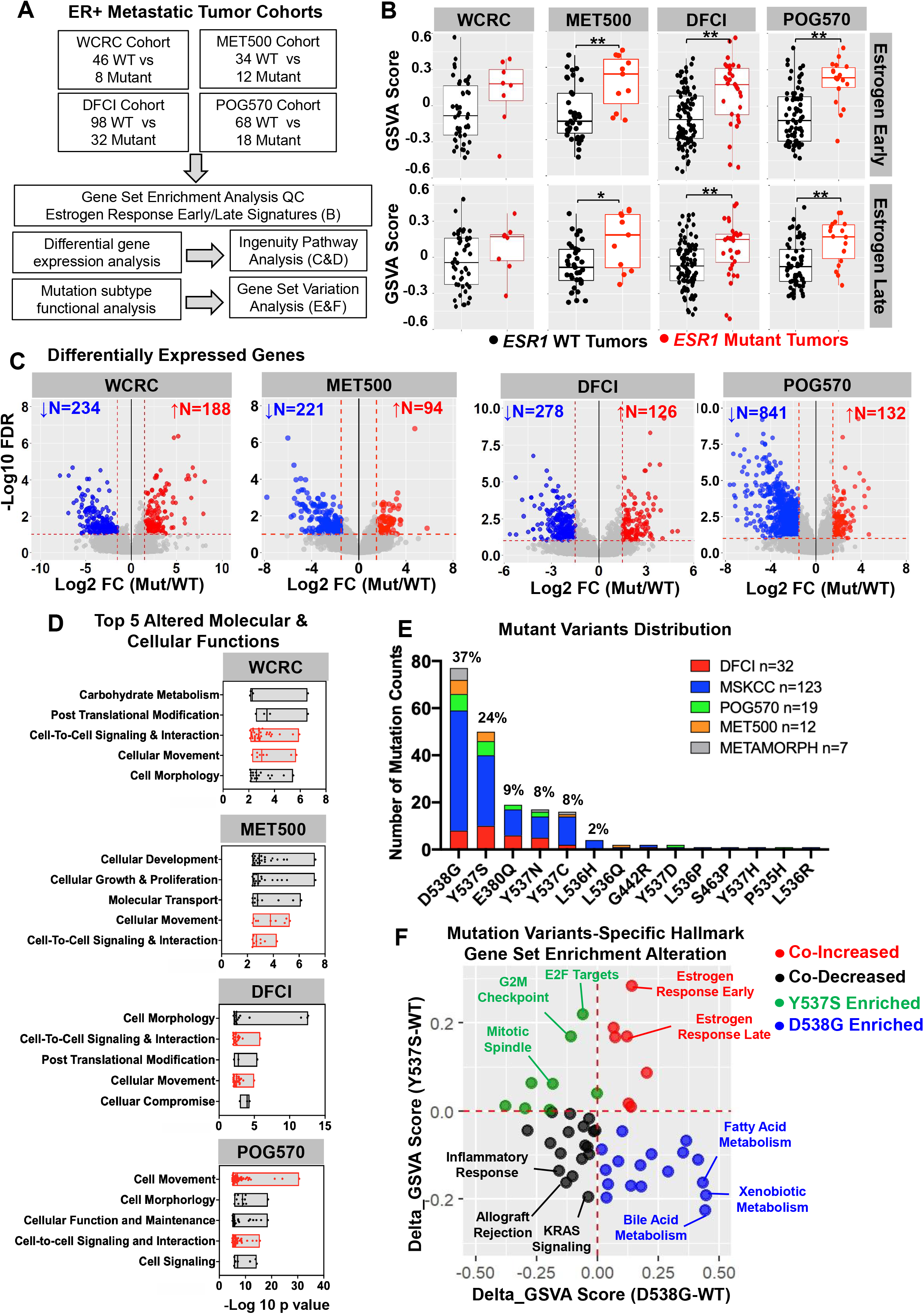
Transcriptomic landscape of *ESR1* mutant metastatic breast cancers. A. Schematic overview of transcriptomic analysis of four ER+ metastatic breast cancer cohorts. B. Box plots representing the enrichment levels of “Estrogen Response Early” and “Estrogen Response Late” signatures in *ESR1* mutant versus *ESR1* WT metastatic tumors in each cohort. (WCRC, 46 *ESR1* WT/8 mutant; MET500, 34 *ESR1* WT/12 *ESR1* mutant; DFCI, 98 *ESR1* WT/32 mutant; POG570, 68 *ESR1* WT/18 mutant). Four quantiles are shown in each plot. Mann-Whitney U test was used to compare the enrichment of the signatures in WT and mutant tumors. (* p<0.05, ** p<0.01) C. Volcano plots representing the differentially expressing genes (DE genes) in *ESR1* mutant tumors versus WT tumors in the three metastatic breast cancer cohorts. DE genes were selected using the cutoff of FDR<0.1 and |log_2_FC|>1.5. Genes that were upregulated or downregulated were labelled in red and blue respectively with corresponding counts. D. Dot plots showing the top 5 altered cellular and molecular functional categories derived from DE genes analysis using Ingenuity Pathway Analysis software. Specific sub-functions within overarching categories are presented as individual dots. Consistently altered pathways across all four cohorts are indicated in red. E. Stacked bar plot showing the distribution of 14 hotspot *ESR1* mutations identified in six independent cohorts using unbiased DNA sequencing approaches. Specific sample numbers were indicated in the plots. Variants with percentages above 1% were labelled on the top of each bar. F. Scatterplot representing enrichment level distribution of 50 hallmark gene sets in 10 Y537S and 8 D538G metastatic tumors (after being normalized against 98 WT counterparts) from the DFCI cohort. Top enriched pathways from each quartile are labelled.

Although principal component analyses on global transcriptomes did not segregate *ESR1* WT and mutant tumors (Supplementary Fig. S2A), both “Estrogen Response Early” and “Estrogen Response Late” signatures were significantly enriched in *ESR1* mutant tumors in 3 out of 4 cohorts, with a trend towards enrichment in the fourth cohort (Fig. 1B). These results recapitulate the observation of ER hyperactivation as a result of hotspot mutations, previously described in other preclinical studies (12,14,28). Differential gene expression analysis identified a considerable number of altered genes that were associated with *ESR1* mutations (Fig. 1C & Supplementary Table S5), which further inferred functional alterations in various metastasis-related pathways. Remarkably, “Cell-To-Cell Signaling & Interaction” and “Cell Movement” were featured among the top five altered pathways for *ESR1* mutant tumors in all four cohorts (Fig. 1D).

In addition to the broad effects associated with *ESR1* mutations, we next questioned whether different *ESR1* mutant variants could display divergent functions. A meta-analysis of the five above-mentioned ER+ MBC cohorts examining *ESR1* mutations underscored D538G (37%) and Y537S (24%) as the predominant variants (Fig. 1E). Given the challenge of merging RNA-seq data sets from multiple cohorts due to immense technical variations, we selectively compared mutation variant specific transcriptomes of ten Y537S- or eight D538G-harboring tumors to the WT counterpart (n=32) respectively from the DFCI cohort, which provided the largest numbers and thus maximized statistical power. Aligning enrichment levels of 50 hallmark gene sets for the two mutant variants again confirmed “Estrogen Response Early” and “Estrogen Response Late” as the top co-upregulated pathways (Fig. 1F), with Y537S tumors displaying higher ER activation (Supplementary Fig. S2B), consistent with cell line studies (12,29). We also identified enriched cell cycle related pathways (E2F targets, G2M checkpoint and mitotic spindle) and metabolic related pathways (fatty acid, bile acid and xenobiotic metabolisms) in Y537S and D538G tumors, respectively, implying that different *ESR1* mutant variants might hijack distinct cellular functions to promote malignancy. Taken together, these results provide support that despite mutant variant-specific alterations, *ESR1* mutations might broadly mediate metastatic phenotypes through effects on cell-to-cell interactions and cell movement. We next validated the *in silico* results using previously established genome-edited MCF7 and T47D cell line models (12).

### *ESR1* mutant-cells exhibit stronger cell-cell adhesion

We first addressed the enrichment of cell-cell interaction signaling in the mutant tumors through morphological inspection of cell cluster formation in suspension culture (Fig. 2A). We observed more compact cell clusters in MCF7 and T47D mutant cell lines compared to their WT counterparts after six days of suspension culture. A time course study confirmed enhanced cluster formation 24-48hrs past cell seeding (Supplementary Fig. S3A). Similar observations were made in individual clones, eliminating the possibility for clonal effects (Supplementary Fig. S3B).

**Figure 2.**
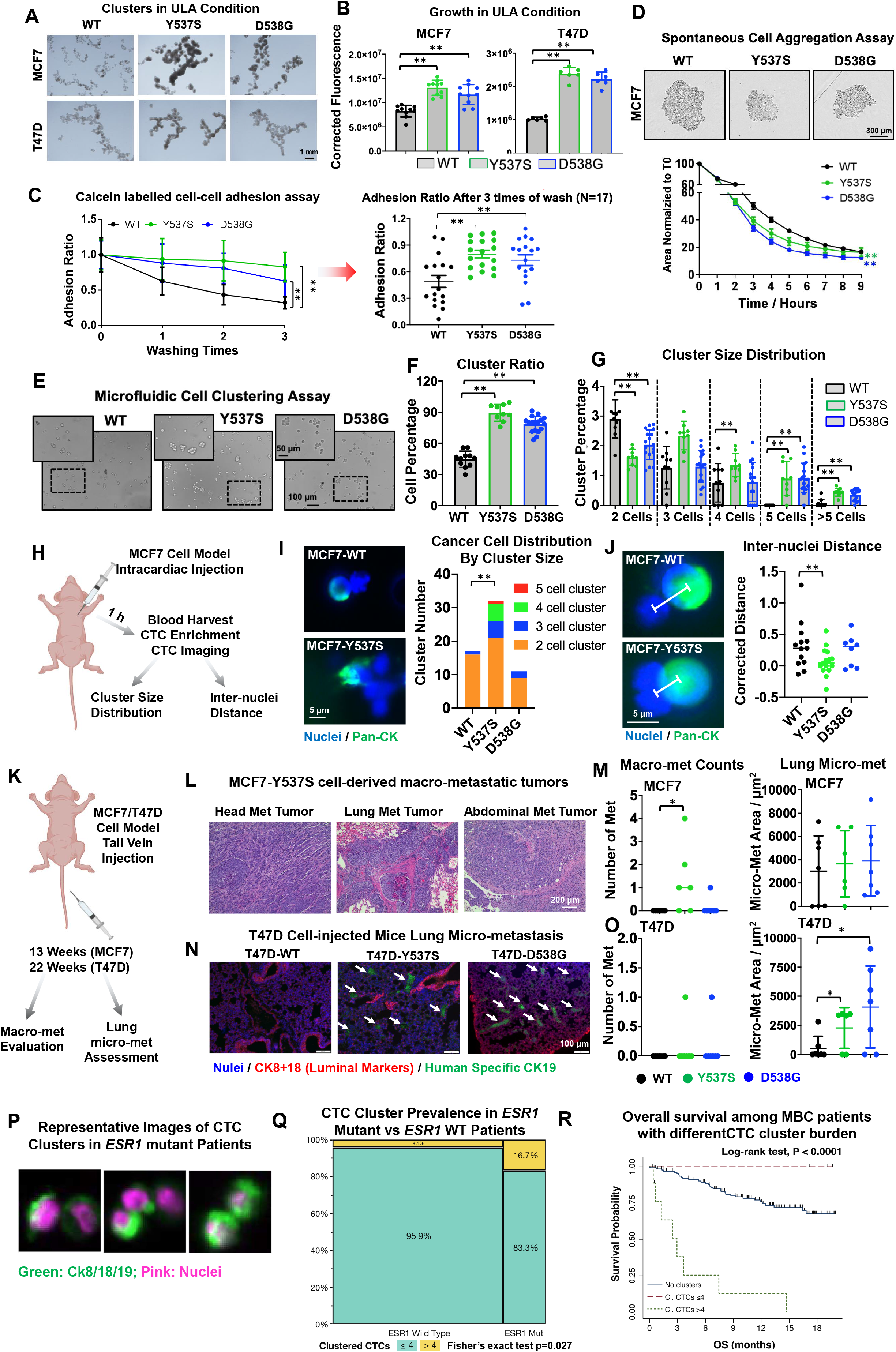
*ESR1* mutant cells exhibit stronger cell-cell adhesion. A. Representative images of day 6 hormone deprived MCF7 and T47D spheroids seeded in 6-well ultra-low attachment (ULA) plates. Images were taken under 1.25x magnification. Representative experiment from three independent repeats is shown. B. Bar plot representing day 7 cell numbers of MCF7 or T47D WT and *ESR1* mutant cells seeded into flat bottom ULA plates. Cell abundance were quantified using Celltiter Glo. Fluorescence readouts were corrected to background measurements. Each bar represents mean ± SD with 10 (MCF7) or 6 (T47D) biological replicates. Representative experiment from six independent repeats is shown. Dunnett’s test was used between WT and each mutant. (** p<0.01) C. Left panel: A calcein labelled cell-cell adhesion assay was performed in MCF7 WT and mutant cells. Adhesion ratios were calculated by dividing the remaining cells after each wash to the initial readout from unwashed wells. A pairwise two-way ANOVA between WT and each mutant was utilized. Each point represents mean ± SD with five biological replicates. Representative experiment from 17 independent repeats is shown. Right panel: Adhesion ratios after three washes were extracted from 17 independent experiments displayed as mean ± SEM. Dunnett’s test was used to compare between WT and each mutant. (* p<0.05, ** p<0.01) D. Line plot representing the aggregation ratio of MCF7 cells seeded into round bottom ULA plates. Cell aggregation processes were followed by the IncuCyte living imaging system every hour. Spheroid areas were normalized to time 0. Each dot represents mean ± SD with eight biological replicates. Representative images after 3 hours of aggregation are shown across the top panel. Images were captured under 10x magnification. Representative experiment from five independent repeats is shown. A pairwise two-way ANOVA between WT and each mutant was utilized. (** p<0.01) E. Representative images of MCF7 cell cluster status after two hours of flow under physiological shear stress produced by the ibidi microfluidic system. Images were taken under 10x magnification. A regional 2x zoom in is presented on the top of each image. Representative experiment from three independent repeats is shown. F. Bar graph representing the percentage of MCF7 cells in a cluster based on the quantification of cluster and single cell numbers from 12 representative images per group. Each bar represents mean ± SD. Cell cluster ratios after 2 hours of flow were further normalized to time 0 to correct for baseline pre-existing clusters. Representative experiment from three independent repeats is shown. Dunnett’s test was used between WT and mutant cells. (** p<0.01) G. Bar plots showing the cluster size distribution of MCF7 cells after normalization to time 0. Each bar represents mean ± SD from 12 representative images per group. Representative experiment from three independent repeats is shown. Dunnett’s test was used between WT and each mutant cell type within the same cluster size category. (** p<0.01). H. Schematic overview of short-term *in vivo* circulating tumor cell evaluation experimental procedure. I. Left panel: Representative images of two-cell clusters (WT) and a multicellular cluster (Y537S). Images were taken under 40x magnification. Right panel: Stacked bar chart representing the distribution of cancer cells in each cluster type. This experiment was performed once. Fisher’s exact test was applied to test whether multicellular clusters were enriched in *ESR1* mutant cells. (** p<0.01) J. Left panel: Representative images of a WT and Y537S two cell cluster. Lines connecting the two nuclei centers were indicated. Images were taken under 40x magnification. Right panel: Dot plot represents the inter-nuclei distance of all two-cell clusters in MCF7 WT and mutant cells. Measured distances were normalized to the average radius of both cells of this cluster size to avoid cell size bias. This experiment was performed once. Mann-Whitney U test was performed between WT and each mutant cell. (** p<0.01) K. Schematic overview of *in vivo* metastatic evaluation of *ESR1* mutant cells introduced via tail vein injections. L. Representative H&E staining images the tumorous portion of MCF7-Y537S induced macro-metastatic (macro-met) tumors from 3 different mice. This experiment was performed once. Images were taken under 20x magnification. M. Left panel: Dot plots showing the number of macro-met per mouse from MCF7 *ESR1* WT and mutant cells-injected mice. Pairwise Mann-Whitney U test was used to compare the macro-met numbers in each mutant group to WT cell-injected groups. Right panel: Quantification of lung micro-met areas based on human specific CK19 staining quantification. This experiment was performed once. Pairwise Mann-Whitney U test was applied for statistical analysis. (WT, n=7; Y537S, n=6; D538G, n=7) (* p<0.05) N. Representative images of micro-metastatic loci on the lung sections of T47D-*ESR1* mutant cell-injected mice. Images were taken under 10x magnification. Metastatic loci were indicated with white arrow. This experiment was p once. (WT, n=7; Y537S, n=6; D538G, n=7) (Blue: nuclei; Red: CK8+18; Green: Human specific CK19) O. Left panel: Dot plots showing the macro-metastatic counts per mouse from T47D *ESR1* mutant-injected mice. Pairwise Mann-Whitney U test was used to compare the macro-met numbers in each mutant group to WT cell-injected groups. Right panel: Quantification of lung micro-met areas based on CK19 staining and was performed in a blind manner. This experiment was performed once. Pairwise Mann-Whitney U test was applied for statistical analysis. (N=1, * p<0.05) P. Representative images of CTCs clusters detected through the CellSearch Platform after EpCAM dependent enrichment (Pink: nuclei, Green: CK8/CK18/CK 19). Image resolution and magnification were achieved in accordance with the CellSearch Platform. Q. Mosaic plot showing the association between *ESR1* genotype status and clustered CTCs. A significant positive association was observed by Fisher’s exact test between *ESR1* mutations and high CTC cluster burden. (CTC cluster > 4). R. Kaplan Meier plot representing the impact of clustered CTCs in terms of Overall Survival (OS). Patients with clustered CTCs > 4 experienced the worse prognosis in terms of OS both with respect to those without clusters and those with clusters but with ≤ 4 clustered CTCs (P < 0.0001). Patients at risk are reported at each time point. Log rank test was to compare the survival curves of the two patient subsets.

Since *ESR1* mutant cells displayed significantly increased ligand-independent growth in suspension (Fig. 2B), we sought to rule out the possibility that increased cluster formation was simply a result of increased cell number by assessing cell-cell adhesive capacity using multiple approaches in short term culture (within 1 day). We therefore directly quantified homotypic cell-cell interactions by measuring the adhesion of calcein-labelled *ESR1* WT or mutant cells. This assay showed that both MCF7 mutant cells exhibited significantly stronger cell-cell adhesion compared to the WT cells (Fig. 2C). In T47D cells, a similar effect was observed, but was limited to the T47D-Y537S mutant cells (Supplementary Fig. S4A). These assays were complemented by quantification of cell aggregation rates as a direct reflection of cell-cell adhesion, which confirmed faster aggregation in MCF7-Y537S/D538G and T47D-Y537S cells (Fig. 2D & Supplementary Fig. S4B-S4D). In addition, these stronger cell-cell adhesive properties were also reproduced in additional *ESR1* mutant cell models from other laboratories (19,28) (Supplementary Fig. S4E and S4F).

Cell-cell interaction has been reported to affect several stages of metastasis, including collective invasion, intravasation, dissemination and circulation (30–32). To test whether ER mutations may affect tumor cell-cell adhesion in circulation, we utilized a microfluidic pump system to mimic arterial shear stress. Comparing representative images before and after 2 hours of microfluidic flow, we found MCF7 *ESR1* mutant cells had a greater tendency to aggregate together (Fig. 2E and 2F). Larger clusters comprised of five or greater cells were more prevalent in the *ESR1* mutant cell lines, whereas smaller two-cell clusters were diminished (Fig. 2G). A similar phenotype was also identified in additional MCF7 *ESR1* mutant cells and in our T47D-Y537S cell line (Supplementary Fig. S5A-S5I), consistent with our observations in static conditions. In an additional orthogonal approach, we utilized a quantitative microfluidic fluorescence microscope system simulating blood flow (33). Quantification of dynamic adhesion events normalized to adhesion surfaces revealed a consistent enhanced cell-cell adhesion capacity of *ESR1* mutant MCF7 cells (Supplementary Fig. S5J-S5K, Supplementary videos 1-3). Together, these results show that hotspot *ESR1* mutations confer increased cell-cell attachment under static and fluidic conditions, and that the effect size is dependent upon mutation type and genetic backgrounds. These findings are at odds with increased EMT features (18), and indeed the majority of *ESR1* mutant models and tumors did not show increased EMT signature or increased expression of EMT marker genes (Supplementary Fig. S6).

We next sought to assess whether this unexpected phenotype translated into numbers of CTC clusters and subsequent metastasis *in vivo*. One hour post intracardiac injection into athymic mice, circulating MCF7 WT and mutant cells were enriched from blood using a previously described electrical CTC filtering method (34) (Fig. 2H). 41%-81% of CTC clusters were composed of both cancer and non-cancer cells (Supplementary Fig. S7A). Despite no difference in the average amount of single CTCs and CTC clusters per mouse between the WT and mutant *ESR1* (Supplementary Fig. S7B & S7C), we found that overall MCF7-Y537S mutant cells were significantly enriched in clusters with greater than 2 cells (Fig. 2I). Furthermore, quantification of inter-nuclei distances between two-cell clusters revealed denser MCF7-Y537S clusters (Fig. 2J), supporting stronger MCF7-Y537S cell-cell interactions in an *in vivo* blood circulation environment. The data from the MCF7-D538G mutant cells did not recapitulate the adhesive phenotype we discerned *in vitro*, suggesting mutation site-specific interactions with the *in vivo* microenvironment potentially affect cluster formation.

We next performed tail vein injection and monitored bloodborne metastatic development in longer-term *in vivo* experiments without estradiol supplement (Fig. 2K). We observed multiple distant macro-metastatic tumors in 4/6 (67%) MCF7-Y537S mutant cell-injected mice (Fig. 2L). In contrast, distant macro-metastatic tumor was observed in only one mouse of MCF7-D538G group (1/7) and none in MCF7-WT group (0/7) (Fig. 2M, left panel). We detected no difference in lung micro-metastatic foci areas between WT and mutant cell-injected mice, potentially due to a high baseline of MCF7 lung colonization capacity (Fig. 2M, right panel). In contrast to our MCF7 results, we only discerned macro-metastatic tumors from each T47D mutant group (Y537S: 1/6; D538G: 1/7) and none in T47D-WT group (0/7) after 23 weeks of injection (Fig. 2O, left panel), underpinning its less aggressive behavior as compared to MCF7 cells (35,36). However, both T47D-Y537S and T47D-D538G mutant cells resulted in enlarged lung micro-metastases, with a more pronounced effect in the T47D-D538G cells (Fig. 2N and 2O, right panel). Interestingly, our *in vitro* assays did not suggest altered cell-cell adhesion in the T47D-D538G model, suggesting the potential use of alternative mechanisms to strengthen its metastatic properties *in vivo*.

Encouraged by our *in vitro* and *in vivo* findings, we next examined CTC clusters in patients with *ESR1* mutant tumors. Taking advantage of a recent CTC sequencing study (37), we sought to generate CTC cluster gene signatures. Differential gene expression analysis in two patients with ER+ disease who had at least two CTC clusters and single CTCs sequenced identified CTC cluster enriched genes (Supplementary Fig. S8A and Table S6), which we subsequently applied to our RNA-seq dataset with 51 pairs of ER+ primary-matched metastatic tumors (44 *ESR1* WT and 7 mutant) merged from the WCRC and DFCI cohorts. *ESR1* mutant metastatic tumors exhibited significantly higher enrichment of CTC cluster-derived gene signatures (Supplementary Fig. S8B).

To examine the interplay between *ESR1* mutations status, numbers of CTCs, and clinical outcome, we analyzed a cohort of 151 patients with MBC. Median age at the first blood draw for CTCs enumeration was 55 years (IQR: 44 - 63 years), 63 patients (45.7%) were diagnosed with ER+ MBC, 37 (26.8%) with HER2-positive MBC and 38 (27.5%) with TNBC. Bone (49.7%), lymph nodes (41.1%), lung (34.4%) and liver (34%) were the most common sites of metastasis (Supplementary Table S7). Median number of CTCs was 1 (IQR: 0-10), clusters were detectable in 14 patients (9.3%) (Fig. 2P) and in this subgroup the median number of clustered CTCs was 15.5 (IQR: 4 - 20). Classifying the MBC by CTC numbers, with CTC >=5/7.5ml blood being more aggressive, and CTC<5/7.5 ml blood more indolent, there were 101 Stage IV indolent (69.9%) and 50 Stage IV aggressive cases (33.1%) in the study. If cases were classified by presence of CTC clusters in blood, there were 10 (6.6%) and 141 (93.4%) cases with >4 CTC clusters and ≤4 clusters, respectively. (Supplementary Table S7). Mutations in hotspots D538 and Y537 of *ESR1* were detected in 30 patients (19.9%), while mutations in hotspots E453 and H1047 of *PIK3CA* were detected in 40 patients (26.5%) (Supplementary Table S7). A significant association was observed between *ESR1* genotype status and clustered CTCs > 4 (P = 0.027) (Fig. 2Q), while no association was observed with respect to *PIK3CA* (P=0.725). Notably, patients with > 4 CTCs clusters experienced the worse prognosis in terms of OS (6 months OS: 12.7%) both with respect to those without clusters (6 months OS: 88.5%) and those with clusters but with ≤ 4 clustered CTCs (6 months OS: 100%) (P < 0.0001) (Fig. 2R).

### Mutant *ESR1* cells show increased desmosome gene and gap junction gene families

To elucidate the mechanism of enhanced cell-cell adhesion, we investigated the enrichment of four major cell-cell junction subtypes – desmosomes, gap junctions (connexons), tight junctions and adherens junctions within the cell model RNA-seq data (12) (Supplementary Table S6). Enrichment of the desmosome gene and gap junction gene families was observed in both MCF7-Y537S/D538G and T47D-Y537S cells (Fig. 3A). Tight junctions were enriched in WT cells, and there were no differences in the adherens junction gene family expression (Supplementary Fig. S9A). Individual gene expression analysis (FC>1.2, p<0.05) identified 18 commonly upregulated desmosome genes and 4 gap junction genes in both MCF7 *ESR1* mutant cell lines (Fig. 3B). In addition to keratins, induction of classical desmosome genes *DSC1/2, DSG1/2* and *PKP1*, and gap junction genes *GJA1, GJB2* and *GJB5* were observed and validated by qRT-PCR in MCF7 cells (Fig. 3D). Higher protein levels were also observed for *DSC1, DSG2, PKP1, GJA1* (Cx43), and *GJB2* (Cx26) (Fig. 3C). Immunofluorescence staining revealed significantly higher *DSG2* expression in MCF7-Y537S at cell-cell contact surfaces, with a trend observed in MCF7-D538G (Fig. 3E). Consistent with the weaker *in vitro* cell-cell adhesion phenotypes in T47D mutant cells, we observed less pronounced desmosome and gap junction gene expression changes in T47D-Y537S cells (Supplementary Fig. S9B). We validated the overexpression of the key desmosome and gap junction genes in RNA-seq datasets from seven additional *ESR1* mutant cell models and performed further validation studies in two of them (Supplementary Fig. S9C-S9E) (11,15,19). Moreover, mining RNA-seq data from recently reported *ESR1* WT and mutant *ex vivo* CTC models (38), we observed overexpression of three gap junction and desmosome genes in the *ESR1* mutant CTC lines (Supplementary Fig. S9F). Finally, the top upregulated desmosome and gap junction genes (Supplementary Table S6) were also found significantly enriched in intrapatient matched primary and metastatic lesions with *ESR1* mutations (Fig. 3F).

**Figure 3.**
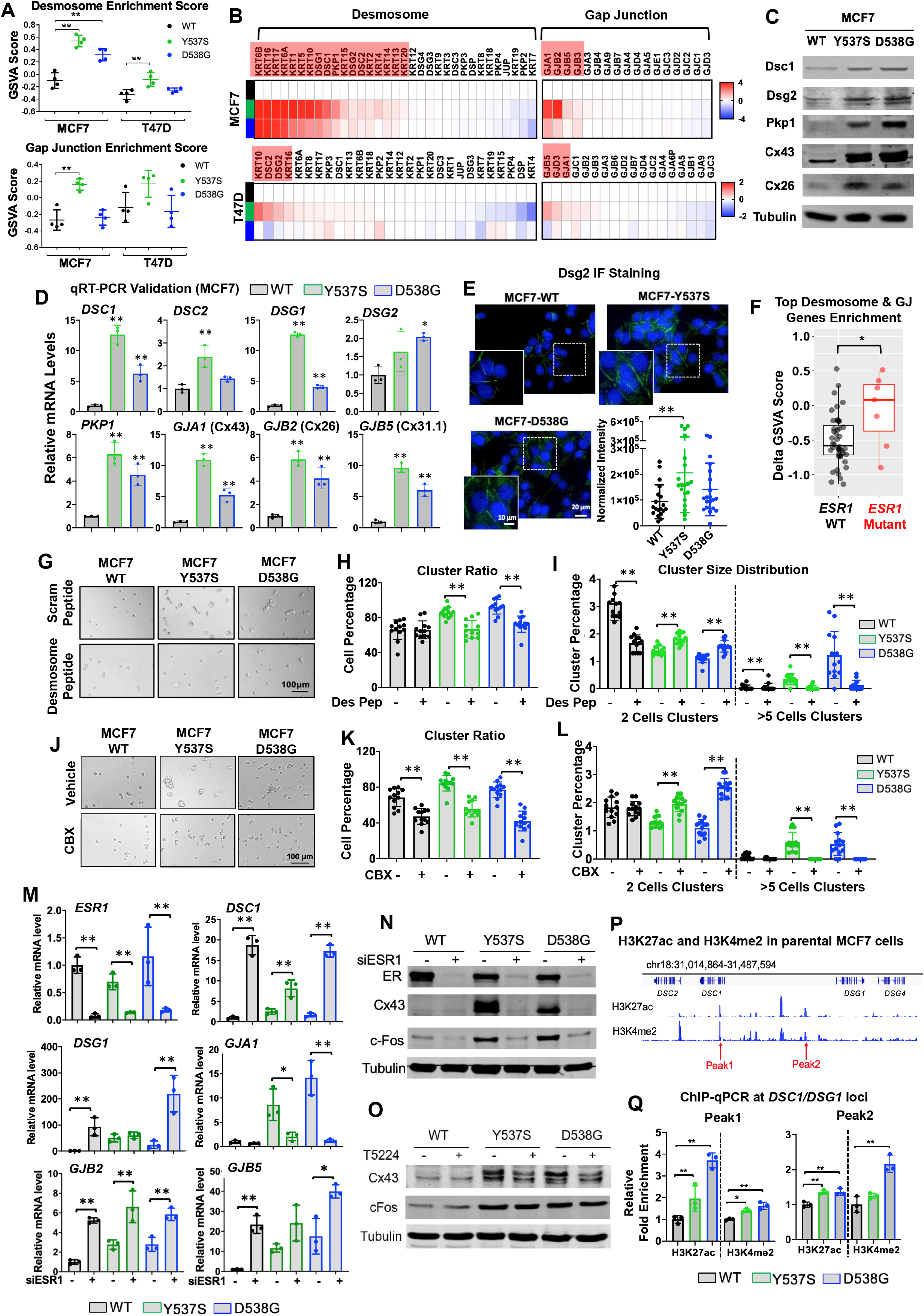
Desmosome and gap junction adhesome reprogramming confers enhanced adhesive properties in *ESR1* mutant cells. A. Gene Set Variation Analysis (GSVA) scores of desmosome and gap junction gene sets enrichment in MCF7 and T47D *ESR1* mutant vs WT cell RNA-seq data sets. Each cell type has four biological replicates. Dunnett’s test was used to test the significance between WT and mutant cell lines. (** p<0.01) B. Heatmaps showing all desmosome and gap junction component genes in MCF7 and T47D *ESR1* mutant cells. Data were extracted from RNA-sequencing results with four biological replicates. Color scale represents the Log2 fold changes in each mutant normalized to WT counterparts using the log_2_(TPM+1) expression matrix. Genes with counts=0 in more than one replicate in each cell type were filtered out of analysis. Genes with a log_2_FC>1.2 and a p<0.05 in at least one group are labelled in red. C. Western blot validation of the expression level of *DSG2, DSC1, PKP1*, Cx43 and Cx26 in MCF7 WT and *ESR1* mutant cells after hormone deprivation. Tubulin was blotted as a loading control. Representative blots from three independent repeats was shown for each protein. D. qRT-PCR validation of selected altered candidate desmosome and gap junction genes in MCF7 *ESR1* mutant cells. ΔΔCt method was used to analyze relative mRNA fold changes normalized to WT cells and *RPLP0* levels were measured as an internal control. Each bar represents mean ± SD with biological triplicates. This experiment was a representative from four independent repeats. Dunnett’s test was used to compare the gene expression between WT and each mutant. (* p<0.05, ** p<0.01) E. Representative images of immunofluorescence staining showing the distribution of desmoglein 2 (*DSG2*) in MCF7 WT and *ESR1* mutant cells. Images were taken under 20x magnification. A 2x zoom in of each image is presented. Right lower panel: *DSG2* signal intensities were quantified and normalized to cell numbers in each image. Data from 20 regions within the collected images were combined from four independent experiments. Mean ± SD is presented in each plot. Dunnett’s test was used to test the significance between WT and mutant cells. (** p<0.01) F. Box plots representing GSVA scores of the enrichment of the top desmosome and gap junction candidate genes (genes with log_2_FC>2 in at least one mutant line) in patient matched primary-metastatic paired samples. Delta GSVA score of each sample was calculated by subtracting the scores of primary tumors from the matched metastatic tumors. Four quantiles are shown in each plot. Mann-Whitney U test was performed to compare the Delta GSVA scores between *ESR1* WT (n=44) and mutation (n=7) harboring tumors. (* p<0.05) G & J. Representative images of cell cluster status after two hours of flow under physiological shear stress in the ibidi microfluidic system, with or without 300μM of the desmosomal blocking peptide (G) or 100μM of carbenoxolone (J) treatment. Images were taken under 10x magnification. This experiment was a representative from two (desmosome peptide treatment) and three (CBX treatment) independent repeats. H & K. Bar graphs representing the T0 normalized percentage of cells in cluster status after quantification of cluster and single cell numbers under each treatment. Each bar represents mean ± SD quantified from 12 images per group. This experiment was a representative from two (desmosome peptide treatment) and three (CBX treatment) independent repeats. Student’s t test was used to examine the effects of treatment between each group’s cluster ratio. (** p<0.01) I & L. Bar graphs representing the T0 normalized 2 cell and greater than 5 cell cluster percentages under each treatment. Each bar represents mean ± SD quantified from 12 images per group. This experiment was a representative from two (desmosome peptide treatment) and three (CBX treatment) independent repeats. Pairwise student’s t test was used to examine the effects of treatment between each group’s cluster ratio. (** p<0.01) M. Bar graphs representing qRT-PCR measurement of *DSC1, DSC2, GJA1, GJB2* and *GJB5* mRNA levels in MCF7 WT and *ESR1* mutant cells following siRNA knockdown of *ESR1* for 7 days. ΔΔCt method was used to analyze relative mRNA fold changes normalized to WT cells and *RPLP0* levels were measured as an internal control. Each bar represents mean ± SD with three biological replicates. Representative experiment from three independent repeats is displayed. Student’s t test was used to compare the gene expression between scramble and knockdown groups of each cell type. (* p<0.05, ** p<0.01) N & O. Western blot validation of the expression level of ER, Cx43 and cFOS in MCF7 WT and *ESR1* mutant cells after seven days of *ESR1* knockdown (N) or three days of 20μM T-5224 treatment (O). Tubulin was blotted as a loading control. Representative blot from three (N) and five (O) independent repeats is displayed. P. Screen shot of H3K27ac and H3K4me2 binding peaks at proximity to genomic *DSC1* and *DSG1* loci in MCF7 parental cells. ChIP-seq data were visualized at WashU Genome Browser based on public available data set from ENCODE (H3K4me2: ENCSR875KOJ; H3K27ac: ENCSR752UOD). Y axis represents the binding intensity of each ChIP-seq data set. Selected peaks for ChIP-qPCR assessment in Q were indicated. Q. Bar graph showing the fold enrichment levels of the two active histone modification markers at the two selected peaks around *DSC1* and *DSG1* gene loci illustrated in P. Each bar represents mean ± SD from biological triplicates. Fold enrichment levels were calculated by normalizing to IgG controls and further normalized to WT levels. This experiment is representative from two independent repeats. Dunnett’s test was used within each group. (N=2, * p<0.05, ** p<0.01)

We next investigated the functional roles of the reprogrammed adhesome in the *ESR1* mutant MCF7 cells. Transient individual knockdown of *DSC1, DSC2, GJA1 or GJB2* did not cause significant changes in adhesion in either *ESR1* mutant line (Supplementary Fig. S10A). However, we found compensatory effects observed in the desmosome and gap junction knockdowns as exemplified by increased *GJA1* levels after *DSC1* or *DSC2* knockdown (Supplementary Fig. S10B). The adhesive phenotype was disrupted, however, with an irreversible pan-gap junction inhibitor, Carbenoxolone (CBX), or with blocking peptide cocktails against desmocollin1/2 and desmoglein1/2 proteins. Both treatments caused significant inhibition of cell-cell aggregation in static conditions (Supplementary Fig. S10C & S10D) as well as diminished cluster propensities and size in microfluidic conditions (Fig. 3G–3L), suggesting redundancy in the mutant-driven reprogrammed desmosome and connexon pathways. In summary, MCF7-Y537S/D538G and T47D-Y537S mutants showed increased expression of desmosome and gap junction gene family components, which contributes to our observed enhanced cell-cell adhesion phenotype.

We next investigated the mechanisms underlying the elevated desmosome and gap junction components in *ESR1* mutant cells. Because hotspot *ESR1* LBD mutations are well-described as conferring constitutive ER activation, we first examined if these cell-cell adhesion target genes are direct outcomes of ligand-independent transcriptional programming. Interrogating publicly available RNA-seq and microarray datasets of six estrogen treated ER+ breast cancer cell lines (12,39–41), we found limited and inconsistent E2 induction of all examined cell-cell adhesion genes when compared to classical E2 downstream targets such as *GREB1* and *TFF1* (Supplementary Fig. S11A). Surprisingly, mining our MCF7 *ESR1* mutant cell model ER ChIP-seq data (42) showed an absence of proximate Y537S or D538G mutant ER binding sites (± 50kb of TSS) at desmosome and connexon target gene loci. These results suggest that the reprogrammed cell-cell adhesome is not a direct consequence of mutant ER genomic binding.

We therefore hypothesized that these altered adhesion target genes might be regulated via a secondary downstream effect of the hyperactive mutant ER. A seven-day siRNA ER knockdown assessment identified *GJA1* as the only target gene that could be blocked in mutant cells following ER depletion, whereas, strikingly, *DSC1, DSG1, GJB2* and *GJB5* mRNA levels were increased in all cell lines (Fig. 3M). This was congruent with *ESR1* knockdown in five additional ER+ parental cell lines, with the majority exhibiting a decrease in *GJA1* expression levels (Supplementary Fig. S11B). To unravel potential intermediate transcription factors (TFs) involved in the secondary regulation, we examined the levels of TFs previously reported to regulate *GJA1* expression (43) (Supplementary Fig. S11C). Among those, the AP1 family component *FOS* (cFos) was identified as the top TF upregulated in *ESR1* mutant cells in a ligand-independent manner. In addition, the AP1-associated transcriptional signature was also significantly enriched in MCF7 *ESR1* mutant cells (Supplementary Fig. S11D), and hence we tested if *GJA1* overexpression was dependent on the cFOS/AP1 transcriptional network. Higher cFOS mRNA and protein levels in *ESR1* mutant cells were confirmed, which declined along with *GJA1* levels after *ESR1* knockdown (Fig. 3N & Supplementary Fig. S11E). Importantly, pharmacological inhibition of cFOS-DNA binding partially rescued *GJA1* overexpression in *ESR1* mutant cells (Fig. 3O, Supplementary Fig. S11F-S11G). In conclusion, our results denote *GJA1* as an indirect target of mutant ER through activation of the cFOS/AP1 transcriptional axis in MCF7 cell models.

Since the majority of the cell-cell adhesion targets altered in the *ESR1* mutant cells were not direct ER target genes (Supplementary Fig. S11A & S11B), we investigated potential impacts of epigenetic remodeling on these targets. Using our recently reported ATAC-seq dataset from T47D *ESR1* mutant cells (19), we observed that one of the connexon targets, *GJB5*, exhibited increased chromatin accessibility at its gene locus in T47D-Y537S cells (Supplementary Fig. S12A & S12B), suggesting that epigenetic activation modulates gene expression in this particular context. We further evaluated active histone modifications on our target gene loci in the MCF7 model. We observed enhanced H3K27ac and H3K4me2 recruitment in both MCF7-Y537S and D538G cells at the nearest two histone modification sites around the *DSC1* and *DSG1* loci, the two most upregulated desmosome component genes in MCF7 mutant cells (Fig. 3P), suggesting activation of desmosome genes via an indirect ER-mediated epigenetic activation (Fig. 3Q).

### *ESR1* mutations promote reduced adhesive and enhanced invasive properties via altered *TIMP3-MMP* axis

In addition to altered cell-cell adhesion, metastasis is also mediated by coordinated changes in cell-matrix interaction (44,45). Therefore, we assessed whether mutant ER affects interaction with the extracellular matrix (ECM). Computational analysis showed inverse correlation between ECM receptor pathway signatures and *ESR1* mutation status in the DFCI cohort with the same trend appearing in 2/3 of the remaining cohorts (Fig. 4A, Supplementary Fig. S13A & Table S6). Employing an adhesion array on seven major ECM components, we observed that the MCF7 *ESR1* mutant cell lines consistently lacked adhesive properties on almost all ECM components with the exception of fibronectin, and T47D *ESR1* mutant cells displayed reduced adhesion on collagen I, collagen II and fibronectin (Fig. 4B). Considering that collagen I is the most abundant ECM component in ER+ breast cancer (Supplementary Fig. S13B), we repeated the adhesion assay on collagen I (Fig. 4C & 4D; Supplementary Fig. S13C & S13D) and similarly found reduced adhesion in both ER mutant cells. In an orthogonal approach, we visualized and quantified adhesion in a co-culture assay on collagen I using differentially labelled *ESR1* WT and mutant cells, which confirmed significantly decreased adhesive properties in the mutant cells (Supplementary Figure S13E & S13F). Of note, *ESR1* mutant adhesion deficiency on collagen I was also observed in two additional *ESR1* mutant models (Supplementary Fig. S13G).

**Figure 4.**
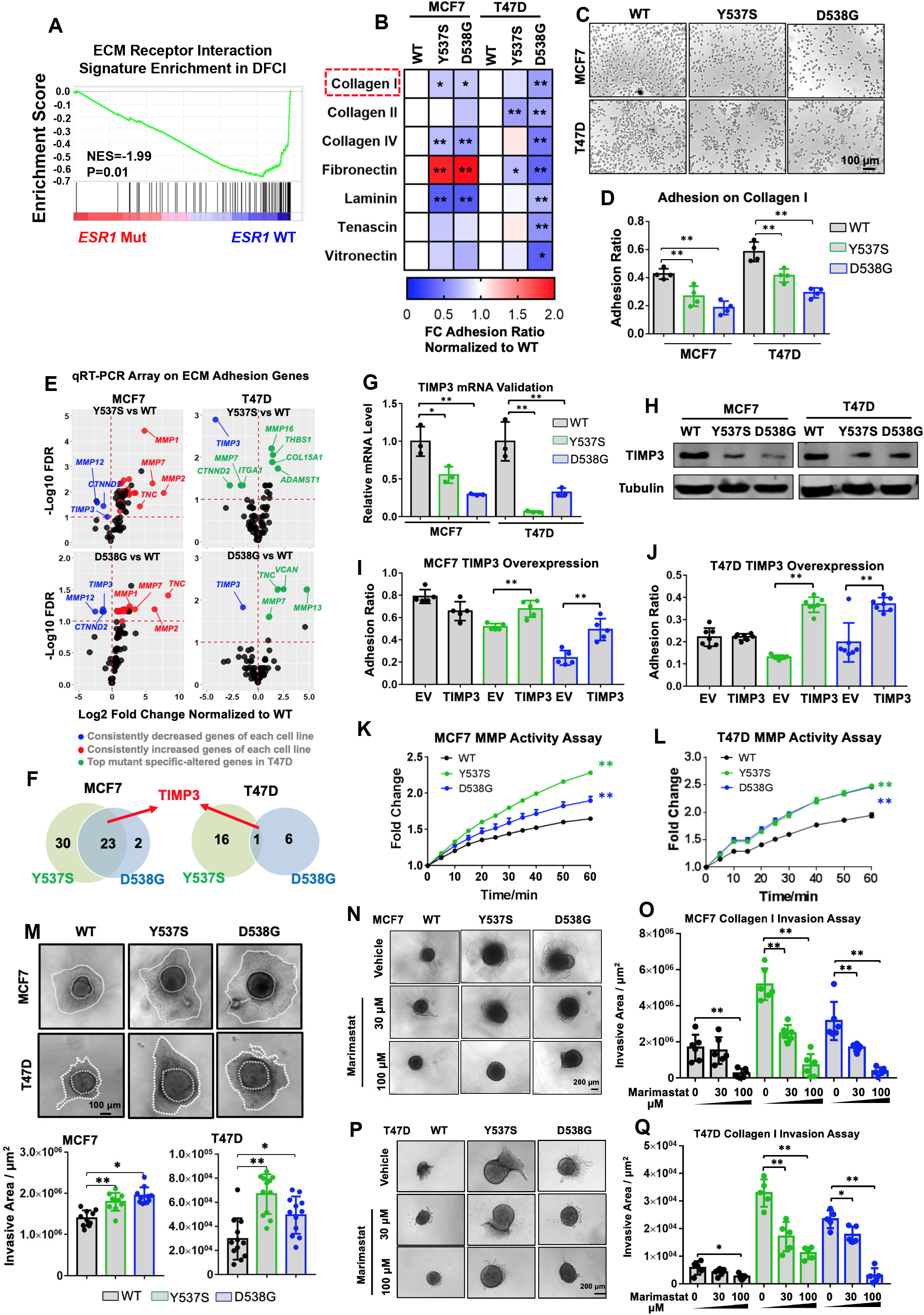
*ESR1* mutant cells show diminished ECM adhesion and enhanced invasion via an altered *TIMP3-MMP* axis. A. Gene set enrichment plots showing the comparison of enrichment levels of the “KEGG ECM Receptor Interaction” gene set (MSigDB, M7098) between WT and mutant tumors in DFCI cohort. (98 *ESR1* WT and 32 mutant tumors) B. Heatmap representation of adhesion ratio on 7 ECM components performed with MCF7 and T47D *ESR1* WT and mutant cells. Adhesion ratio of each condition with biological quadruplicates was quantified by dividing the number of remining cells after washing to the original total cells plated. All data was further normalized to WT cells within each cell line. This experiment was performed once. Dunnett’s test was applied to each condition of each cell line. (* p<0.05, **p<0.01) C. Representative images *ESR1* WT and mutant cells remaining on collagen I after three PBS washes. Images were taken using 4x magnification. Experiment displayed is representative from three independent repeats. D. Quantification of adhesion ratios on collagen I in each cell type. Bar graphs represent the mean ± SD with four biological replicates in each group. Dunnett’s test was utilized within each cell line to compare WT and mutant adhesion ratios. Experiment displayed is representative from 12 (MCF7) and 11 (T47D) independent repeats. (* p<0.05, ** p<0.01) E. Volcano plots showing the alterations of 84 ECM adhesion genes in all mutant cell types in a pairwise comparison to the WT counterparts. Genes were pre-filtered with an average Ct<35 in at least one group. An FDR<0.1 was considered as a significantly altered gene in *ESR1* mutant cells. Overlapping downregulated (blue) or upregulated (red) genes between the two mutants of each cell line were further highlighted, with gene name labels for the top targets. Top changed genes in each T47D mutant cells were labelled in green. This experiment was performed once. F. Venn diagrams showing the consistently differentially expressed genes between the two mutant variants within each cell line. *TIMP3* was highlighted as the only overlapping gene in all four *ESR1* mutant cell types. G. qRT-PCR validation of *TIMP3* expression in WT and *ESR1* mutant cells. Ct values were normalized to *RPLP0* and further normalized to WT cells. Bar graphs represent the mean ± SD with biological triplicates in each group. Representative experiment from seven independent repeats is shown. Dunnett’s test was utilized within each cell line. (* p<0.05, ** p<0.01) H. Western blot validation of *TIMP3* from whole cell lysates after hormone deprivation. Tubulin was used as a loading control. Representative experiment from six independent repeats is shown. I & J. Quantification of adhesion ratios on collagen I in each mutant variant following transfection of pcDNA empty vector or *TIMP3* plasmids in MCF7 (I) and T47D (J) cell models. Bar graphs represent the mean ± SD from 5 (MCF7) and 7 (T47D) biological replicates. Representative experiment from four independent repeats is shown. Student’s t test was used to compare the empty vector and *TIMP3* overexpressing groups. (* p<0.05, ** p<0.01) K & L. Graphical view of pan-MMP FRET kinetic assay. MMPs in MCF7 (K) and T47D (L) cell lysates were pre-activated and mixed with MMP substrates. Fluorescence was measured in a time course manner and normalized to T0 baseline and further normalized to WT cell readouts. Each point represents the mean ± SD value from three biological replicates. Representative experiment from four independent repeats is shown. Pairwise two-way ANOVA between WT and each mutant cell type was performed. (* p<0.05, ** p<0.01) M. Top panel: Representative images of the spheroid-based collagen invasion assay in *ESR1* WT and mutant cell models. MCF7 and T47D spheroids were mixed in collagen I for 4 and 6 days, respectively. Bright field images were taken accordingly with 10x magnification. Bottom panel: Quantification of invasive areas within images. Invasive areas were calculated by subtracting each original spheroid area from the corresponding endpoint total area. Each bar represents mean ± SD with 10 biological replicates. Experiments displayed are representative from three independent repeats from each cell line. Dunnett’s test was used to compare the difference between WT and mutant cells. (* p<0.05, ** p<0.01) N & P. Representative images of the spheroid-based collagen invasion assay with different doses of Marimastat treatment in MCF7 (N) and T47D (P) cell models for 4 and 6 days, respectively. Images were taken under 10x magnification. Experiments displayed are representative from three independent repeats from each cell line. Q & O. Quantification of corresponding invasive areas from 4N and 4P. Experiments displayed are representative from three independent repeats from each cell line. Student’s t test was used to compare the effects of Marimastat treatment to vehicle control. (* p<0.05, ** p<0.01)

We sought to investigate the molecular mechanisms underlying the unique defect of collagen I adhesion in *ESR1* mutant cells. There was no consistent change in expression of members of the integrin gene family, encoding well-characterized direct collagen I adhesion receptors, in our cell line models (Supplementary Fig. S14A and Supplementary Table S6). We therefore hypothesized that another gene critical in regulation of ECM genes might be altered and to test this directly, we performed gene expression analysis of 84 ECM adhesion-related genes using a qRT-PCR array (Supplementary Table S8). Pairwise comparisons between each mutant cell line and corresponding WT cells revealed a strong context-dependent pattern of ECM network reprogramming, with more pronounced effects in MCF7 cells (Fig. 4E). Intersection between Y537S and D538G mutants showed 23 and 1 consistently altered genes in MCF7 and T47D cells, respectively (Fig. 4F). *TIMP3*, the gene encoding tissue metallopeptidase inhibitor 3, was the only shared gene between all four mutant cell models (Fig. 4F), and we confirmed its decreased expression at the mRNA (Fig. 4G & Supplementary Fig. S14B) and protein level (Fig. 4H), as well as in other genome-edited *ESR1* mutant models (Supplementary Fig. S14C). E2 treatment represses *TIMP3* expression, suggesting that it’s downregulation in ESR1 mutant cells is likely due to ligand-independent repressive ER activity (Supplementary Fig. S14C). Overexpression of *TIMP3* rescued the adhesion defect in *ESR1* mutant cells (Figure 4I, 4J & Supplementary Fig. S14D), with no impact on cell proliferation (Supplementary Fig. S14E). Collectively, these data imply a selective role for *TIMP3* downregulation in causing the decreased cell-matrix adhesion phenotype of the *ESR1* mutant cells, consistent with a critical role for TIMP3 in metastasis in other cancer types (46,47).

Given the role of *TIMP3* as an essential negative regulator of matrix metalloproteinase (MMP) activity (48), we compared MMP activity between *ESR1* WT and mutant cells. A pan-MMP enzymatic activity assay revealed significantly increased MMP activation in all mutant cells (Fig. 4K & 4L), indicating that the *ESR1* mutant cells have increased capacity for matrix digestion. This was validated in spheroid-based invasion assays in which cells were embedded in collagen I (Fig. 4M) but without notable growth differences (Supplementary Fig. S15A & S15B). This was additionally visualized in co-culture spheroid invasion assays using differentially labelled T47D *ESR1* WT and mutant cells, which showed an enrichment of *ESR1* mutant cells at the leading edge of the spheroids (Supplementary Fig. S15C). Lastly, we tested if MMP blockade could repress the *ESR1* mutant-driven invasiveness. Marimastat treatment substantially reduced the invasive phenotype of *ESR1* mutant cells in a dose dependent manner (Fig. 4N–4Q). These data demonstrate that decreased TIMP3 expression, resulting in increased MMP activation causes enhanced matrix digestion associated with decreased adhesion to ECM, ultimately conferring invasive properties to *ESR1* mutant cells.

### *De novo* FOXA1-mediated Wnt pathway activation enhances of the T47D-D538G cell migration

T47D D538G cells showed increased *in vivo* tumorigenesis despite showing less pronounced adhesive phenotypes compared to T47D Y537S and MCF7 Y537S/D538G cells. Reasoning mutation and context-dependent metastatic activities of the mutant ER protein and having identified “Cellular Movement” as another top hit in our initial pathway analysis of differentially expressed genes in *ESR1* mutant tumors (Fig. 1D), we assessed potential differences in cellular migration between the different models. Wound scratch assays identified significantly increased cell motility in the T47D-D538G model (Fig. 5A & 5B), but not in T47D-Y537S (Fig. 5B) or MCF7 mutant cells (Supplementary Fig. S16A & S16B). This enhanced motility was shared between the three individual T47D-D538G clones again excluding potential clonal artifacts (Supplementary Fig. S16C & S16D). Furthermore, we observed a different morphology of T47D-D538G cells at the migratory leading edges (Fig. 5C) further confirmed by larger and stronger assembly of F-actin filaments at the edge of T47D-D538G cell clusters (Supplementary Fig. S16E-S16H). To mimic collective migration from a cluster of cells, we utilized a spheroid-based collective migration assay on type I collagen (Fig. 5D). The distance to the leading edges of T47D-D538G mutant cells was significantly longer compared to WT spheroids (Fig. 5E). In orthogonal approaches, enhanced migratory capacities of T47D-D538G cells were observed in co-culture assay using labelled T47D-WT and D538G cells (Supplementary Fig. S16I & S16J) and in Boyden chamber transwell assays (Supplementary Fig. S16K & S16L). Finally, in T47D overexpression models, we also observed significantly enhanced migration in D538G compared to WT overexpressing cells (Supplementary Fig. S17).

**Figure 5.**
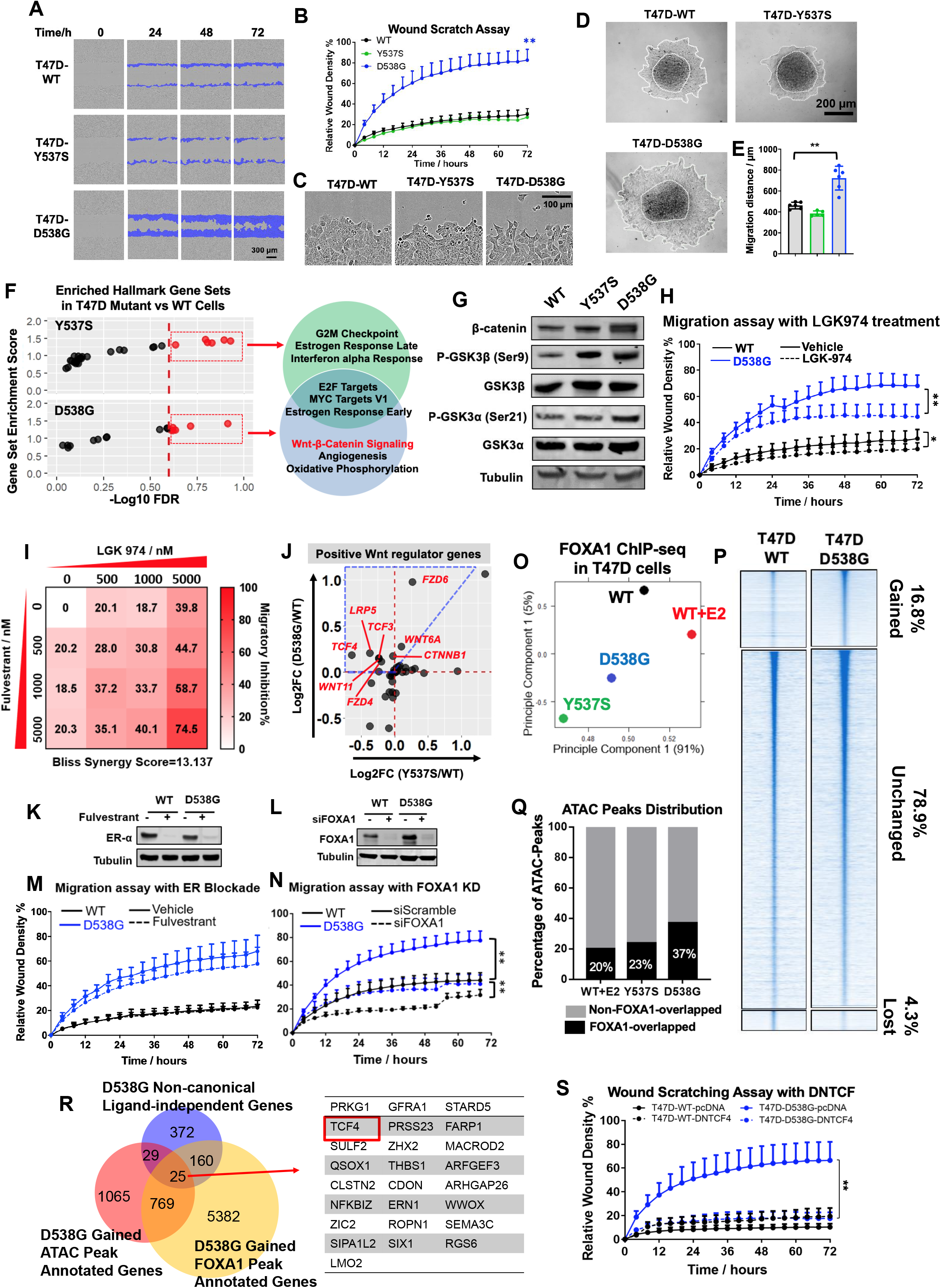
*De novo* FOXA1-mediated Wnt pathway activation enhances migratory property of the T47D-D538G cells. A & B. Representative images (A) and quantification (B) of wound scratch assay of T47D WT and *ESR1* mutant cells performed using IncuCyte living imaging system over 72 hours. The migratory region normalized to T0 are labelled in blue. Images were taken under 10x magnification. Cell migration rates were quantified based on relative wound densities with 8 biological replicates. Representative experiment from 11 independent repeats is shown. Pairwise two-way ANOVA between WT and each mutant was performed. (** p<0.01) C. Representative magnified images of the migratory edge of each group in wound scratch assays in A. D & E. Representative images (D) and quantification (E) of spheroid collective migration assays in T47D mutant cells. T47D cells were initially seeded into round bottom ULA plates to form spheroids, which were then transferred onto collagen I coated plates. Collective migration was measured after 4 days. The migratory edge of each spheroid is circled with a white line. Migratory distances were calculated based on the mean radius of each spheroid normalized to corresponding original areas. Representative experiment from three independent repeats is shown. Dunnett’s test was used for statistical analysis. (** p<0.01) F. Dot plots representing the enrichment distribution of the 50 MSigDB curated Hallmark gene sets in T47D-Y537S and T47D-D538G models normalized to WT cells.Significantly enriched gene sets (FDR<0.25) are highlighted in red, with names labeled in the venn diagram plot on the right panel. Gene sets enriched in Y537S and D538G cell models are in green and blue circles respectively. G. Immunoblot detection of ß-catenin, phospho-GSK3ß (Ser9), phospho-GSK3α (Ser21) total GSK3ß and total GSK3α levels in T47D WT and mutant cells after hormone deprivation. Tubulin was blotted as a loading control. Representative blots from three independent repeats is displayed for each protein. H. Quantification of IncuCyte wound scratch assay with or without 5μM LGK974 treatment for 72 hours. The migratory region normalized to T0 are labelled in blue. Images were taken under 10x magnification. Cell migration rates were quantified based on relative wound densities with eight biological replicates. Representative experiment from three independent repeats is shown. Pairwise two-way ANOVA between WT and each mutant was performed. (** p<0.01) I. IncuCyte migration assay with combination treatment of four different doses of LGK974 and Fulvestrant in T47D-D538G cells. Inhibition rates were calculated using the wound density at 48 hours normalized to vehicle control with values labelled using color scales in the heatmap. Positive Bliss scores are considered a synergistic combination. Representative experiment from three independent repeats is shown. J. Dot plot representing the fold changes of all Wnt signaling component genes in both T47D *ESR1* mutant cell models normalized to WT cells. The blue dotted frame highlights the unique T47D-D538G enriched genes as well as genes that are enriched in both mutants, but with a larger magnitude of enrichment in the T47D-D538G cells. K & L. Immunoblot validation of Fulvestrant-induced ER degradation (K) and FOXA1 knockdown (L). Cell lysates were subjected to ER and FOXA1 detection. Tubulin was blotted as a loading control. These validation experiments were performed once. M & N. Wound scratch assay in T47D-D538G and WT cells with 1μM of Fulvestrant treatment (M) or knockdown of FOXA1 (N) for 72 hours. Cell migration rates were quantified based on wound closure density. For fulvestrant treatment, data were merged from 3 (WT) or 6 (D538G) independent experiments. For FOXA1 knockdown, representative result from three independent repeats is displayed. Pairwise two-way ANOVA between siScramble/siFOXA1 or vehicle/Fulvestrant conditions in each cell type was performed. (* p<0.05, ** p<0.01) O. PCA plot showing the FOXA1 peak distribution of T47D WT, WT+E2, T47D-Y537S and T47D-D538G groups. P. Heatmaps representing the comparison of FOXA1 binding intensities in T47D-D538G mutants to FOXA1 binding in WT cells. Displayed in a horizontal window of ± 2kb from the peak center. The pairwise comparison between WT and mutant samples was performed to calculate the fold change (FC) of intensities. Binding sites were sub-classified into sites with increased intensity (FC>2), decreased intensity (FC<-2), and non-changed intensity (−2<FC<2). Percentages of each subgroup are labelled on the heatmaps. Q. Bar charts showing the percentage of ATAC peaks overlapping (black) or not overlapping (grey) with FOXA1 binding sites in T47D-WT, T47D-Y537S and T47D-D538G cells. R. Left panel: Venn diagram showing the intersection of genes annotated from dually gained ATAC and FOXA1 peaks (±3kb of TSS with 200kb of the peak flank) and RNA-seq differentially expressed non-canonical ligand-independent genes (gene with |fold change|>2, FDR<0.005 in D538G vs WT excluding genes with fold change|>1.5, FDR<0.01 in WT+E2 vs WT groups). Intersected genes are indicated in the right panel. S. Wound scratch assay in T47D-WT and T47D-D538G cells with or without prior transfection of a dominant negative *TCF4* plasmid for 72 hours. Pairwise two-way ANOVA between vehicle and treatment conditions was performed. Data from one representative experiment of three independent experiments (each with six biological repeats) is shown. (** p<0.01)

To understand the mechanisms underlying the migratory phenotype of T47D-D538G cells we identified pathways uniquely enriched in these cells. GSEA identified endocrine resistance-promoting pathways (e.g. E2F targets) in both T47D mutants, whereas Wnt-β-catenin signaling was one of the uniquely enriched pathways in T47D-D538G (Fig. 5F). Hyperactivation of the canonical Wnt-ß-catenin pathway was further confirmed by a Top-Flash luciferase assay (Supplementary Fig. S18A). We also observed increased phosphorylation of GSK3β and GSK3α as well as β-catenin (both total and nuclear) protein levels in T47D-D538G cells (Fig. 5G and Supplementary Fig. S18B). To address the potential clinical relevance of these findings, we utilized the porcupine inhibitor LGK974, which prevents the secretion of Wnt ligands and is currently being tested in a clinical trial for patients with advanced solid tumors including breast cancer (NCT01351103) (49,50). Treatment with LGK974 resulted in a 20% and 40% inhibition of T47D *ESR1* WT and D538G mutant cell migration respectively (Fig. 5H and Supplementary Fig. S18C) yet had no effect on cell proliferation (Supplementary Fig. S18D). We next studied the combination of LGK974 and the selective ER degrader (SERD), Fulvestrant, in migration assays, in which we detected significant synergy (Fig. 5I), suggesting that combination therapy co-targeting the Wnt and ER signaling pathways might reduce the metastatic phenotypes of Wnt hyperactive *ESR1* mutant tumors.

We sought to decipher the mechanisms underlying T47D-D538G Wnt hyperactivation. Comparing the fold changes of canonical Wnt signaling positive regulators between T47D-Y537S and T47D-D538G mutant cells, we identified eight candidate genes exhibiting pronounced enrichment in T47D-D538G cells (Fig. 5J), including ligands (e.g. *WNT6A*), receptors (e.g. *LRP5*) and transcriptional factors (e.g.*TCF4*). With the exception of *LRP5*, none of these candidate genes were induced by E2 stimulation in T47D *ESR1* WT cells (Supplementary Fig. S19A). Lack of consistent E2 regulation was confirmed in five additional ER+ breast cancer cell lines (Supplementary Fig. S19B). Hence, we alternatively hypothesized that D538G ER might gain *de novo* binding sites proximal to Wnt pathway genes allowing their induction. We mapped ER binding globally by analyzing ER ChIP-sequencing in T47D WT and *ESR1* mutant cells. Consistent with previous studies (14,28), mutant ER were recruited to binding sites irrespective of hormone stimulation (Supplementary Fig. S19C & Table S9). However, none of the mutant ER bound regions mapped to identified Wnt pathway genes (± 50kb of TSS), again suggesting a lack of direct canonical ER regulation. Moreover, short-term fulvestrant treatment only weakly dampened T47D-D538G cell migration (Fig. 5K & 5M) suggesting that ER activation may not be an essential prerequisite for enhanced cell migration in D538G cells.

Given our recent findings of enriched FOXA1 motifs in gained open chromatin of T47D-D538G cells (19), we decided to validate this pivotal *in silico* prediction, focusing on our observed migratory phenotype. In contrast to the limited effects of ER depletion, strikingly, FOXA1 knockdown fully rescued the enhanced migration in T47D-D538G cells (Fig. 5L & 5N), indicating a more dominant role of FOXA1 in controlling T47D-D538G cell migration. Ligand-independent 2D growth of T47D-D538G cells was inhibited by both fulvestrant and FOXA1 knockdown (Supplementary Fig. S19D), suggesting a canonical ER-FOXA1 co-regulatory mechanism in growth, distinguished from the role of FOXA1 in the regulation of migration.

To further explore how FOXA1 contributes to the migratory phenotype, we performed FOXA1 ChIP-sequencing to decipher the genomic binding profiles. We identified approximately 30,000 peaks in T47D WT cells regardless of E2 stimulation and a ~1.6 fold increase in binding sites of the Y537S (61,934) and D538G (54,766) ER mutants (Supplementary Fig. S20A & Supplementary Table S9). PCA distinctly segregated all four groups (Fig. 5O), suggesting unique FOXA1 binding site redistribution. Comparison of binding intensities revealed 14%, 28% and 21% FOXA1 binding sites were altered in WT+E2, Y537S and D538G groups, respectively, with a predominant gain of binding intensities in the two T47D mutants (Fig. 5P and Supplementary Fig. S20B).

Since FOXA1 is a well-known essential pioneer factor of ER in breast cancer, we examined interplay between FOXA1 and WT and mutant ER. Interestingly, both Y537S (39%) and D538G (25%) ER binding sites showed a significantly lower overlap between FOXA1 compared to the WT+E2 group (56%), albeit with the increased number of gained mutant FOXA1 binding sites (Supplementary Fig. S20C). This discrepancy suggests that FOXA1 exhibits a diminished ER pioneering function and instead might contribute to novel functions via gained *de novo* binding sites. Co-occupancy analysis using isogenic ATAC-seq data (19) uncovered that the open chromatin of T47D-D538G cells was more associated with FOXA1 binding sites compared to WT and T47D-Y537S cells (Fig. 5Q). FOXA1 binding intensities were also stronger in D538G ATAC-sites (Supplementary Fig. S20D). Collectively, these results provide evidence that FOXA1 likely plays a critical role in the D538G mutant cell to reshape its accessible genomic landscape.

We further investigated the impact of the gained FOXA1-associated open chromatin on transcriptomes, particularly exploring *ESR1* mutant-specific genes. Intersection of the gained FOXA1- and ATAC-sites for annotated T47D-D538G genes with non-canonical ligand-independence identified 25 potential targets that could be attributed to *de novo* FOXA1 bound open chromatin, exemplified by *PRKG1* and *GRFA* as top targets (Fig. 5R & Supplementary Fig. S21A). Notably, one of our identified D538G specific Wnt regulator genes, *TCF4*, was uncovered in this analysis. Higher *TCF4* expression in T47D-D538G cells was validated by qRT-PCR and furthermore this increased expression could be fully blocked following FOXA1 knockdown (Supplementary Fig. S21B). Additionally, stronger FOXA1 recruitment at the *TCF4* gene locus was validated via ChIP-qPCR (Supplementary Fig. S21C and S21D). Importantly, overexpression of dominant negative *TCF4* strongly impaired cell migration in T47D-D538G, while it only slightly affected WT cells (Fig. 5S). Together, these results support that FOXA1 binding site redistribution leads to novel chromatin remodeling and enhanced expression of genes with roles in metastases including *TCF4*, which subsequently activate Wnt-driven migration in T47D-D538G cells.

## Discussion

Hotspot somatic mutations clustered in the LBD of ER represent a prevalent molecular mechanism that drives antiestrogen resistance in ~30% of advanced ER+ breast cancer. There is an urgent need for a deeper understanding of this resistance mechanism in order to develop novel and personalized therapeutics. Utilizing clinical samples, *in silico* analysis of large datasets, and robust and reproducible experimentation in multiple genome-edited cell line models, our study uncovers complex and context-dependent mechanisms of how *ESR1* mutations confer gain-of-function metastatic properties. We identified *ESR1* mutations as multimodal metastatic drivers hijacking adhesive and migratory networks, and thus likely influencing metastatic pathogenesis and progression. Mechanistically, we uncovered novel ER-indirect regulation of metastatic candidate gene expression, distinct from previously described (11,12,51) canonical ligand-independent gene induction (Fig. 6). Nonetheless, some limitations were noted in our study, such as the lack of *in vivo* validation of studied therapeutic approaches. In addition, our numbers for clinical samples of paired primary-metastatic tumors harboring *ESR1* mutations is finite, necessitating validation in future studies with larger clinical cohorts.

**Figure 6.**
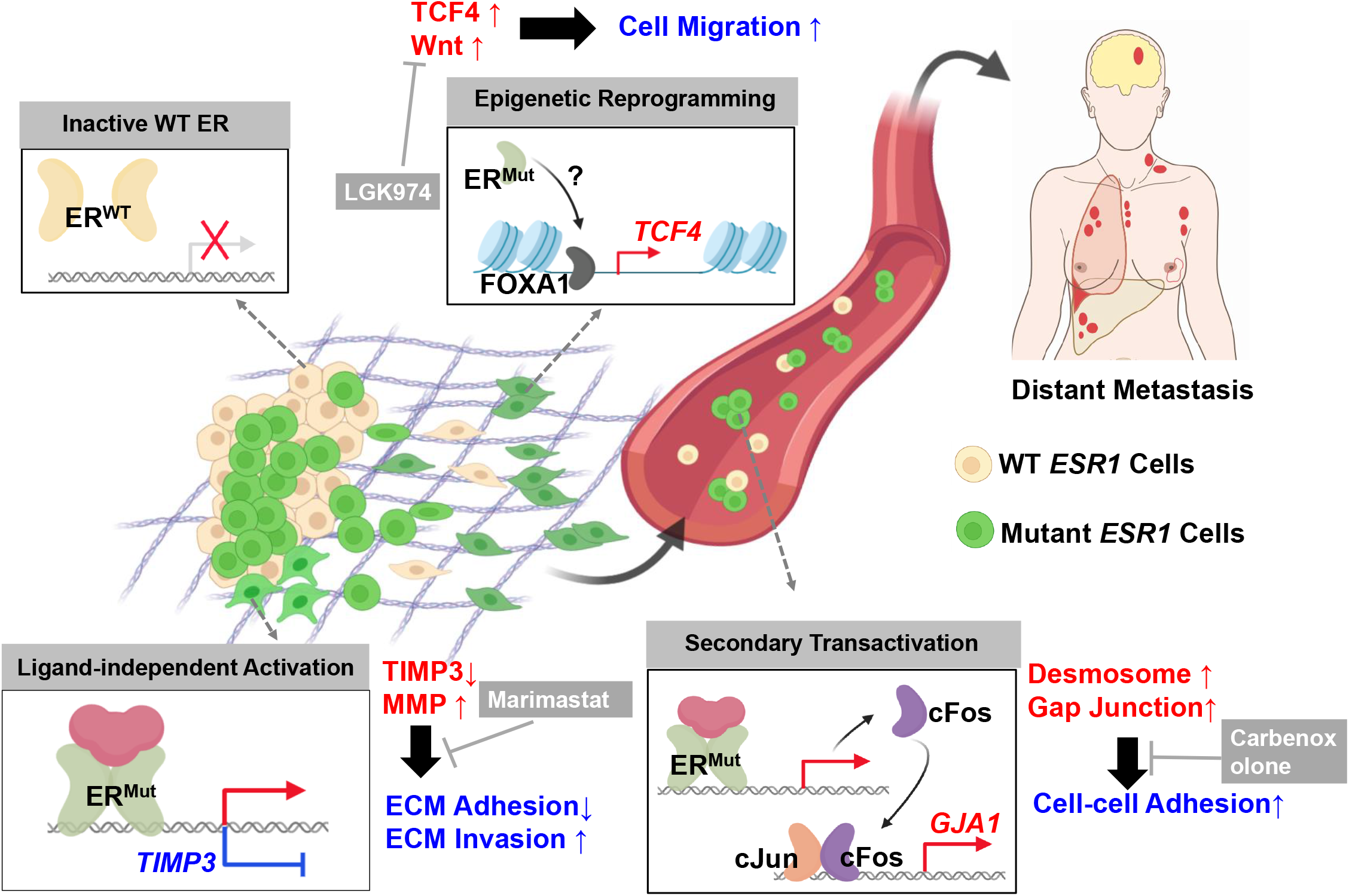
Schematic model of *ESR1* mutation-driven breast cancer metastases. Mutated ER triggers differential gene regulatory reprogramming through 1) ligandin-dependent transcriptional gene activation or repression, 2) secondary transcriptional regulation and 3) FOXA1-driven epigenetic remodeling. Ligand independent transcription constitutively induces or represses canonical ER regulated sites (e.g. *TIMP3*). Secondary transactivation induces gene expression indirectly via activation of an intermediate regulator (e.g. *GJA1*). Novel epigenetic remodeling includes *de novo* FOXA1 redistribution and increased chromatin accessibility at specific gene loci (e.g. *TCF4*). Consequently, increased desmosome and gap junction expression, *TIMP3*-MMP axis alteration and hyperactivation of the Wnt pathway results in enhanced cell-cell adhesion, collagen invasion, migration and decreased cell-ECM adhesion, ultimately facilitating metastases of *ESR1* mutant cells. Corresponding therapeutic vulnerabilities can be efficiently targeted by carbenoxolone, marimatsat and LGK974. These mechanisms are highly context dependent with phenotypes labeled for specific cell line models.

We discovered enhanced cell-cell adhesion via upregulated desmosome and gap junction networks in cell lines and clinical samples with *ESR1* mutations. These transcriptional alterations are associated with a specific clinical phenotype characterized not only by treatment resistance, but also by high CTC count and a different metastatic organotropism (52,53). We propose that this key alteration may support increased metastases in ER mutant tumors through facilitating the formation of homo- or heterotypic CTC clusters, providing a favorable environment for CTC dissemination, as previously described (30). This idea is further supported by previous data showing upregulation of the desmosome gene plakoglobin (*JUP*) and cytokeratin 14 (*KRT14*), which may play a role in a CTC cluster formation signature (30,54). We observed increased expression of plakophilin, desmocollin, and desmoglein in *ESR1* mutant cells, suggesting the importance of the broad desmosome network reprogramming for functional cell clustering activity. Moreover, enhanced gap junction genes might potentiate intercellular calcium signaling, facilitating the prolonged survival of various metastatic cell types tethered to *ESR1* mutant cells *en route* (55). Dissociation of CTC cluster using Na+/K+ ATPase inhibitors decreased metastasis *in vivo* (37). In addition, previous studies have validated the anti-tumor effects of FDA-approved gap junction blockers carbenoxolone and mefloquine *in vivo* (56,57). Our results warrant additional preclinical studies using drugs targeting desmosome and gap junctions, with the ultimate goal of applying these treatments in a CTC-targeted clinical trial to improve outcomes for patients harboring breast cancers with *ESR1* mutations.

Previous studies using similar *ESR1* mutant cell models described enhanced migratory properties (15,16), but no mechanistic explanations were uncovered. Here we identify a critical role for Wnt-ß-catenin signaling and show that co-targeting of Wnt and ER resulted in synergistic inhibition of cell migration. Intriguingly, the strong effect we observed on migration was unique to T47D-D538G cells, a discovery that was made possible through our use of multiple genome-edited mutation models. This finding might help explain the higher frequency of D538G mutations in metastatic samples, despite the stronger endocrine resistance phenotype of Y537S mutation (5,12,14,29,58). Of note, slightly higher Wnt activity and β-catenin accumulation were also observed in T47D-Y537S cells, but this failed to convert into a migratory phenotype. It is possible that some genes uniquely regulated by Y537S ER in T47D cells might inhibit migratory phenotypes. For instance, the gap junction component, connexin 43, which is exclusively upregulated in T47D-Y537S cells, has been reported to play an inhibitory role in epithelial cell migration (59). *In vivo* experiments revealed striking enhancement of metastatic capacity in the MCF7-Y537S but not D538G model. This discrepancy with *in vitro* data could possibly be explained by the longer distant metastatic latency requirement of D538G cells *in vivo*, consistent with a recent study using overexpression cell models (14). These data support strong allele and context dependent effects of the *ESR1* mutation on metastatic phenotypes, in line with context dependent effects on transcriptome, cistromes and accessible genome in *ESR1* mutant cells (11,12,14,19). Of note, previous efforts using multiple cell line models with *ESR1* mutations elucidated several congruent molecular and functional alterations associated with endocrine resistance (14,15,51,58), suggesting that mechanisms underlying metastasis of *ESR1* mutant clones exhibit a higher degree of heterogeneity. This is also supported by clinical data: the recent BOLERO2 trial showed significant differences in overall survival and everolimus response between Y537S and D538G mutations (9), and results from the recent PALOMA3 trial suggest a potential Palbociclib resistance uniquely gained in tumors bearing the Y537S mutation (60). Taken together, these proof-of-concept studies are setting the stage for a more contextual and personalized therapeutic targeting strategy in *ESR1* mutant breast cancer.

Of note, our comprehensive clinical investigation from four different cohorts (>900 samples) suggest that *ESR1* mutations are uncommon in local recurrences. The significant exclusion of *ESR1* mutations in local recurrences is likely due to that *ESR1* mutant clones are more equipped to escape from local-regional microenvironment. A recently published study identified hotspot *ESR1* mutations in 15 out of 41 (36%) of local-regional ER+ recurrences albeit at significantly lower mutation allele frequencies (61). The reasons for this discrepancy are not clear, and future efforts are warranted to explore details of potential differences in clinic-pathological features of the cohorts, and technical approaches.

Lastly, we also sought to address the ER regulatory mechanisms involved in induction of candidate metastatic driver genes utilizing ChIP-seq technology. Interestingly, none of the metastatic candidate genes in *ESR1* mutant cells gained proximal ER binding sites. This could be a result of our stringent hormone deprivation protocol resulting in depletion of weaker binding events, and thus less sensitive binding site readouts (62). This idea is supported by ChIP-seq data from Harrod et al. (28), which shows stronger ER binding sites around *DSC2, DSG2* and *TIMP3* gene loci in MCF7-Y537S cells. Our data, however, clearly shows that ER mutant cells display changes in indirect gene regulation, resulting in metastatic phenotypes. This observation is due to non-canonical ER action on chromatin structure remodeling, which was alternatively validated from our ATAC-seq and FOXA1 ChIP-seq data. We propose that mutant ER reprograms FOXA1, resulting in redistribution of FOXA1 binding to specific enhancers controlling the key migratory driver gene(s). In addition, several recent studies uncovered the promising role of androgen receptor (AR) in *ESR1* mutant tumors and cell models (18,63,64), and additional studies are warranted to study *de novo* interplay between FOXA1, AR and mutant ER.

Overall, our study serves as a timely and important preclinical report uncovering mechanistic insights into *ESR1* mutations that can pave the way towards personalized treatment of patients with advanced metastatic breast cancer.

## Acknowledgement

We are grateful for advice, discussions and technical support from Dr. Ye Qin, Dr. Yu Jiang, Dr. Min Yu, Yonatan Amzaleg and Meghan S. Mooring. We would like to thank Dr. Peilu Wang for her contribution to earlier studies in the Lee-Oesterreich group on *ESR1* mutations. This project used the University of Pittsburgh HSCRF Genomics Research Core, the University of Pittsburgh Center for Research Computing, and the UPMC Hillman Cancer Center Tissue and Research Pathology Services supported in part by NIH grant award P30CA047904. The authors would like to thank the patients who contributed samples to the tissue bank as well as all the clinicians and staff for their efforts in collecting tissues.

## Materials and methods

Additional details are provided in the Supplementary Materials and Methods section.

### Human tissue studies from the Womens Cancer Research Center (WCRC) and Charite cohorts

All patients enrolled were approved within IRB protocols (PRO15050502) from the University of Pittsburgh and Charite Universitaetsmedizin Berlin. Informed consent was obtained from all participating patients. Biopsies were obtained and divided into distant metastatic or local recurrent tumors. Genomic DNA was isolated from formalin fixed paraffin embedded (FFPE) samples and *ESR1* mutation status was detected with droplet digital PCR (ddPCR) targeting Y537S/C/N and D538G mutations in preamplified *ESR1* LBD products as previously reported (7).

For the 54 ER+ metastatic tumor samples, genomic profiles were determined based on tumor RNA sequencing provided in previous publications (25,26,65).

### CTCs analysis from the NU16B06 Cohort

A retrospective cohort comprising 151 Metastatic Breast Cancer (BC) patients characterized for CTCs, and ctDNA at the Robert H. Lurie Comprehensive Cancer Center of Northwestern University (Chicago, IL) between 2015 and 2019 was analyzed. Patients’ enrollment was performed under the Investigator Initiated Trial (IIT) NU16B06 independently from treatment line. The overall baseline staging was performed according to the investigators’ choice, CTCs and ctDNA collection was performed prior to treatment start. CTC enumeration was performed though the CellSearch™ immunomagnetic System (Menarini Silicon Biosystems). Mutations in *ESR1* (hotspots D538 and Y537) and PIK3CA (hotspots E453 and H1047) were detected by either ddPCR assay using the QX200 ddPCR System (Bio-Rad) or through the Guardant360™ high sensitivity next-generation sequencing platform (Guardant Health, CA). More details for CTC enumeration, mutation detection and statistical analysis can be found in Supplementary Materials and Methods.

### Cell culture

Genome-edited MCF7 and T47D *ESR1* mutant cell models from different sources were maintained as previously described (12,19,28). Hormone deprivation was performed for all experiments, unless otherwise stated. Other parental cell lines, ZR75-1 (CRL-1500), MDA-MB-134-VI (HTB-23), MDA-MB-330 (HTB-127) and MDA-MB-468 (HTB-132), were obtained from ATCC. BCK4 cells were developed as previously reported (66).

### Reagents

17β-estradiol (E2, #E8875) was obtained from Sigma, and Fulvestrant (#1047), carbenoxolone disodium (#3096) and EDTA (#2811) were purchased from Tocris. LGK974 (#14072) and T-5224 (#22904) were purchased from Cayman. Marimastat (S7156) was obtained from SelleckChem. Recombinant human Wnt3A (5036-WN-010) was purchased from R&D Systems. For knockdown experiments, siRNA against *FOXA1* (#M-010319), *DSC1* (#L-011995), *DSC2* (#L-011996), *GJA1* (#L-011042) and *GJB2* (#L-019285) were obtained from Horizon Discovery. Desmosome and scramble peptides were designed based on previous studies (67,68) and synthesized from GeneScript. Peptide sequences are presented in Supplementary Table S10.

### Animal Studies

#### Long term metastatic evaluation

4-week old female *nu/nu* athymic mice were ordered from The Jackson Laboratory (002019 NU/J) according to University of Pittsburgh IACUC approved protocol #19095822. MCF7 and T47D *ESR1* mutant cells were hormone deprived and resuspended in PBS with a final concentration of 10^7^ cells/ml. 100μl of cell suspension was then injected via tail vein into nude mice with 7 mice per group. Mice were under observation weekly. According to the IACUC protocol, if greater than 50% of mice in any group show predefined signs of euthanasia, the entire cohort needs to be euthanized. Cohorts were euthanized at 13 weeks for MCF7 cell-injected mice and 23 weeks for T47D cell-injected mice. Macro-metastatic tumors and potential organs (lung, liver, UG tract) for metastatic spread were harvested. Solid macro-metastatic tumors (non-lymph node) were counted for comparison. All tissues were processed for FFPE preparation and hematoxylin and eosin (H&E) staining by the Histology Core at Magee Women’s Research Institute. Macro-metastatic tumor FFPE sections were further evaluated by a trained pathologist. Micro-metastatic lesions in the lung were further examined and quantified by immunofluorescence staining as described in supplementary materials and methods.

#### Short term CTC cluster assessment

4-week old female *nu/nu* athymic mice were ordered from The Jackson Laboratory (002019 NU/J) according to University of Pittsburgh IACUC approved protocol #19095822. MCF7 WT and mutant cells were stably labelled with RFP-luciferase by infection with the pLEX-TRC210/L2N-TurboRFP-c lentivirus plasmid. Labelled cells were hormone deprived and resuspended in PBS at a final concentration of 10^7^ cells/ml. 100μl of cell suspension was then injected into nude mice with 6 mice per group via an intracardiac left ventricle injection. Post-injected mice were immediately imaged using the IVIS200 *in vivo* imaging system (124262, PerkinElmer) after D-luciferin intraperitoneal injection to confirm successful cell delivery into the circulation system. All mice were euthanized after one hour of injection and their whole blood were extracted via cardiac puncture and collected into CellSave Preservative Tubes (#790005, CellSearch). Blood samples were mixed with 7ml of RPMI media and shipped to University of Minnesota for CTC enrichment. CTCs were extracted using an electric size-based microfilter system (FaCTChekr) and stained with antibody against pan-cytokeratins (CK) and DAPI. Slides with stained CTCs were manually scanned in a blind manner and all visible single CTCs or clusters were imaged under 5X or 40X magnification respectively. To set up criteria for identifying CTC clusters via images, we analyzed seven single CTCs with intact CK signal distribution and calculated the average nuclei-edge to membrane distance (x). Inter-nuclei-edge distance greater than 2x for any two CTCs were excluded in CTC cluster calling. All measurements were performed in a blind manner. Details of filter and staining are included in the supplementary materials and methods.

### qRT-PCR

MCF7 and T47D cells were seeded in triplicates into 6-well plates with 120,000 and 90,000 cells per well respectively. After desired treatments, RNA was and cDNA was synthesized using iScript kit (#1708890, BioRad, Hercules, CA). qRT-PCR reactions were performed with SybrGreen Supermix (#1726275, BioRad), and the ΔΔCt method was used to analyze relative mRNA fold changes with *RPLP0* measurements serving as the internal control. All primer sequences can be found in Supplementary Table S10.

### Immunoblotting

After desired treatments, cells were lysed with RIPA buffer spiked with a fresh protease and phosphatase cocktail (Thermo Scientific, #78442) and sonicated. Protein concentrations were quantified using the Pierce BCA assay kit (Thermo Fisher, #23225). 80-120μg of protein for each sample was loaded onto SDS-PAGE gels, and then transferred onto PVDF membranes. The blots were incubated with the following antibodies: desmocollin 1 (sc-398590), desmoglein 2 (sc-80663), plakophilin (sc-33636), connexin 26 (sc-7261) and cFOS (sc-52) from Santa Cruz; ER-α (#8644), HA (#3724), Non-phospho-ß-catenin (#19807), Histone H3 (#4499), AIF (#5318), GSK3β (Ser9, #5558), phospho-GSK3α (Ser21, #9316), GSK3β (#12456) and GSK3α (#4337) from Cell Signaling Technology; β-catenin (#610154) from BD; Tubulin (T6557) and connexin 43 (C6219) from Sigma Aldrich; and *TIMP3* (ab39184) from Abcam.

### IncuCyte Live Cell Imaging System

#### Wound scratch assay

MCF7 or T47D cells were seeded at 150,000 cells/well into Imagelock 96-well plates (Essen Bioscience, #4379) pre-coated with Matrigel (Corning, #356237). Wounds were scratched in the middle of each well using a Wound Maker (Essen Bioscience, #4493). Desired treatments mixed with 5μg/ml of proliferation blocker Mitomycin C (Sigma-Aldrich, #10107409001) were loaded after two washes with PBS. The IncuCyte Zoom system was used to record wound images every 4 hours and wound closure density was calculated using the manufacturer’s wound scratch assay module. For the dominant negative *TCF4* overexpression experiment, Myc-tagged DN*TCF4* plasmids (Addgene, #32729) were transiently transfected into targeted cells for a total of 24 hours before being subjected to the wound scratch assay.

#### Aggregation rate assay

3,000 MCF7 or 4,000 T47D cells were seeded into 96-well round bottom ultra-low attachment plates (Corning, #7007) with 100μl of respective media in each well. Cell aggregation was monitored by the IncuCyte living imaging system every hour. Spheroid areas were normalized to time 0.

### Calcein-labelled cell-cell interaction assay

MCF7 and T47D cells were seeded into black-walled 96 well plate at 15,000 cells per well to achieve a fully confluent monolayer after 24 hours. Separate cultures of cells were digested and labelled with 1μM calcein AM (BD Pharmingen, #564061) for 30 minutes in room temperature. 40,000 labelled cells were loaded on top of the previously plated monolayers and incubated for 1 hour at 37°^C^. Cells were washed three times after incubation by manually pouring out the PBS washing agent. The plates were read using Victor X4 plate reader (PerkinElmer) under the excitation and emission wavelength of 485/535nm. Cell-cell adhesion ratios were calculated by dividing the post-wash readouts to the pre-wash readouts after each wash. For the vacuum aspiration method, we used a standard laboratory vacuum pump with a modified speed of approximately 100 ml/minutes. Adhesion ratios after three washes were plotted separately for each independent experiment.

### Ibidi microfluidic system

MCF7 and T47D *ESR1* mutant cells were hormone deprived for 3 days and diluted to 10^6^ cells in 14ml of respective media before being loaded into the ibidi pump system (ibidi, #10902). Cells were constantly flowing with 15dynes/cm of shear stress for two hours before immediate imaging after being seeded back into a flat bottom ULA plate. For each group, six wells were imaged twice. Time zero (T0) cells were also imaged as the initial time point control. Cell numbers in clusters or non-clusters were manually counted. Cell cluster ratios were calculated by dividing the cell numbers in clusters to the total number of cells. Cell clustering grade was calculated by the cell numbers present in each cluster. For CBX treatment, cells were pre-treated with 100μM CBX for two days before being added to the flow chamber. For the desmosome blocking peptides treatment, 75μM of each *DSC1, DSC2, DSG1* and *DSG2* peptide or 150μM of each scramble peptide were pre-mixed into cell suspension for flow experiments.

### Cell-ECM adhesion assay

30,000 cells/well were seeded into collagen I coated (Thermo Fisher Scientific, A1142803) or uncoated 96-well plates. For the ECM array assay, cells were resuspended and loaded into the ECM array plate (EMD Millipore, ECM540). After a 2-hour incubation at 37°^C^, the plates were washed with PBS three times, and attached cells were quantified using the FluoReporter kit (Thermo Fisher Scientific, F2962). Adhesion ratios were calculated by dividing the remaining cell counts in the washed wells to the initial cell counts in pre-washed plates. For *TIMP3* overexpression, the PRK5M-*TIMP3* plasmid (Addgene, #31715) was transfected into targeted cells, which was subjected to the adhesion assay after a 24-hour transfection period.

### Chromatin-immunoprecipitation (ChIP)

ChIP experimentation was performed as previously described (39). The immunoprecipitation was performed using ERα (sc543) and rabbit IgG (sc2027) antibodies (Santa Cruz Biotechnologies). Histone 3 acetylation at K27 site (ab4739), and Histone 3 di-methylation at K4 site (ab7766) and FOXA1 (ab23738) antibodies were obtained from Abcam.

### ChIP-sequencing Analysis

ChIP-seq reads were aligned to Hg19 reference genome assembly using Bowtie 2.0 (69), and peaks were called using MACS2.0 with a p-value<10^-5^ (ER ChIP-seq) or a q-value<0.05 (FOXA1 ChIP-seq) (70). We used the Diffbind package (71) to perform principle component analysis, identify differentially bound regions and analyze intersection ratios with other datasets. Briefly, all BED files for each cell line were merged and binding intensity was estimated at each site based on the normalized read counts in the BAM files. Pairwise comparisons between WT and mutant samples were performed to calculate fold change (FC). Binding sites were sub-classified into three categories: gained sites (FC>2), lost sites (FC<-2), and not-changed sites (2<FC<2). Heatmaps and intensity plots for binding peaks were visualized by EaSeq (72). For gene annotation from FOXA1 binding sites, gained FOXA1 peaks were selected and annotated genes were inspected in a ± 200kb range of the FOXA1 peaks using ChIPseeker (73). For intersection analysis of the D538G-regulated non-canonical ligand-independent genes, broad differentially expressing genes were first called using a cutoff of |fold change|>2, FDR<0.005 between WT and D538G cells. Meanwhile, E2-regulated genes were called using the cutoff of |fold change|>1.5, FDR<0.01 between WT and WT+E2 groups. D538G ligand-independent genes which are also regulated in WT cells were excluded from the analysis.

### RNA-sequencing analysis

Data generation and processing of the 54 ER+ tumors in the WCRC cohort was described above. For the MET500 cohort, RNA-seq fastq files from 91 metastatic breast cancer samples were downloaded from the Database of Genotypes and Phenotypes (dbGaP) with accession number phs000673.v2.p1. Transcript counts from all samples were quantified with Salmon v.0.8.2 and converted to gene-level counts with tximport. The gene-level counts from all studies were then normalized together using TMM with edgeR. Log2 transformed TMM-normalized counts per million [log_2_(TMM-CPM+1)] were used for analysis. 46 putative ER positive samples were then filtered in the MET500 cohort. *ESR1* mutation status was extracted using the MET500 portal (https://met500.path.med.umich.edu). For the DFCI cohort, raw counts data was obtained and normalized to log_2_(TMM-CPM+1) for further analysis. *ESR1* mutation status was called using separate whole exon sequencing data. For the POG570 cohort, raw count matrixes and mutation statuses were downloaded from the BCGSC portal (https://www.bcgsc.ca/downloads/POG570/). ER status of each patient was additionally requested from the cited original resources and only ER+ metastatic tumors were used for downstream analysis.

For all datasets, differential expression (DE) analysis was performed using the DESeq2 package (74). In brief, genes were prefiltered with a log_2_(CPM+1)>1 in at least one sample criteria across all data sets. DE genes with a q-value below 0.1 and an absolute log_2_ fold change above 1.5 were used for Ingenuity Pathway Analysis (75). GSEA analysis was performed using the Broad GSEA Application (76). Gene set variation analyses were performed using the GSVA package (77). All gene sets used in this study are reported in Supplementary Table S6. Data visualizations were performed using “ggpubr” (78) and “VennDiagram” packages (79).

### Statistical Analysis

GraphPad Prism software version 7 and R version 3.6.1 were used for statistical analysis. All experimental results included biological replicates and were shown as mean ± standard deviation, unless otherwise stated. Specific statistical tests were indicated in corresponding figure legends. All tests were conducted as two-tailed, with a p<0.05 considered statistically significant. Drug synergy was calculated based on the Bliss independence model using the SynergyFinder (https://synergyfinder.fimm.fi/) (80). Bliss synergy scores were used to determine synergistic effects.

### Data Availability

The ER and FOXA1 ChIP-seq data has been deposited onto the Gene Expression Omnibus database (GSE125117 and GSE165280). All publicly available resources used in this study are summarized in Supplementary Table S11. All raw data and scripts are available upon request from the corresponding author.

