## Supplementary material for "Hotspot *ESR1* mutations are multimodal and contextual drivers of breast cancer metastasis": Suppl. Figure_ESR1_Mut_Metastasis_Final.pdf

Supplementary Figure

### Supplementary Figure S1

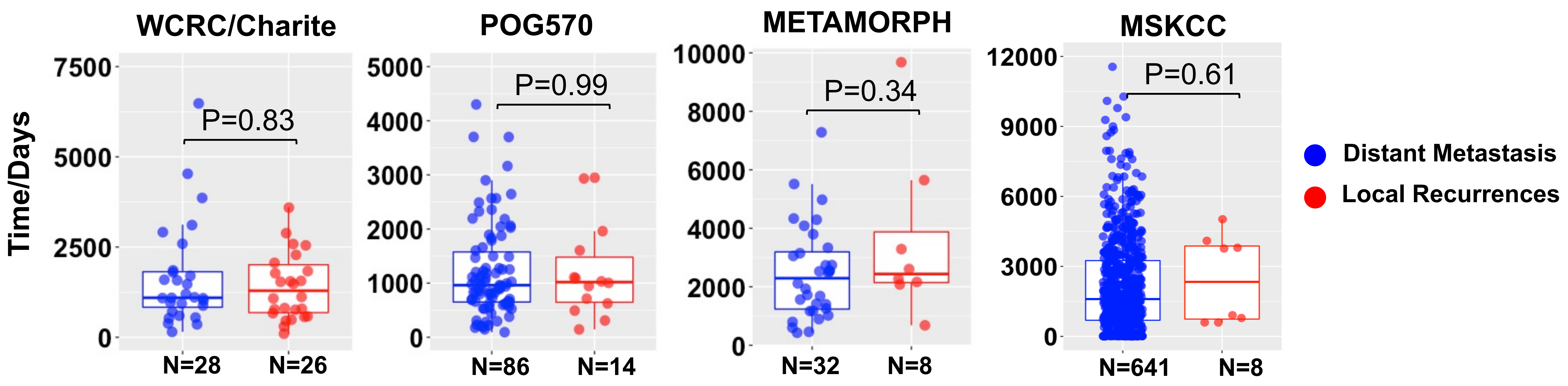


**Figure S1. There is no difference in time to recurrence between local and distant recurrent samples used in this study. (Related to Table 1)**
Comparison of time to recurrence between distant metastatic and local recurrent samples in all four cohorts. Patients with time to recurrence=0 were excluded in this analysis. Mann-Whitney U test was performed to compare the RFS between local recurrent and distant metastatic samples. For WCRC and POG570, recurrence free survival was used. For the METAMORPH cohort, duration between primary tumor resection and metastatic tumor biopsy was used in this analysis. For MSKCC, time to recurrence was calculated as days between diagnosis of the primary tumor and recurrence. Specific number for each group in each cohort is labelled.

### Supplementary Figure S2

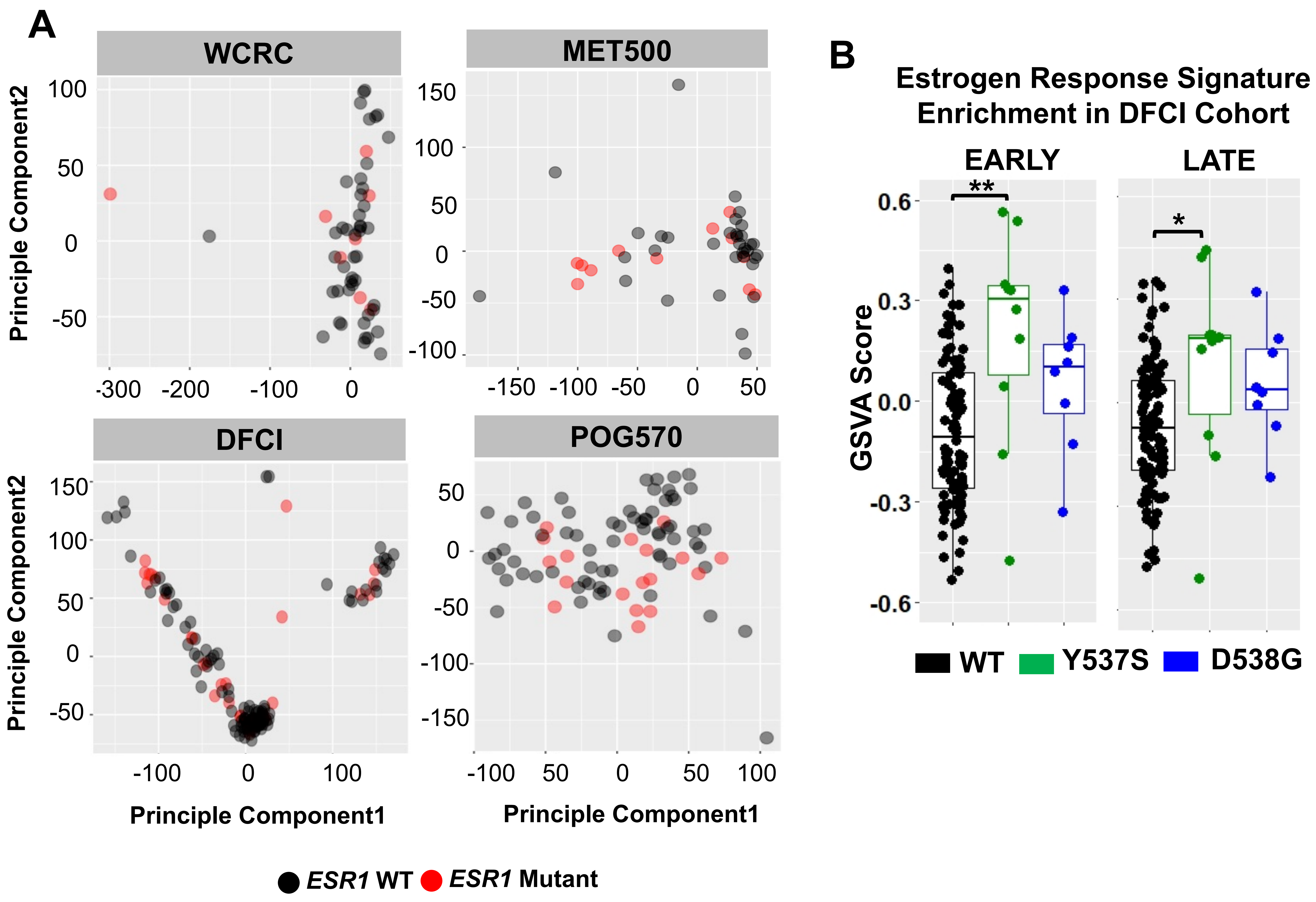


**Figure S2. *ESR1* mutant and WT tumors do not segregate and estrogen receptor activation is different between predominant mutant variants. (Related to Fig. 1)**

A. Principle component analysis of the global transcriptomes from all four ER+ metastatic tumor cohorts. (WCRC, 46 *ESR1* WT/8 mutant; MET500, 34 *ESR1* WT/12 *ESR1* mutant; DFCI, 98 *ESR1* WT/32 mutant; POG570, 68 *ESR1* WT/18 mutant).

B. Box plot representing “Estrogen Response Early” and “Estrogen Response Late” signatures enrichment levels in *ESR1* WT (n=98), Y537S (n=10) and D538G (n=8) mutant metastatic tumors from the DFCI cohort. A pairwise Mann-Whitney U test was applied comparing mutant *ESR1* to WT samples. (* $p<0.05$, ** $p<0.01$)

### Supplementary Figure S3

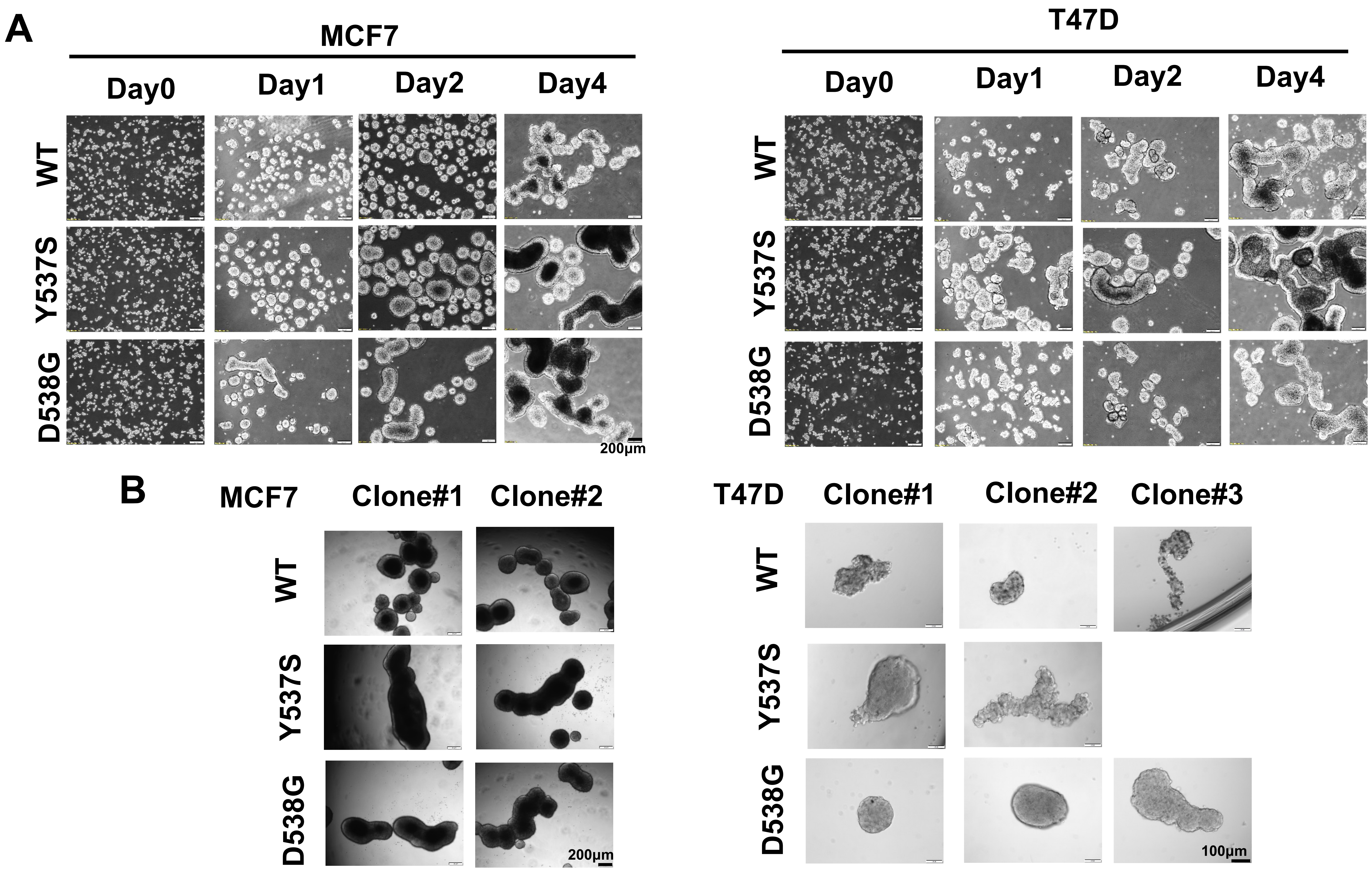


**Figure S3.** ***ESR1*** **mutant-cells form tighter clusters and exhibit ligand-independent 3D growth. (Related to Fig. 2)**

A. Representative images of MCF7 and T47D spheroids after being seeded into 6-well round bottom ULA plates at days 0, 1, 2 and 4. 4x objective was used. This experiment was done once.

B. Representative images at day 6 of MCF7 (2 clones for each cell type) and T47D (3 clones for WT and D538G, 2 clones for Y537S) individual clone spheroids after 3,000 MCF7 cells or 4,000 T47D cells were initially seeded in 96-well ULA plates. The images were captured under a 4x (MCF7) and 10x (T47D) objective. This experiment was done once.

### Supplementary Figure S4

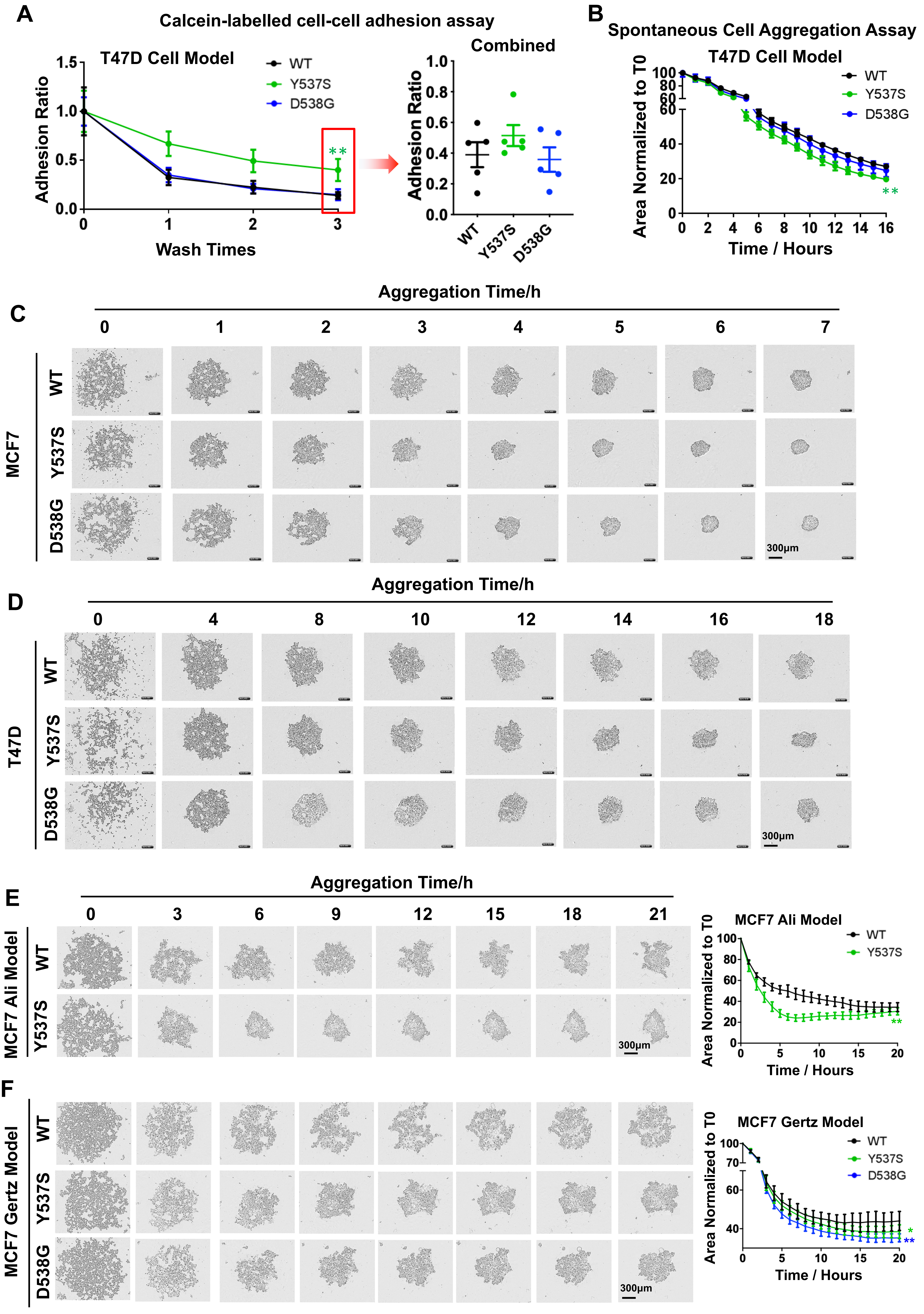

### Supplementary Figure S4-Continued

**Figure S4. *ESR1* mutant cells exhibit stronger cell-cell adhesion in static conditions. (Related to Fig.2)**
A. Left panel: A calcein labelled cell-cell adhesion assay was performed in T47D WT and mutant cells. Adhesion ratios were calculated by dividing the remaining cells after each wash to the initial readout from the unwashed wells. A pairwise two-way ANOVA between WT and each mutant was utilized. Each point represents mean ± SD with five biological replicates. Representative experiment from five independent repeats is shown. Right panel: Adhesion ratios after three washes were extracted from 5 independent experiments displayed as mean ± SEM. Dunnett's test was used to compare WT and each mutant. (* $p<0.05$, ** $p<0.01$)
B. Aggregation curves of T47D cells seeded into round bottom ULA plates. Cell aggregation progression was followed by the IncuCyte living imaging system every hour. Spheroid areas were normalized to time 0. Each bar represents mean ± SD with eight biological replicates. Representative experiment from two independent repeats is shown. A pairwise two-way ANOVA between WT and each mutant was utilized. (** $p<0.01$)
C & D. Representative images of the IncuCyte cell aggregation assay in MCF7 (S4C, 0-7 hours) and T47D (S4D, 0-18 hours) WT and mutant cells. Images were taken with a 10X objective. Representative experiments from five (MCF7) and two (T47D) independent repeats are shown.
E & F. Left panels: Representative images of IncuCyte cell aggregation assay on Ali (F) and Gertz (G) MCF7 genome-edited cell model during 0-21 hours. Images were taken with a 10X objective. Right panels: Quantfiication of the spontaneous cell aggregation seen in left panels. Each dot represents mean ± SD with eight biological replicates. These experiments were done once for each model. A pairwise two-way ANOVA between WT and each mutant was utilized. (** $p<0.01$)

### Supplementary Figure S5

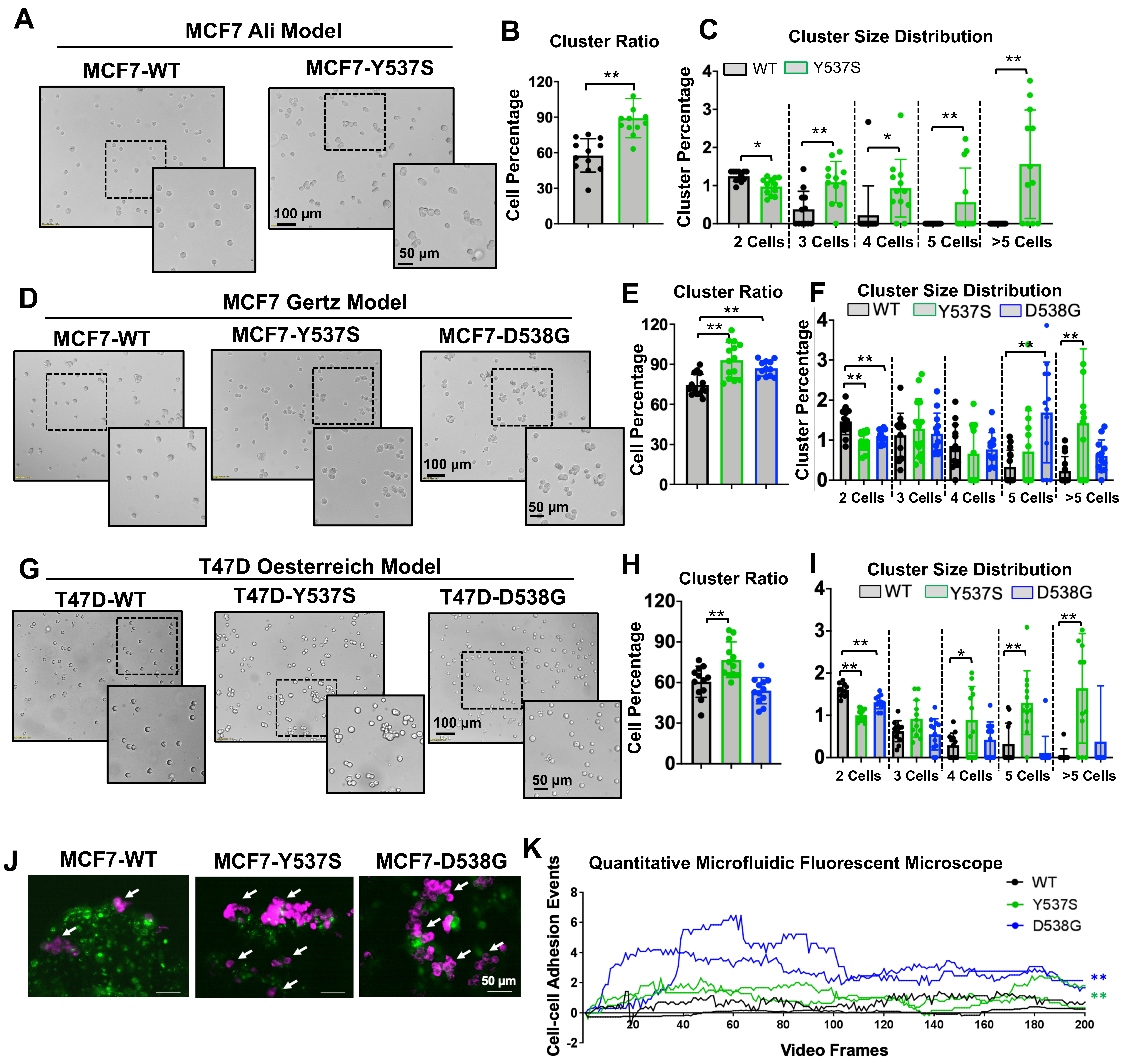


**Figure S5. *ESR1* mutant cells show enhanced cell-cell adhesion in microfluidic conditions. (Related to Fig. 2)**
A, D & G. Representative images of Ali (A) and Gertz (D) MCF7 and T47D (G) cell model cluster status after two hours of flow under physiological shear stress with the ibidi microfluidic system. Images were taken under 10x magnification. A regional 2x zoom in is presented for each image. For T47D cells, representative experiment from two independent repeats is shown. For Ali and Gertz MCF7 cells, these experiments were done once.
B, E & H. Bar graph representing the percentage of clusters in the three cell models based on the quantification of cluster and single cell numbers from 12 representative images per group. Each bar represents mean ± SD. Cells cluster ratios after 2 hours of flow were further normalized to time 0 to correct for the baseline preexisting clusters. Dunnett's test (B & H) or student's t test (E) was used between WT and mutant cells. (**p<0.01)
C, F & I. Bar plots displaying cluster size distribution among the three cell models after normalization to time 0. Each bar represents mean ± SD from 12 representative images per group. Dunnett's test or student's t test was used for statistical analysis. (*p<0.05, **p<0.01)
J & K. Representative images (J) and dynamic adhesion curves (K) of MCF7 *ESR1* mutant cells with qMFM. Imges were taken under 20x magnification (J). Adhered cells are indicated with white arrows. Each dot in K represents calculated cell-cell interaction events per frame normalized to the total calcein signals of attached cells in each video. Two representative curves for each group were shown. Dunnett's test was used for statistical analysis. This experiment was done once. (**p<0.01)

### Supplementary Figure S6

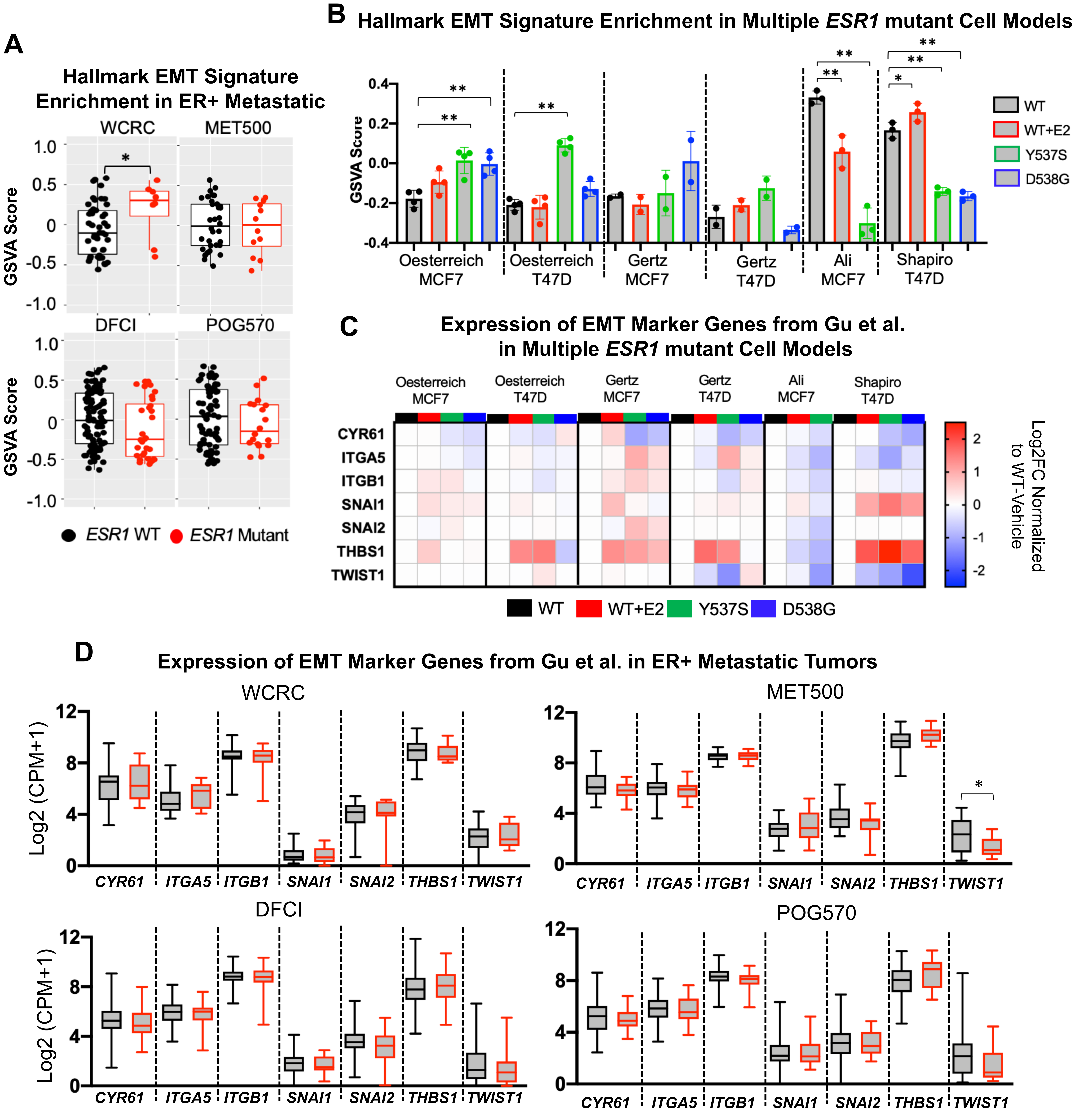


**Figure S6. EMT signature and targeted genes are not consistently altered in *ESR1* mutant tumors and cell models.**

A. Box plots representing the enrichment levels of “Hallmark EMT Pathway” in *ESR1* mutant versus *ESR1* WT metastatic tumors in each cohort. (WCRC, 46 ESR1 WT/8 mutant; MET500, 34 *ESR1* WT/12 *ESR1* mutant; DFCI, 98 *ESR1* WT/32 mutant; POG570, 68 *ESR1* WT/18 mutant). Four quantiles are shown in each plot. Mann-Whitney U test was used to compare the enrichment of the signatures in WT and mutant tumors. (* $p<0.05$)

B. Bar plot representing the enrichment levels of “Hallmark EMT Pathway” in *ESR1* mutant versus *ESR1* WT counterparts in six published genome-edited cell models. Dunnett’s test was used for each individual models. (* $p<0.05$, ** $p<0.01$)

C. A heatmap showing the $\log_2$ fold changes (normalized to WT cells) of the seven selected EMT genes in six published genome-edited *ESR1* mutant cell line models. Data were directly extracted from publicly available RNA-seq datasets. (Oesterreich: GSE89888, Ali: GSE78286; Gertz: GSE148279; Shapiro: GSE108304)

D. Box plots representing expression level of the seven selected EMT marker genes in *ESR1* mutant versus *ESR1* WT metastatic tumors in each cohort. (WCRC, 46 *ESR1* WT/8 mutant; MET500, 34 *ESR1* WT/12 ESR1 mutant; DFCI, 98 *ESR1* WT/32 mutant; POG570, 68 *ESR1* WT/18 mutant). Four quantiles are shown in each plot. Mann-Whitney U test was used to compare the log2 (CPM+1) values of each gene in WT and mutant tumors. (* $p<0.05$)

### Supplementary Figure S7

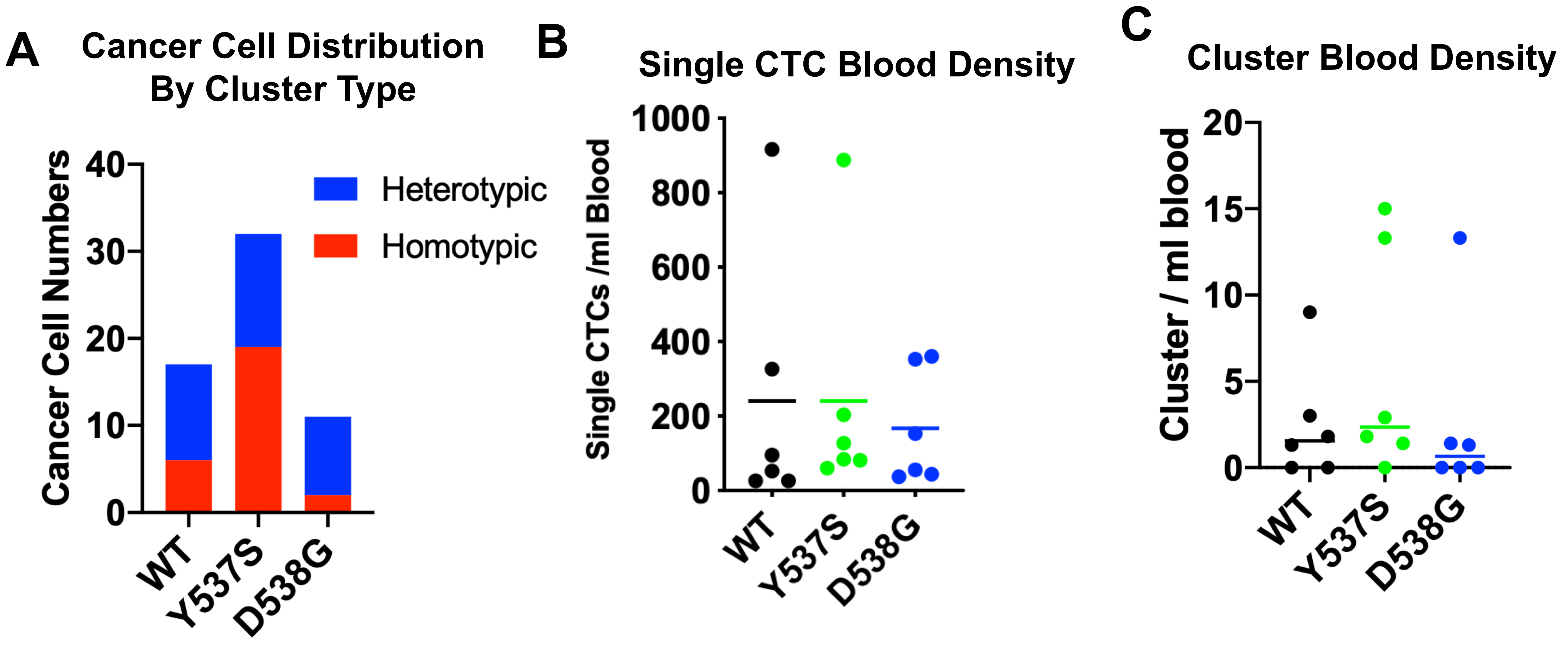


**Figure S7. *ESR1* mutant cells exhibit enhanced metastases *in vivo*. (Related to Fig. 2)**
A. Stacked bar plot showing total number of CTC clusters per mutation subtype and the distribution of cancer cells in homo- or heterotypic clusters. Fisher's exact test was used for statistical analysis.
B&C. Dot plot displaying single CTC (B) and CTC cluster (C) concentrations measured from each mouse. Single CTCs and cluster concentrations were calculated by normalizing single CTC or cluster number from each sample to corresponding blood volumes. Two samples were measured for each mouse, average densities from the two samples were shown. Six mouse were measured for each group. Mann-Whitney U test was performed.

### Supplementary Figure S8

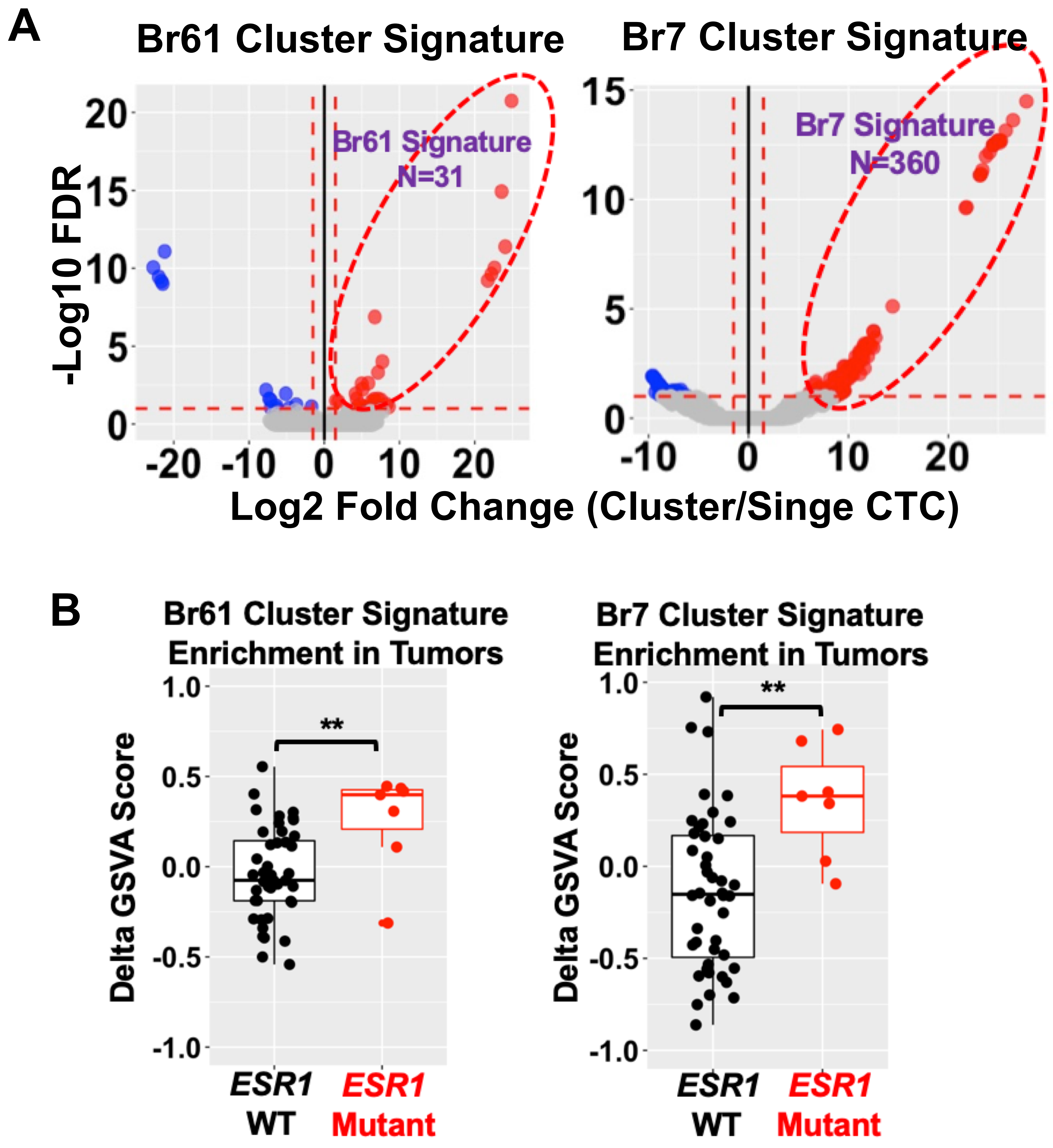


**Figure S8. CTC cluster-derived gene signatures are significantly enriched in *ESR1* mutant metastases. (Related to Fig. 2)**

A. Volcano plots showing the differentially regulated genes in RNA-sequenced CTC clusters versus single CTCs from two patients with ER+ breast cancer (Br61, 7 CTC clusters vs 7 single CTCs; Br7, 2 CTC clusters vs 2 single CTCs). Upregulated genes are circled and used as the CTC cluster gene signatures. Data were download from Gkountela et al. (GSE111065).

B. Gene Set Variation Analyses were performed on patient matched primary-metastatic paired samples examining the enrichment of the two CTC cluster gene signatures derived from S7A. Delta GSVA score of each sample was calculated by subtracting the scores of primary tumors from the matched metastatic tumors. Median values in each group were labeled. Mann-Whitney U test was performed to compare the delta GSVA scores between *ESR1* WT (n=44) or mutation (n=7) harboring tumors. (** $p<0.05$)

### Supplementary Figure S9

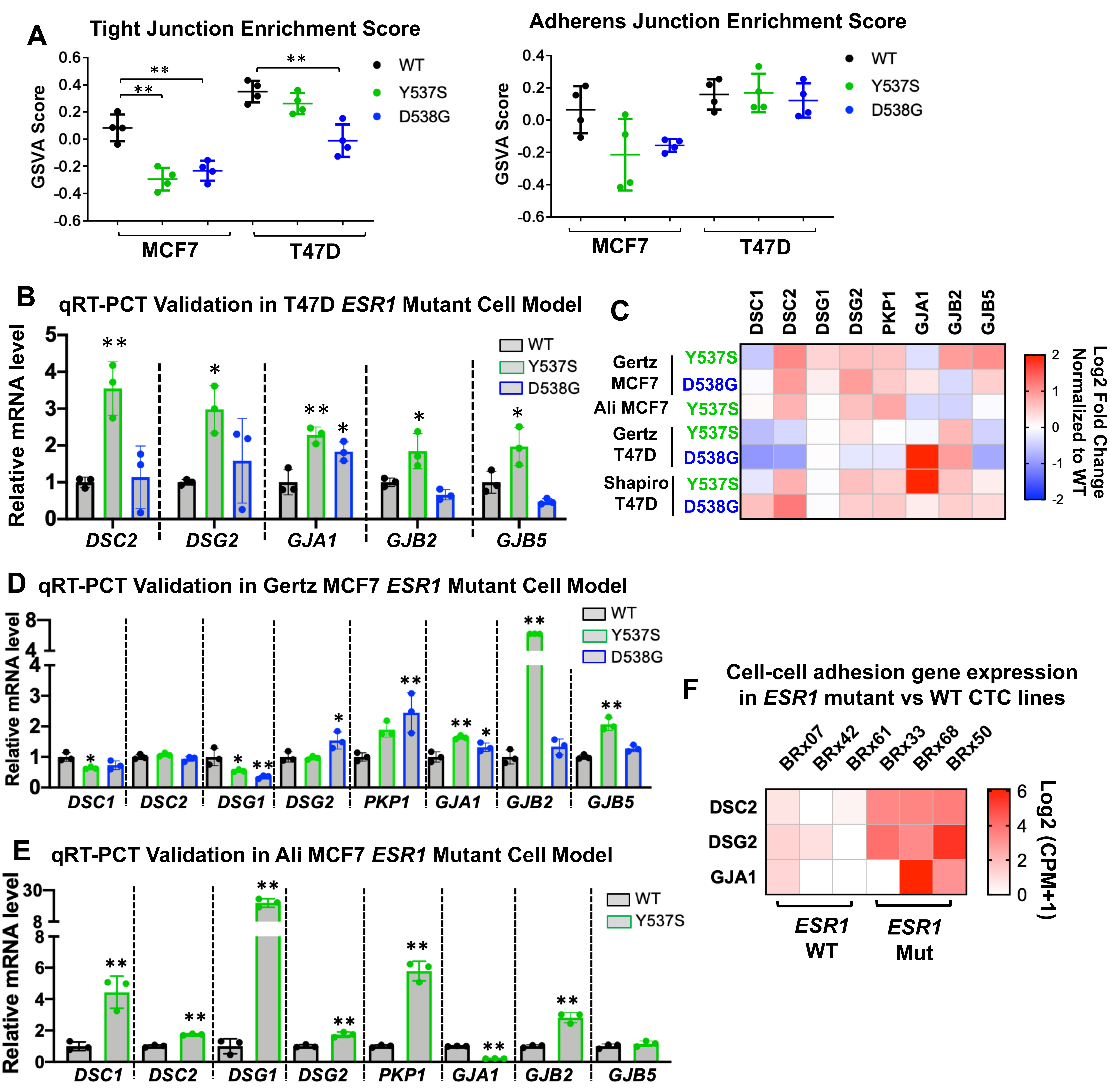


**Figure S9. Desmosome and gap junction genes are increased in *ESR1* mutant cells. (Related to Fig. 3)**

A. Gene Set Variation Analysis (GSVA) scores of tight junction and adherens junction gene sets enrichment in MCF7 and T47D *ESR1* mutant vs WT cell RNA-seq data sets. Each cell type has four biological replicates. Dunnett's test was used to test the significance between WT and mutant cell lines. (** p<0.01)

B, D & E. qRT-PCR validation of selected altered candidate desmosome and gap junction genes in T47D (B), Gertz (D) and Ali (E) MCF7 *ESR1* mutant cells. ΔΔCt method was used to analyze relative mRNA fold changes normalized to WT cells and *RPLP0* levels were measured as an internal control. Each bar represents mean ± SD performed in biological triplicates. For T47D cells, representative experiment from two independent repeats is shown. For Ali and Gertz MCF7 cells, these experiments were done once. Dunnett's test (B & D) and student's t test (E) were used to compare the gene expression between WT and mutants. (* p<0.05; ** p<0.01)

C. A heatmap showing the $\log_2$ fold changes (normalized to WT cells) of the eight selected cell-cell adhesome genes in seven other genome-edited *ESR1* mutant cell line models. Data were directly extracted from publicly available RNA-seq datasets. (Ali: GSE78286; Gertz: GSE148279; Shapiro: GSE108304)

F. Comparison of *DSC2, DSG2 and GJA1* gene expression levels between 3 *ESR1* mutant and 3 *ESR1* WT *ex vivo* circulating tumor cell lines reported by Yu et al.. Data was extracted from publicly available RNA-sequencing (GSE55807) and $\log_2$(CPM+1) was used for visualization.

### Supplementary Figure S10

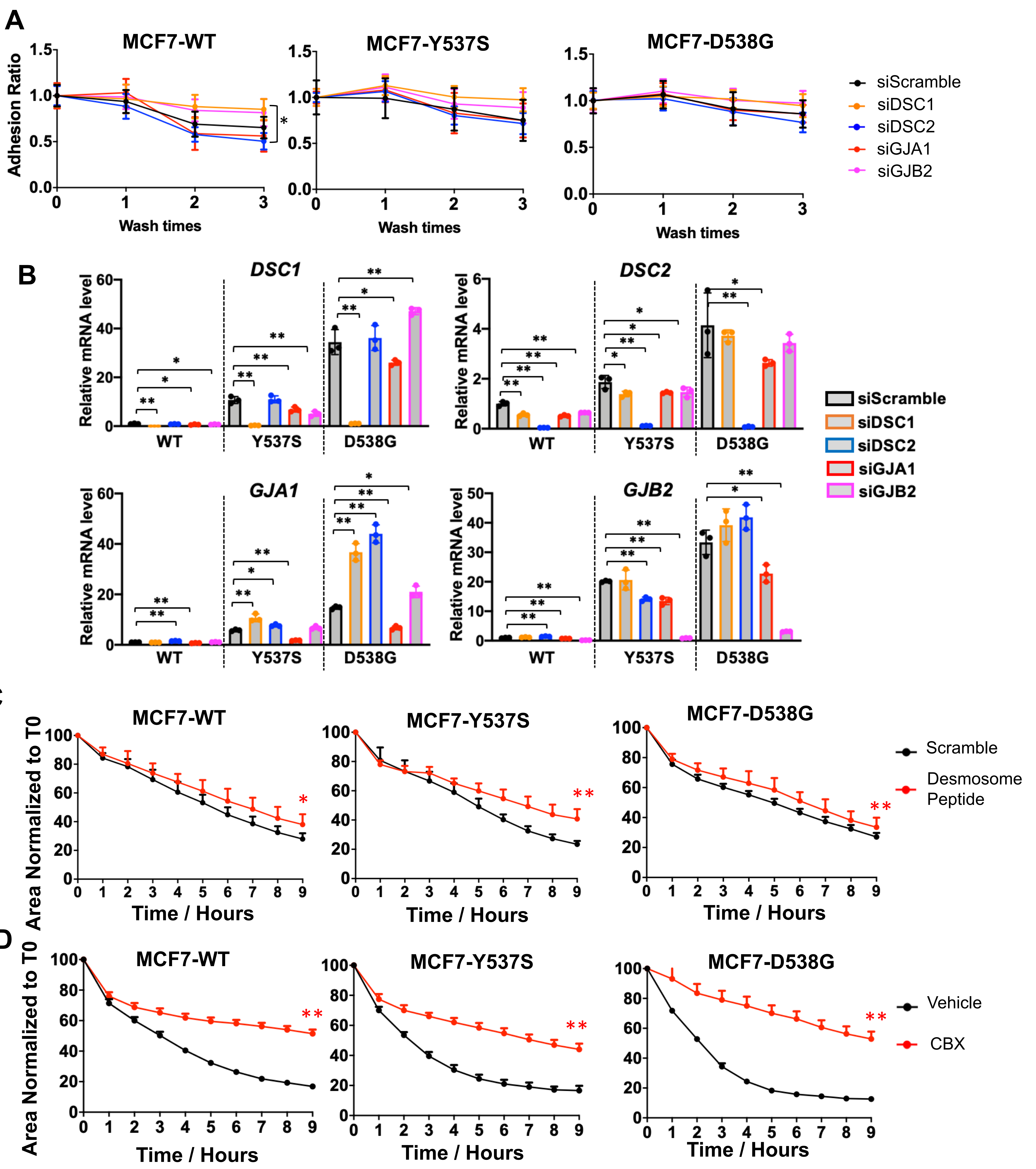


**Figure S10. Functional characterization of desmosome and gap junctions. (Related to Fig. 3)**

A. Line plot representing the percentages of calcein-labelled adhesive MCF7 cells after three washes, normalized to unwashed readouts, after a 24 hour transfection of siRNA targeting scramble control, *DSC1*, *DSC2*, *GJA1* or *GJB2*. Adhesion ratios were calculated by dividing the remaining cells after each wash to the initial readout from unwashed wells. Each point represents mean ± SD with five biological replicates.This experiment was done once. A pairwise two-way ANOVA between siControl and each gene knockdown group was applied. (* $p<0.05$)

B. Bar plots representing qRT-PCR measurement of *DSC1*, *DSC2*, *GJA1* and *GJB2* mRNA levels in MCF7 *ESR1* breast cancer cells with siRNA knockdown of each of the four genes. ΔΔCt method was used to analyze relative mRNA fold changes normalized to WT cells and *RPLP0* levels were measured as an internal control. Each bar represents mean ± SD. This experiment was done once. Dunnett’s test was used to compare each gene knockdown group to siScramble counterparts. (* $p<0.05$; ** $p<0.01$)

C & D. Line plot representing the aggregation ratio of MCF7 *ESR1* WT and mutant cells seeded into round bottom ULA plates. Cells were treated with or without 300μM of desmosomal blocking peptide (C) or 100μM of carbenoxolone (D). Cell aggregation processes were followed by the IncuCyte living imaging system every hour. Spheroid areas were normalized to time 0. Each bar represents mean ± SD with eight biological replicates. The experiment with desmosome peptide treatment was done once whereas the experiment with CBX treatment is representative from three independent repeats. A pairwise two-way ANOVA between vehicle or scramble controls and CBX or desmosome peptide was utilized. (* $p<0.05$; ** $p<0.01$)

### Supplementary Figure S11

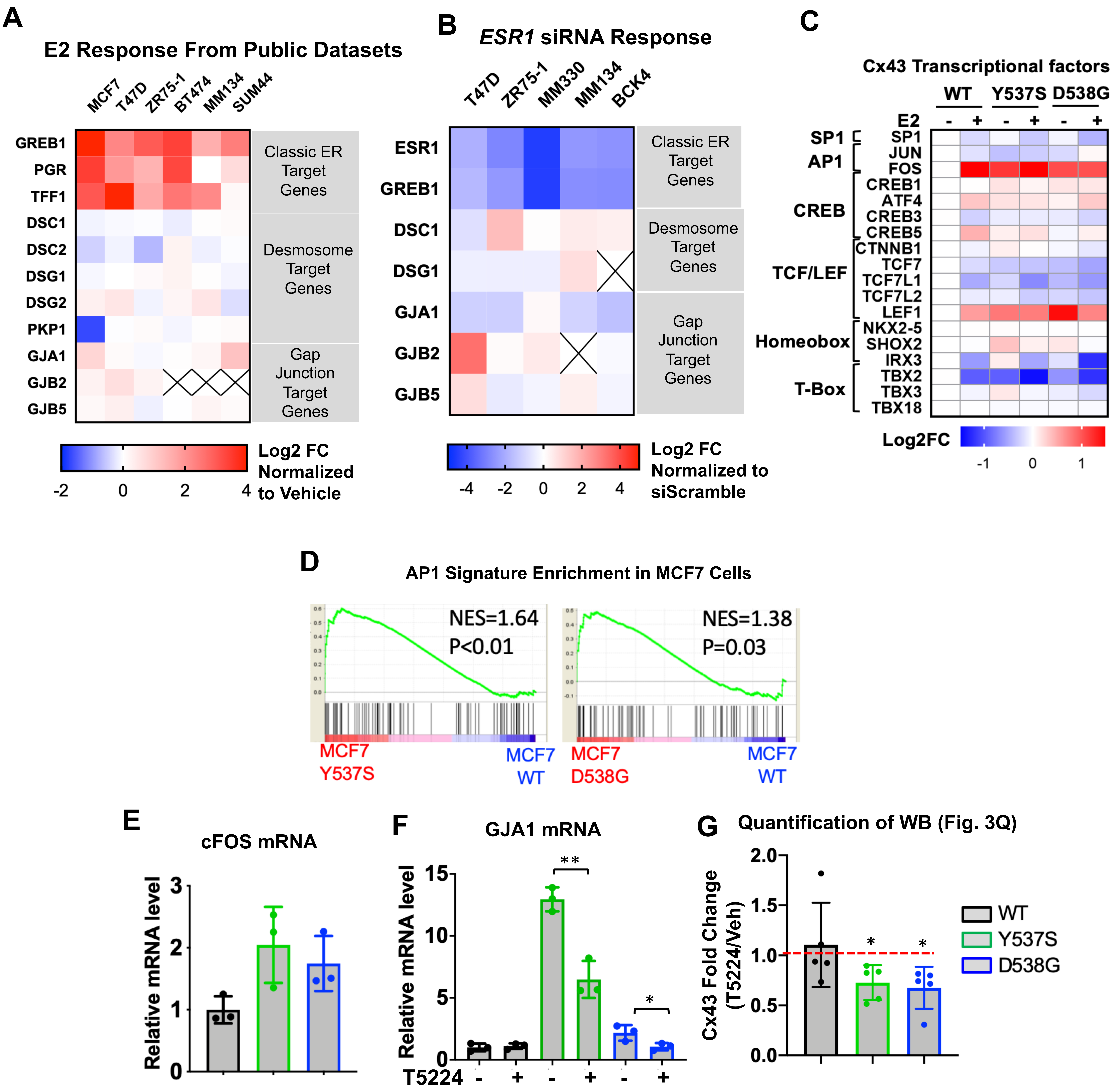


**Figure S11. GJA1 induction is mediated via a cFOS-associated secondary transactivation in *ESR1* mutant cells. (Related to Fig. 3)**

A. A heatmap representing the expression fold changes of the eight cell-cell adhesome genes under E2 treatment in 6 ER+ breast cancer lines from publicly available datasets (GSE89888, GSE3834, GSE38132 and GSE50693). Three classic E2-regulated genes were set as positive controls..

B. A heatmap showing the Log2 fold change of five target cell-cell adhesion genes after seven days of *ESR1* knockdown in five ER+ cell lines. Reduction of three classic E2-regulated genes were set as controls. These experiments were done once for each cell line.

C. Heatmap representing the expression levels of the 18 transcription factors associated with *GJA1*in MCF7 *ESR1* WT and mutant cells. Data was extracted from RNA-seq analysis (GSE89888) with four biological replicates and was normalized to WT-vehicle groups.

D. Gene set enrichment plots showing the comparison of enrichment levels of PID-AP1 gene signature (MSigDB, M167) between MCF7-WT and MCF7-Y537S (left panel) and MCF7-WT and MCF7-D538G (right panel). New enrichment scores (NES) and p-values are labelled on the plots.

E & F. qPCR measurement of *cFOS* (E) basal levels and *GJA1* (F) mRNA expression with or without 20μM of T-5224 treatment for 3 days in MCF7 *ESR1* WT and mutant cells. ΔΔCt method was used to analyze relative mRNA fold changes normalized to WT cells and *RPLP0* levels were measured as an internal control. Each bar represents mean ± SD of biological triplicates. Both experiments were done once. Dunnett's test (E) and Student's t test (F) was used. (* p<0.05; **p<0.01)

G. Bar graph showing the fold changes of connexin 43 expression after T5224 treatment normalized to vehicle groups based on the quantification of western blots in Fig. 3Q. Each bar represents mean ± SD from five independent experiments. (* p<0.05)

Supplementary Figure S12

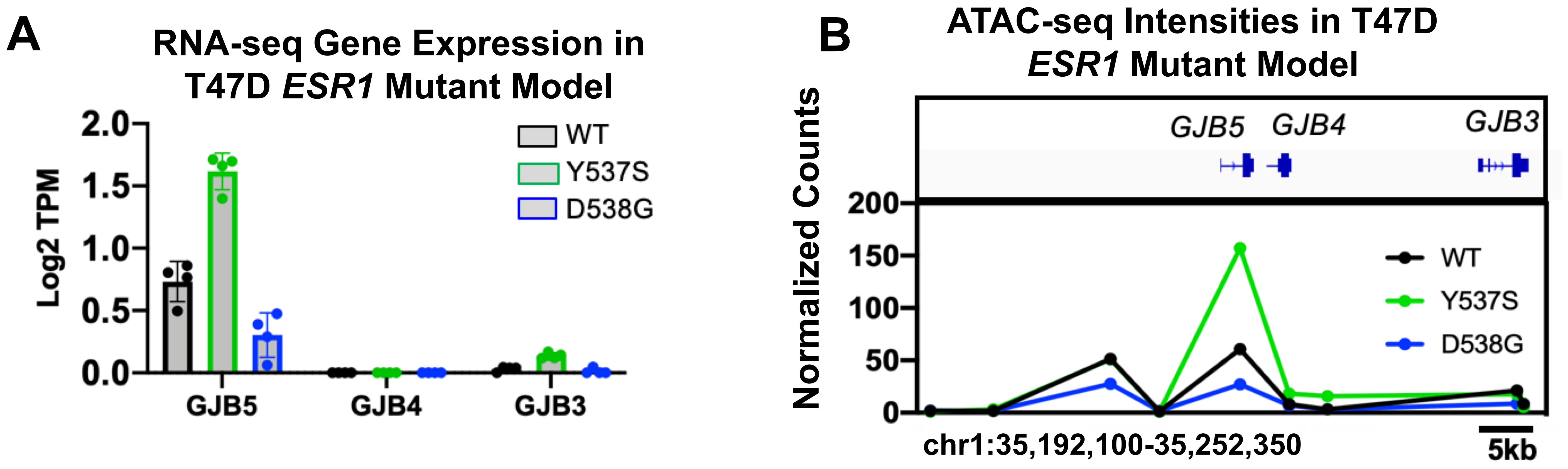


**Figure S12. Accessible chromatin remodeling is associated with increased cell-cell adhesome genes in *ESR1* mutant cells. (Related to Fig. 3)**

A. Bar plot showing the expressional level of *GJB5, GJB4* and *GJB3* in T47D *ESR1* mutant cells based on $\log_2$(TPM+1) values from RNA-seq data with four biological replicates from GSE89888.

B. Dot plot representing ATAC-seq peak signals from nine consecutive sites at *GJB5*/*GJB4*/*GJB3* loci. Average ATAC-seq counts from two clones were calculated to represent each cell type.Y axis represents normalized counts of each peak. ATAC-seq data were downloaded from GSE148279.

### Supplementary Figure S13

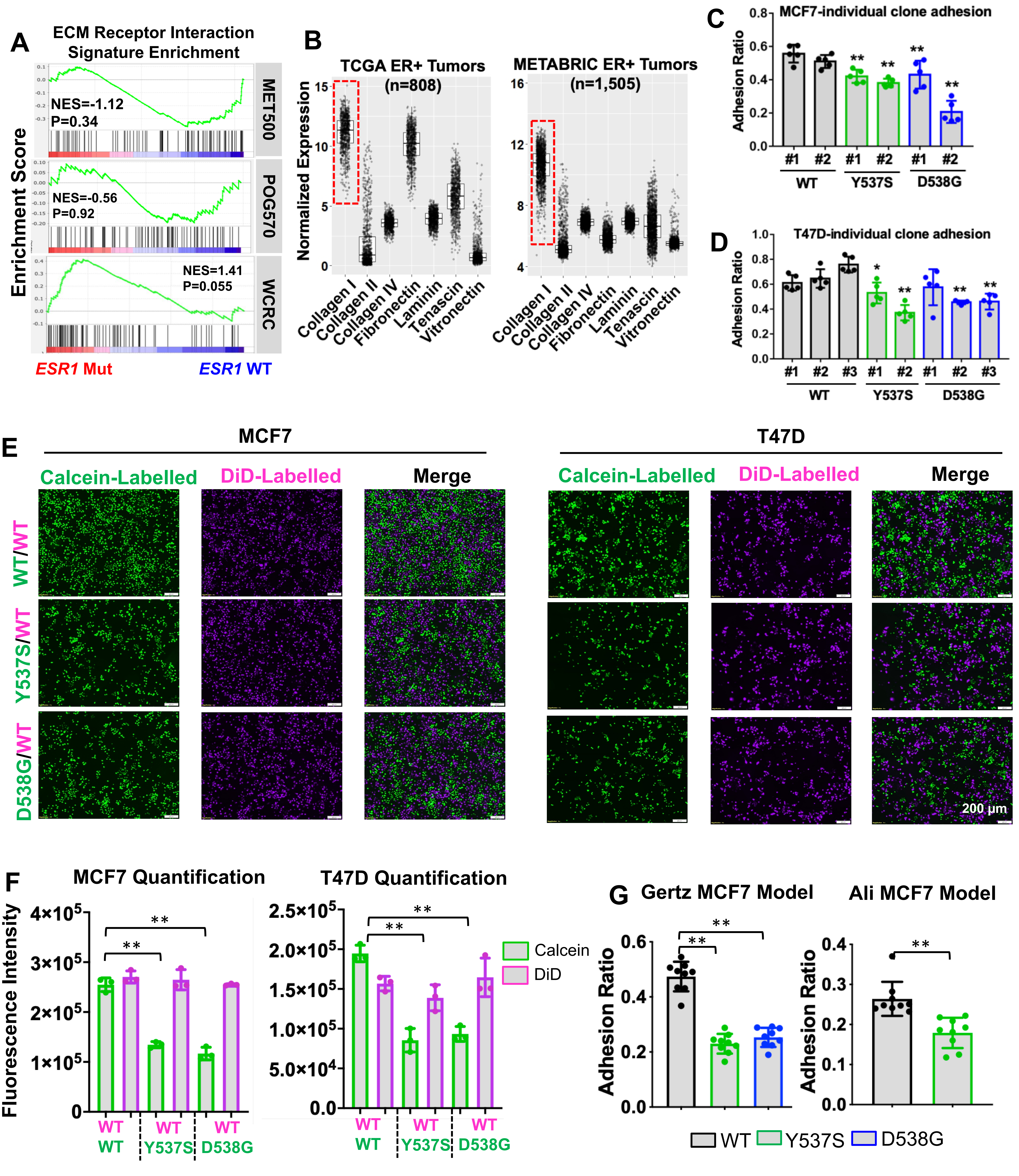

### Supplementary Figure S13-Continued

**Figure S13. *ESR1* mutant cells show diminished adhesion on collagen I. (Related to Fig. 4)**
A. Gene set enrichment plots showing the comparison of enrichment levels of the "KEGG ECM Receptor Interaction" gene set (MSigDB, M7098) between WT and mutant tumors from three ER+ metastatic tumor cohorts. (MET500, 34 ESR1 WT/12 ESR1 mutant; POG570, 68 ESR1 WT/18 mutant; WCRC, 46 ESR1 WT/8 mutant)
B. Box plots with individual values of seven ECM component abundance from ER+ tumors in the TCGA and METABRIC cohorts. The abundance calculations were based on normalized gene(s) expression with $log_2$(TPM+1) values (TCGA) or $log_2$ normalized probe intensities (METABRIC). Collagen I was highlighted as the most abundant component.
C & D. Quantification of adhesion ratios on collagen I for each cell types' individual clones, MCF7 (B) and T47D (C). Bar graphs represent the mean ± SD with five biological replicates in each group. Dunnett's test was utilized within each cell line to compare between the average of WT clones to each single mutant clone. These experiments were done once for each cell line. (* p<0.05, ** p<0.01)
E. Representative images of the co-culture adhesion assay in collagen I of MCF7 (left panel) and T47D (right panel) cell lines. WT cells with DiD (pink) labelling were equally mixed with calcein (green) labelled WT/Y537S/D538G cells. The mixed cells were used in the adhesion assay, and represnetative images of either single GFP/Cy5 channels or merged channels are shown. Images were taken under 4x magnification. Representative experiment from two independent repeats is presented. (** p<0.01)
F. Quantification of GFP or Cy5 single channel signal intensities from S12D. Bar graphs represent the mean ± SD with three biological replicates in each group. Dunnett's test was used to compare the signal intensities between WT and each mutant cell within each individual channel.
G. Adhesion assay on collagen I with Gertz and Ali MCF7 genome-edited *ESR1* mutant cell models. Bar graphs represent the mean ± SD with nine biological replicates in each group. These experiments were done once for each cell model. Dunnett's test (Gerz) and Student's t test (Ali) was utilized to compare adhesion ratio between WT and mutant cell.  (* p<0.05, ** p<0.01).

### Supplementary Figure S14

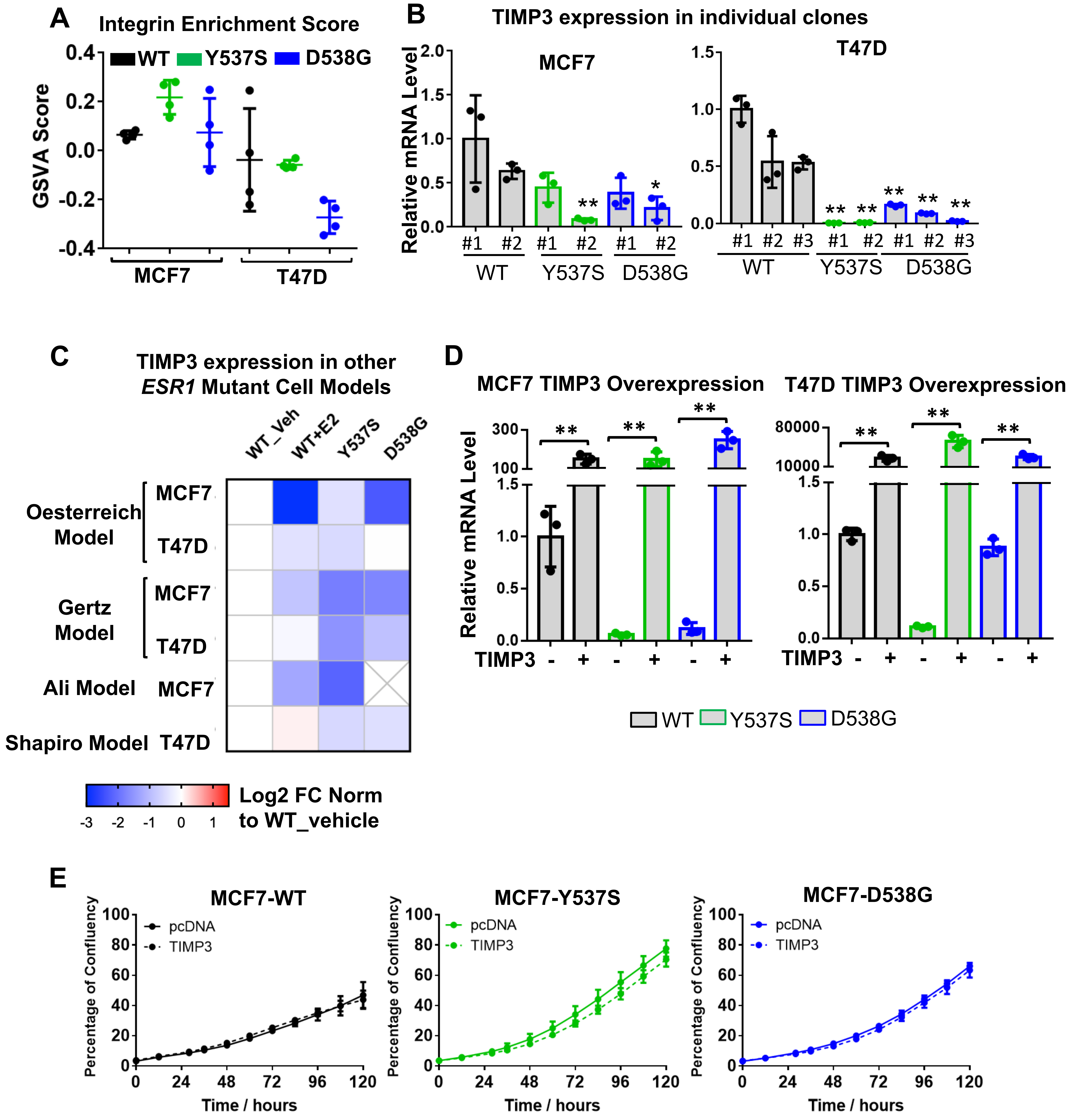


**Figure S14. Decreased *TIMP3* expression drives the loss of collagen adhesion property in *ESR1* mutant cells. (Related to Fig. 4)**

A. Gene Set Variation Analysis (GSVA) scores of integrin gene set enrichment in MCF7 and T47D *ESR1* mutant vs WT cell RNA-seq data sets. Each cell type has four biological replicates. Dunnett's test was used to test the significance between WT and mutant cell lines.

B. qRT-PCR validation of *TIMP3* expression in WT and *ESR1* mutant cells in individual clones. Ct values were normalized to *RPLP0* and further normalized to the WT#1 clone. Bar graphs represent the mean ± SD with biological triplicates in each group. Dunnett's test was utilized within each cell line to compare between the average of WT clones to each single mutant clone. This experiment was done once. (* $p<0.05$; ** $p<0.01$)

C. Expression fold change of *TIMP3* in *ESR1* mutant cells normalized to their WT controls from seven genome-edited cell models. Fold changes were calculated from RNA-seq data except for Ali and Gertz MCF7 models in which independent replicates of qPCR data was used.

D. qRT-PCR validation of *TIMP3* overexpression in MCF7 and T47D cells. ΔΔCt method was used to analyze relative mRNA fold changes normalized to WT-pcDNA cells and *RPLP0* levels were measured as an internal control. Each bar represents mean ± SD of biological triplicates. This experiment was done once. Pair-wised student's t test was applied to compare each pcDNA and TIMP3 overexpression group. (** $p<0.01$)

E. Time course growth comparison of MCF7 WT and *ESR1* mutant cells with pcDNA empty vector or *TIMP3* transfection in the absence of E2. Each dot represents mean ± SD with five biological replicates. A two-way ANOVA was used to test the effects of *TIMP3* overexpression in each cell type.

### Supplementary Figure S15

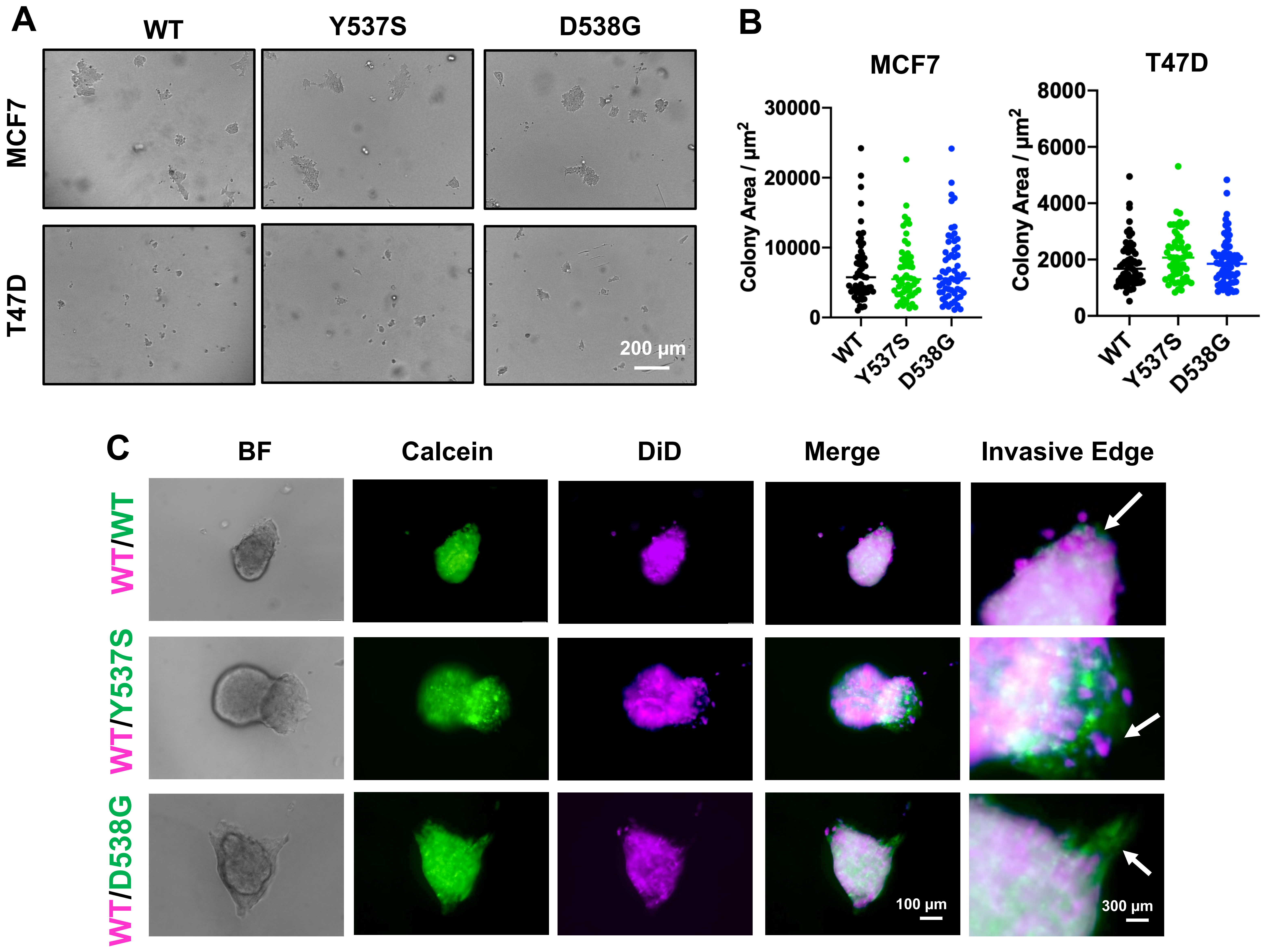


**Figure S15. *ESR1* mutant cells show enhanced invasion in collagen I which can be inhibited by a MMP inhibitor. (Related to Fig. 4)**

A. Representative images of the growth of *ESR1* WT and mutant cells in collagen I after 4 (MCF7) and 6 (T47D) days. Images were taken under 4x magnification. This experiment was performed in biological triplicates for once.

B. Dot plots showing quantification of colony size of *ESR1* WT and mutant cells in collagen I from (A). Around 60 individual colonies were analyzed per group from biological triplicates. This experiment was performed in biological triplicates for once. Dunnet's test was used for statistical analysis in each cell line.

C. Representative images of spheroid co-culture invasion in type I collagen. T47D WT cells with DiD (pink) labelling were equally mixed with calcein (green) labelled T47D WT/Y537S/D538G cells. The mixed spheroids were formed in ULA plates and collagen I was loaded into each well. Images were taken after 6 days under 10x magnification. Invasive edges with GFP and Cy5 channels are identified with white arrows and zoomed for there times shown in the right panel. This experiment was performed in biological triplicates for once.

### Supplementary Figure S16

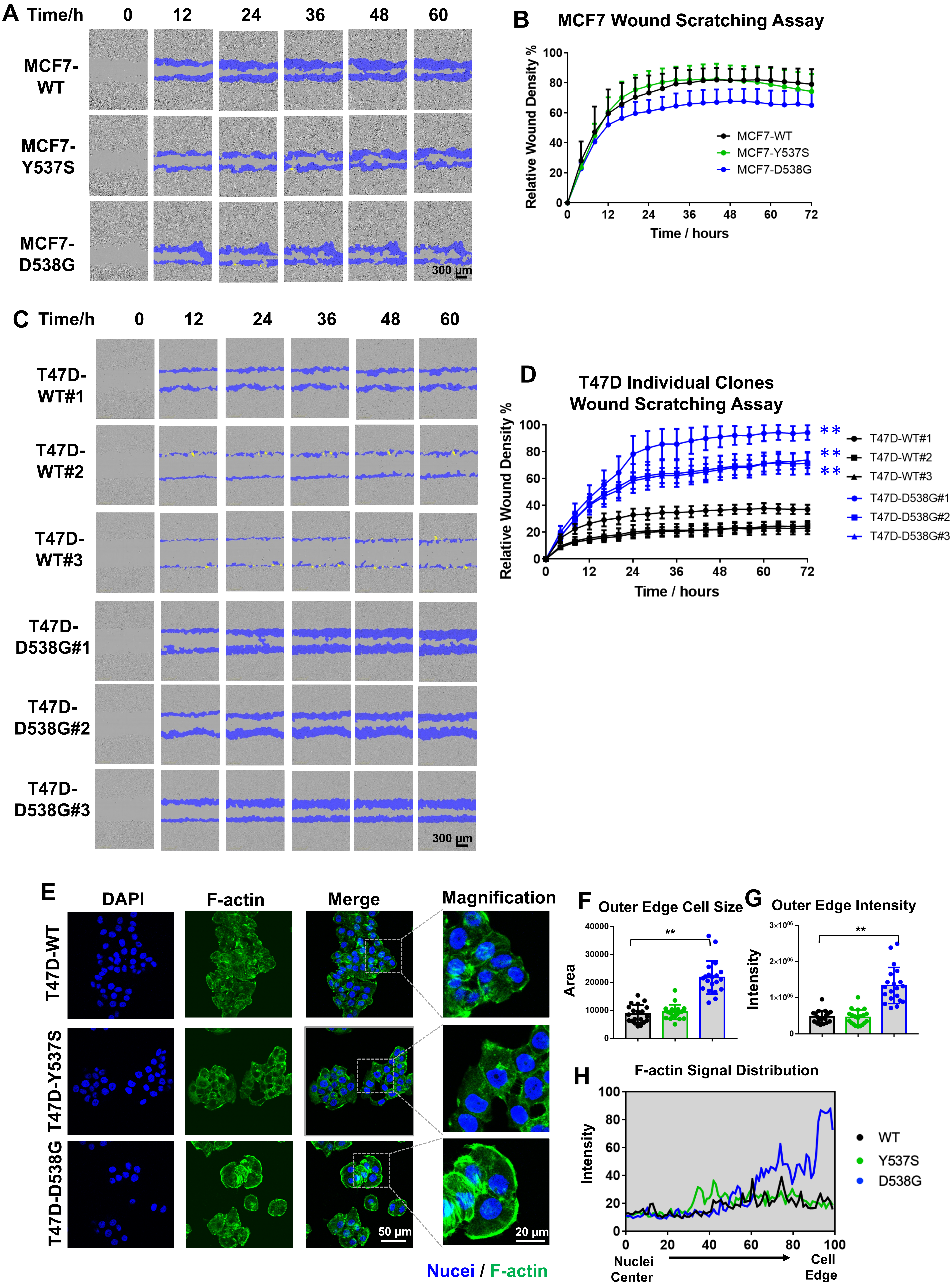

Supplementary Figure S15-Continued

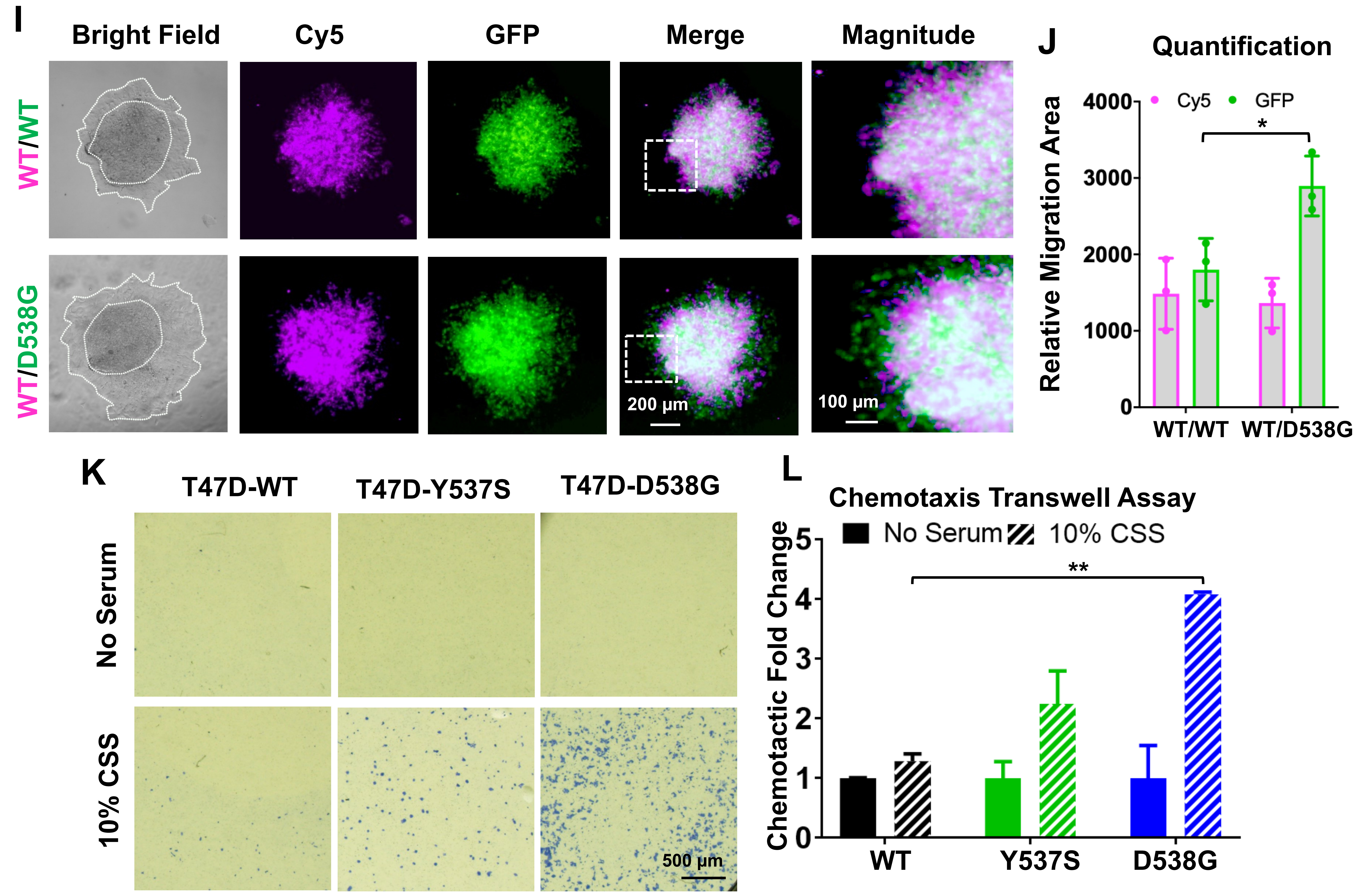


**Figure S16. T47D-D538G cells display enhanced migratory properties. (Related to Fig. 5)**

A & B. Representative images (A) and quantification (B) of wound scratch assay performed with MCF7 *ESR1* WT and mutant cells using IncuCyte living imaging system over 72 hours. The migratory region normalized to T0 are labelled in blue. Images were taken under 10x magnification. Cell migration rates were quantified based on relative wound densities with eight biological replicates. Representative experiment from two independent repeats is shown. Pairwise two-way ANOVA between WT and each mutant was performed.

C & D. Confirmation of migratory alterations in T47D WT and T47D-D538G individual clones for 72 hours with representative images (C) and quantification plot (D). Cell migration rates were quantified based on relative wound densities with eight biological replicates. Pairwise two-way ANOVA was used to test the significance between the mean of WT cells and each T47D-D538G clone. (** $p<0.01$)

E. Representative images of F-actin staining on T47D *ESR1* WT and mutant cell models. Images were taken under 60x magnification. Representative cell edges were further zoomed in the right panel. This experiment was done once with five different regions captured for each group.

F & G. Bar plot representing quantification of outer cell edge F-actin size (G) and intensities (H) from 20 representive cells. 4 cells were randomely selected from 5 regions per group. This experiment was done once. Dunnett's test was applied between WT and each mutant. (** $p<0.01$)

H. Line plot showing the signal intensity distribution from nuclear center to cell edge. Distances measured from each cell were normalized to a 0-100 scale. Representative curve from one cell of each group is shown.

I & J. Representative images of spheroid co-culture collective migration in type I collagen (I). T47D WT cells with DiD (pink) labelling were equally mixed with calcein (green) labelled T47D WT/D538G cells in a 96-well round bottom ULA-plate to form spheroids for 3 days. Mixed spheroids were then transferred to collagen I coated plates for collective migration. 4X objectives were used for the entire spheroids and 10X objectives for the migratory edge. Migration distances of Cy5 and GFP signals were calculated separately (M). This experiment was done once with biological triplicates. Student's t test was used for statistical analysis. (* $p<0.05$)

K & L. Representative images (I) and quantification results (J) of transwell migration asssy using 10% CSS as chemoattractant with T47D *ESR1* WT and mutant cell. No serum serves as negative control. Bilogical dulicates were used for each group. Representative experiment from two independent repeats is shown. Dunnett's test was used for statistical analysis between WT and each mutant within the 10% CSS group. ( ** $p<0.01$)

### Supplementary Figure S17

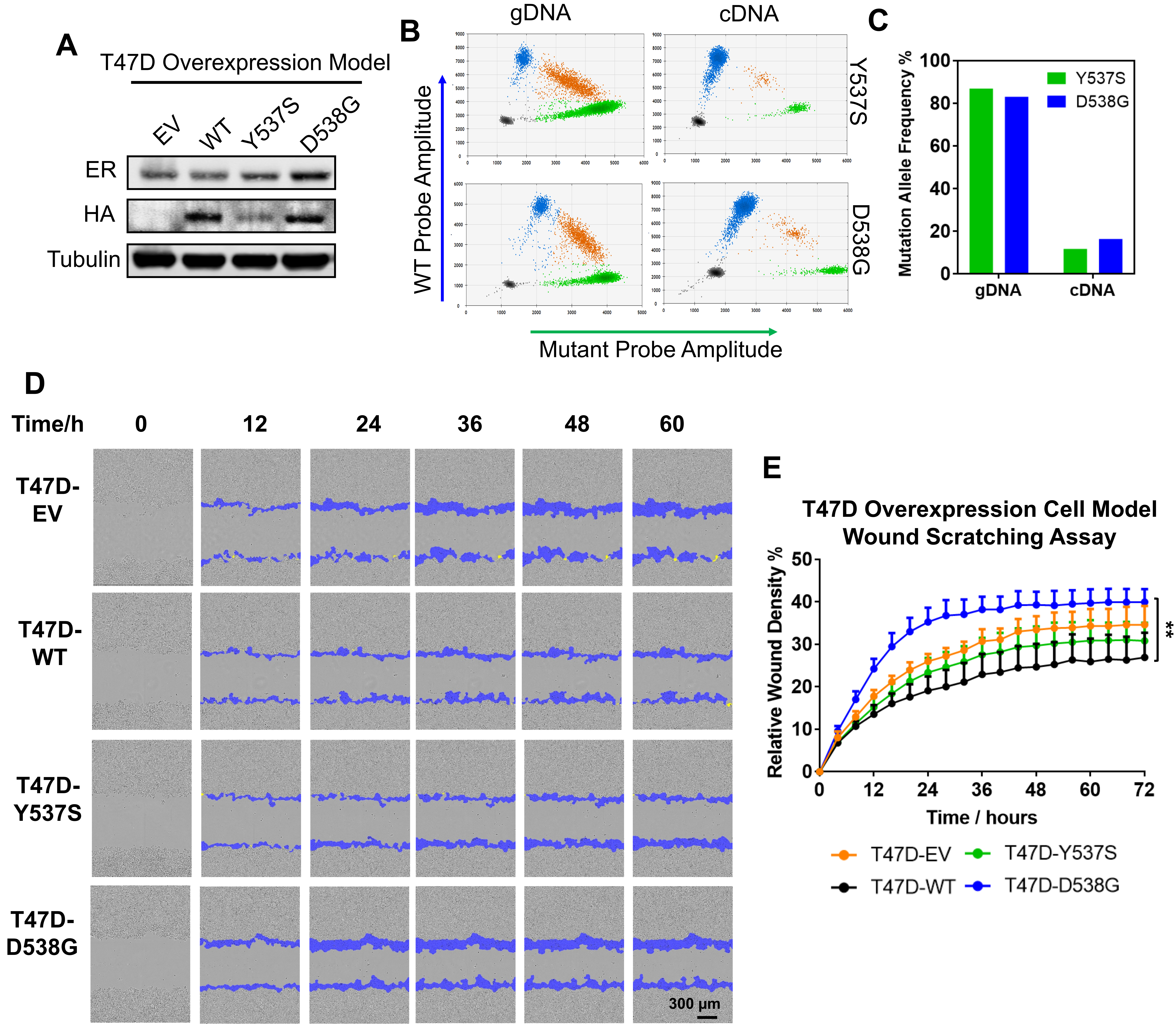


**Figure S17. Enhanced migration is recapitulated in T47D overexpressing D538G cell model. (Related to Fig. 5)**

A. Immunoblot validation of T47D *ESR1* mutant overexpressing cells. Both total and overexpressing ER (HA-tagged) are presented with tubulin used as a loading control. This validation experiment was done once.

B & C. ddPCR validation of mutation allele frequency in Y537S and D538G overexpressing cell model at both DNA (gDNA) and RNA (cDNA) levels. 2D scatter plots of ddPCR measurement are displayed in B. Black dots reflect no DNA droplet, blue and green dots show droplets containing solely WT or mutant DNA, organe dots represents droplets containing both WT and mutant DNA copies. Quantifcation of mutation allele frequencies is shown in C. This experiment was done once.

D & E. IncuCyte wound scratch assay on T47D overexpressing WT and mutant cells using IncuCyte living imaging system for 72 hours. Representative images (D) and quantification curves (E) are shown. The migratory region normalized to T0 are labelled in blue. Images were taken under 10x magnification. Cell migration rates were quantified based on relative wound densities with eight biological replicates. Representative experiment from two independent repeats is shown. Pairwise two-way ANOVA between WT and each mutant was performed. (** $p<0.01$)

### Supplementary Figure S18

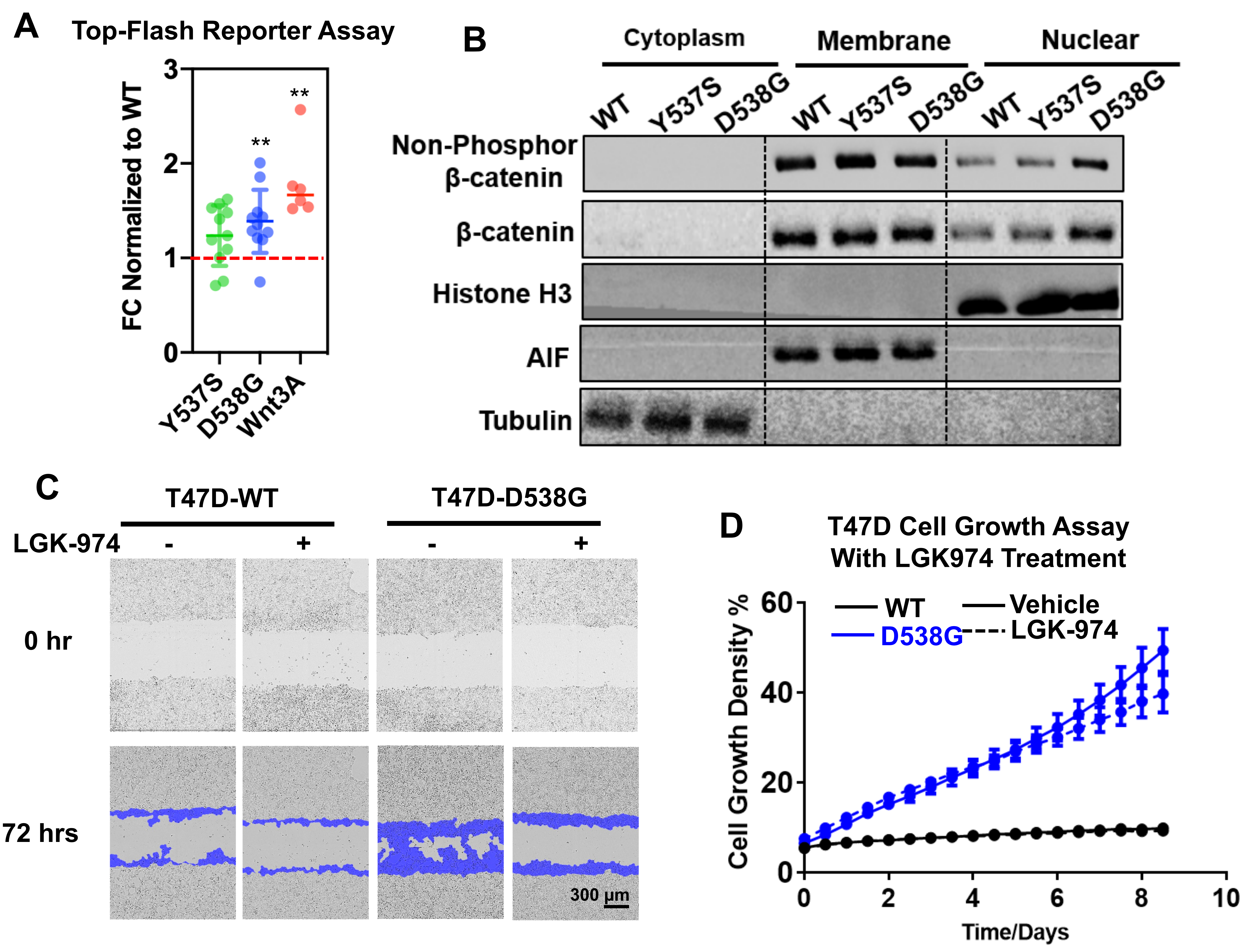


**Figure S18. Wnt hyperactivation drives cell migration in T47D-D538G cells. (Related to Fig. 5)**
A. Top-flash luciferase assay on T47D ESR1 mutant cells at basal levels. Luminescence readouts of top-flash were normalized to Renilla readouts, with fold changes calculated by normalizing to WT cells. Six independent experiments with Wnt3A-induction in T47D WT cells were used as positive controls. The results were combined from eleven independent experiments. Dunnett's test was applied between WT cells and each mutant. (** p<0.01)
B. Immunoblot detection of non-phosphor (active form) and total β-catenin levels in different cell fractions of T47D *ESR1* WT and mutant cells. Histone H3, AIF and tubulin are detected as markers for nuclear, membrane and cytoplasmic portions, respectively. This experiment was done once.
C. Representative images of IncuCyte wound scratch assay with or without 5μM LGK974 treatment for 72 hours. The migratory region normalized to T0 are labelled in blue. Images were taken under 10x magnification. Cell migration rates were quantified based on relative wound densities with eight biological replicates. Representative experiment from three independent repeats is shown. Pairwise two-way ANOVA between WT and each mutant was performed. (** p<0.01)
D. Time course growth comparison of T47D WT and D538G *ESR1* mutant with or without 5μM LGK974 treatment for 9 days in the absence of E2. Each dot represents mean ± SD with five biological replicates. This experiment was done once. A two-way ANOVA was used to test the effects of LGK974 treatment.

### Supplementary Figure S19

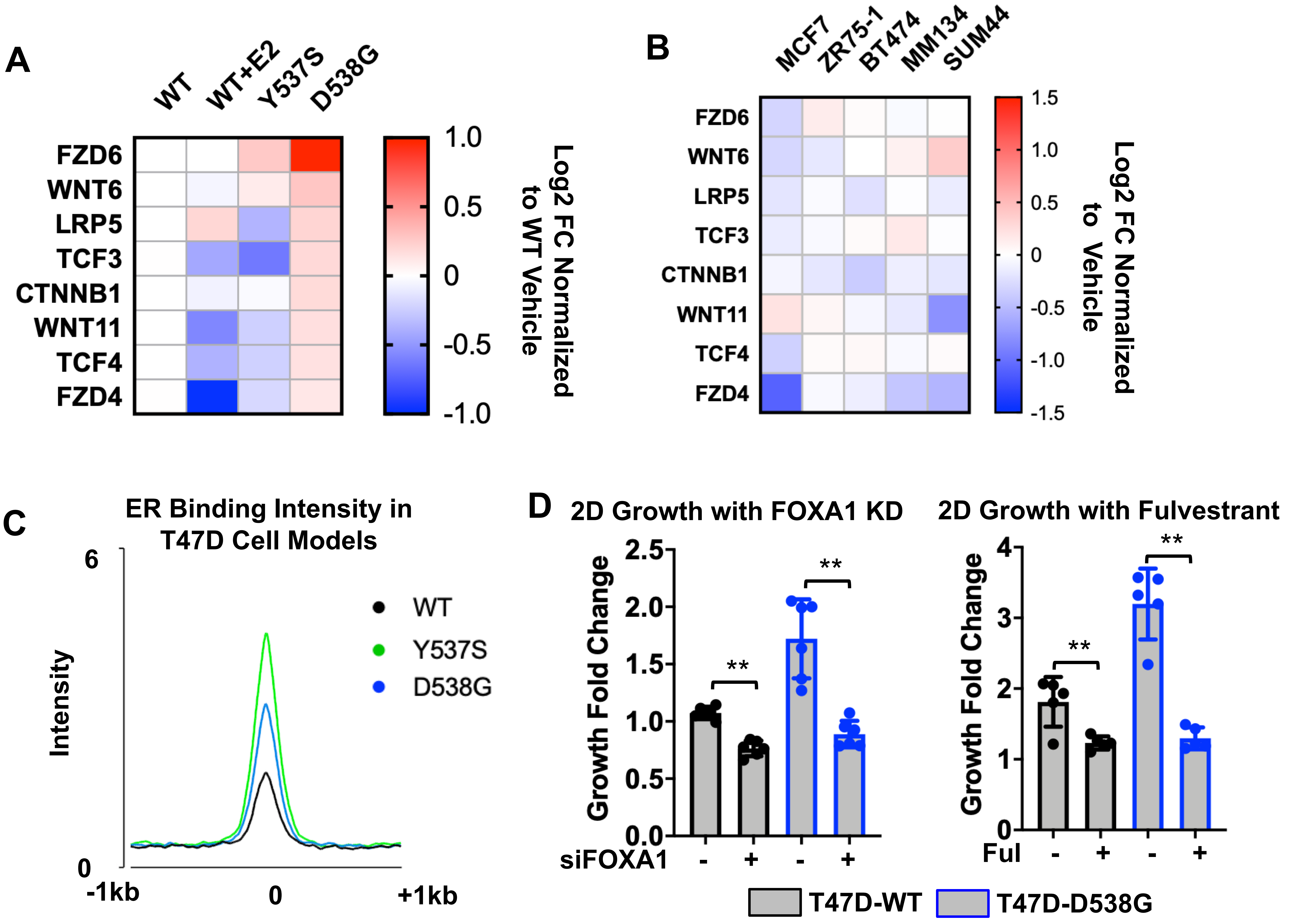


**Figure S19. Target Wnt genes are not direct targets of ER. (Related to Fig. 5)**
A. A heatmap showing $log_2$ fold changes (normalized to WT-vehicle group) of the eight selected Wnt target genes in T47D cell models. Expression data was extracted from RNA-seq performed with four biological replciates.
B. A heatmap representing the expression fold changes (normalized to each vehicle treatment counterpart) of the eight Wnt regulator genes under E2 treatment in 5 ER+ breast cancer cell lines from publicly available datasets. (GSE89888, GSE3834, GSE38132 and GSE50693).
C. Average binding intensities towards all the binding sites from WT, Y537S and D538G in the absence of E2 of each cell lines in a window of ± 2kb from the peak center. Binding intensities are normalized to the WT-E2 region sets.
D. Bar plots showing T47D WT and T47D-D538G cell growth fold changes dat Day 6 (normalized to day 0) following FOXA1 knockdown or 1μM Fulvestrant treatment. This experiment was done once with six (FOXA1 KD) or five (Fulvestrant treatment) biological replicates per group. Student’s t test was used to compare the effect of each treatment in each cell type. (** p<0.01)

### Supplementary Figure S20

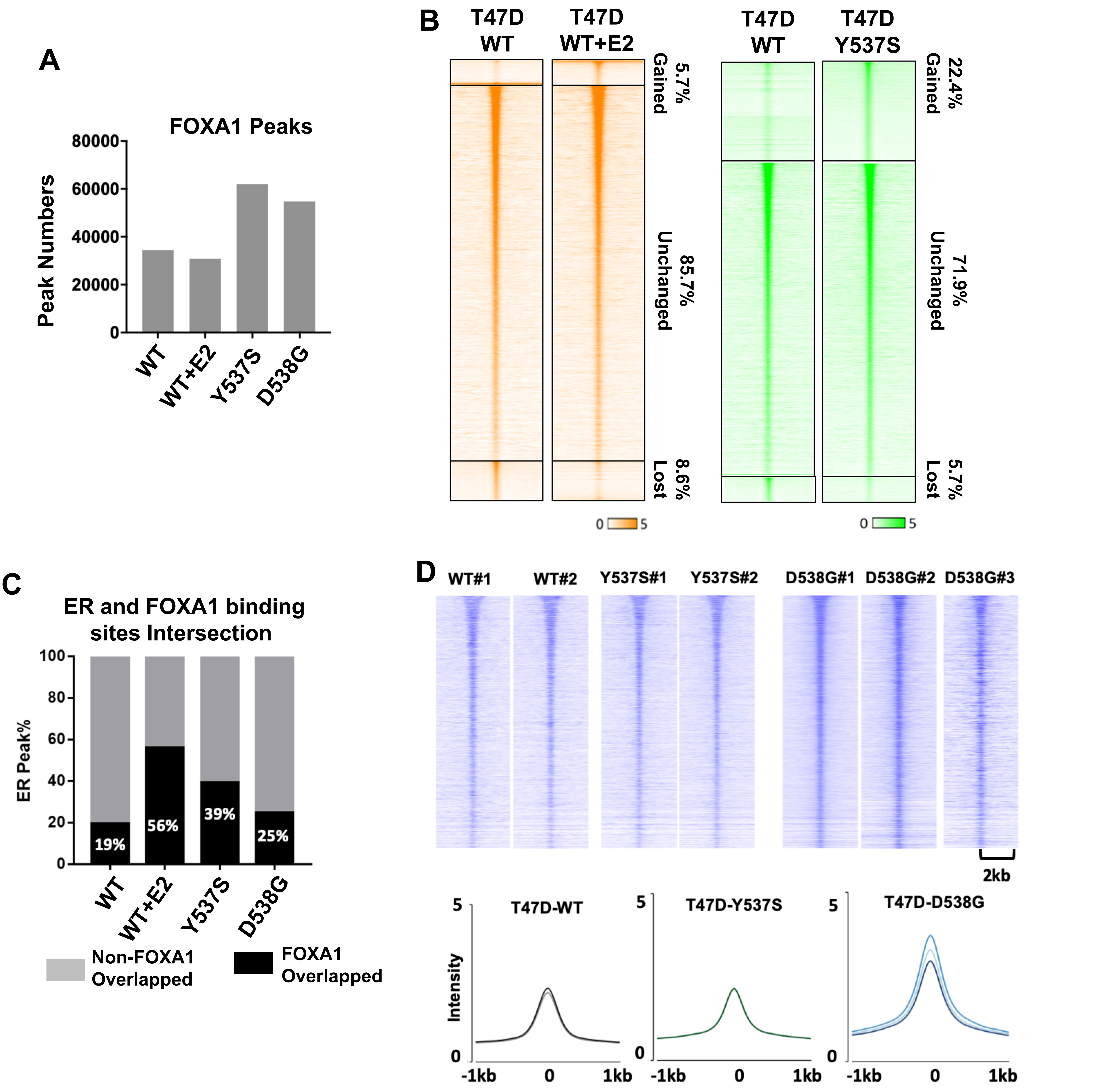


**Figure S20. FOXA1 redistribution rewires accessible genomes in T47D *ESR1* mutant cells. (Related to Fig. 5)**

A. Bar plot showing the called FOXA1 peak numbers in each cell line using MACS2 (q<0.05).

B. Heatmaps of differential FOXA1 binding intensities in T47D-Y537S and WT+E2 groups compared to FOXA1 in WT cells. Displayed in a horizontal window of ± 2kb from the peak center. Pairwise comparison between WT and mutant samples was performed to calculate the fold change (FC) of intensities. Binding sites were sub-classified into sites with increased intensity (FC>2), decreased intensity (FC<-2), and non-changed intensity (-2<FC<2). Percentages for each subgroup are labelled on the heatmaps.

C. Ctacked bar charts showing the percentage of ER peaks overlapping (balck) or not overlapping (grey) with FOXA1 binding sites in all four groups. Percentage of overlapped ER peaks are indicated on each bar.

D. Top panel: Heatmaps showing FOXA1 binding intensities on ATAC-seq detected sites in each individual T47D clone cell model, visualized in a window of ± 2kb from the peak center. Bottom panel: Quantification of FOXA1 binding intensities on detected ATAC sites in a window of ± 2kb from the peak center.

Supplementary Figure S21

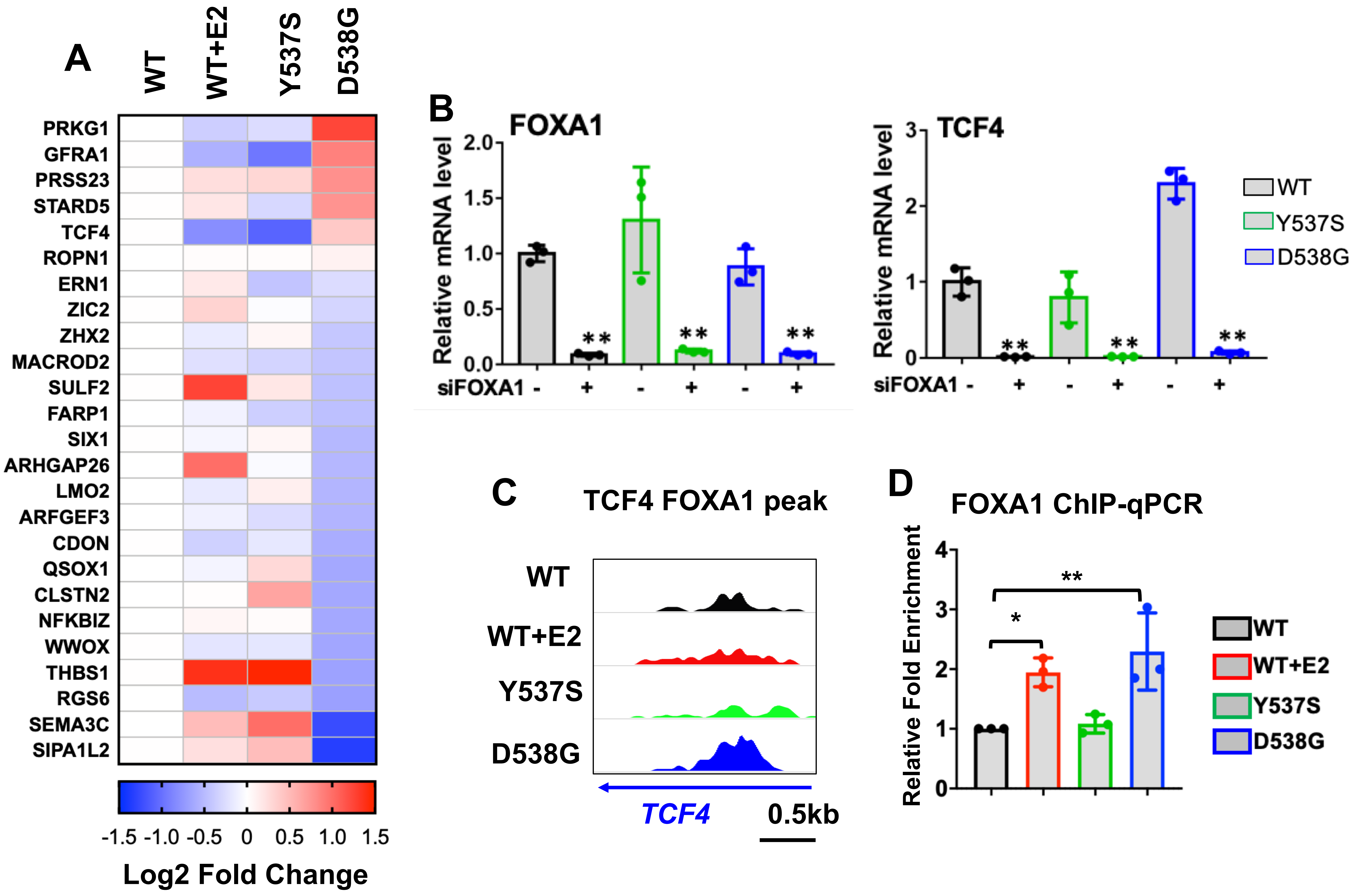


**Figure S21.** ***TCF4*** **upregulation is associated with the uniquely gained FOXA1 binding site in T47D-D538G cells. (Related to Fig. 5)**

A. Heatmap showing $\log_2$ fold changes of the 25 ATAC/FOXA1 gained binding sites annotated T47D-D538G non-canonical ligand-independent genes normalized to WT vehicle group. RNA-seq data (GSE89888) performed with four biological replicates was used.

B. Bar graphs representing qRT-PCR measurement of *TCF4* mRNA levels in MCF7 WT and *ESR1* mutant cells after siRNA knockdown of *FOXA1* for 3 days. ΔΔCt method was used to analyze relative mRNA fold changes normalized to WT cells and *RPLP0* levels were measured as an internal control. Each bar represents mean ± SD with three biological replicates. Representative experiments from three independent repeats is displayed. Student's t test was used to compare the gene expression between scramble and knockdown groups of each cell type. (** $p<0.01$)

C. Screenshot of the gained FOXA1 binding sites at the *TCF4* genomic locus. Y axises represent binding intensity of ChIP-seq and are normalized to the same scale.

D. Bar graphs representing ChIP-qPCR measurements of FOXA1 binding on the gained putative peak (shown on C) proximal to the*TCF4* gene in T47D *ESR1* WT and mutant cells. The relative fold enrichments were calculated by normalizing the Ct values to IgG cotrol groups and further normalized to WT-vehicle group. Each bar represents mean ± SD merged from three independent experiments. Dunnett's test was performed to compare the FOXA1 fold enrichment between the WT vehicle and each other group. (* $p<0.05$, ** $p<0.01$)
