## Supplementary material for "Hotspot *ESR1* mutations are multimodal and contextual drivers of breast cancer metastasis": Supplementary Materials and Methods_ESR1_Mut_Metastasis_Final.docx

**Human tissue studies from the Womens Cancer Research Center (WCRC) and Charite cohorts**

All patients enrolled were approved within IRB protocols (PRO15050502) from the University of Pittsburgh and Charite Universitaetsmedizin Berlin. Informed consent was obtained from all participating patients. Biopsies were obtained and divided into distant metastatic or local recurrent tumors. Genomic DNA was isolated from formalin fixed paraffin embedded (FFPE) samples using Qiagen’s All-prep Kit (#80234). *ESR1* mutation status was detected with droplet digital PCR (ddPCR) targeting Y537S/C/N and D538G mutations in pre-amplified *ESR1* LBD products as previously reported (1). cDNA samples synthesized from RNA were used for ddPCR screening in three local recurrent samples due to poor DNA quality. For the recurrence free survival comparison, patients with RFS=0 were excluded.

For the 54 ER+ metastatic tumor samples, genomic profiles were determined based on tumor RNA sequencing provided in previous publications (2-4). Briefly, biospecimens were reviewed by a trained molecular pathologist to confirm pathology, quantify tumor cellularity and to highlight regions of relatively high tumor cellularity for macrodissection. RNA was extracted from FFPE tissue using Qiagen’s All-Prep Kit, and library preparation was performed using Illumina’s TruSeq RNA Access Library Preparation protocol. Transcript counts from all samples were quantified with Salmon v.0.8.2 (5) and converted to gene-level counts with tximport (6). The gene-level counts from all studies were then normalized using TMM with edgeR (7). Log_2_ transformed TMM-normalized counts per million [log_2_(TMM-CPM + 1)] were used for analysis.

**CTCs analysis from the NU16B06 Cohort**

A retrospective cohort comprising 151 Metastatic Breast Cancer (BC) patients characterized for CTCs, and ctDNA at the Robert H. Lurie Comprehensive Cancer Center of Northwestern University (Chicago, IL) between 2015 and 2019 was analyzed. Patients’ enrollment was performed under the Investigator Initiated Trial (IIT) NU16B06 independently from treatment line. The overall baseline staging was performed according to the investigators’ choice.

CTC enumeration was performed though the CellSearch™ immunomagnetic System (Menarini Silicon Biosystems), samples were collected through CellSave Stabilizing Tubes and processed via Celltracks Autoprep for EpCAM-based immunomagnetic sorting and subsequent characterization for pan-cytokeratin (CK), DAPI and CD45 cells. The enriched and labeled samples were then reviewed via the Celltracks Analyzer II.

Clustered CTCs were defined as those CTCs that were part of a CTC cluster, patients were then classified based on a > 4 clustered CTCs threshold. Patients with ≥ 5 CTC/7.5 ml of blood were defined as Stage IV aggressive as previously reported. (8)

Mutations in *ESR1* (hotspots D538 and Y537) and PIK3CA (hotspots E453 and H1047) were detected by either ddPCR assay using the QX200 ddPCR System (Bio-Rad) or through the Guardant360™ high sensitivity next-generation sequencing platform (Guardant Health, CA). Two 10-mL of whole blood were drawn for each patient using standard stabilizing tubes (Streck, NE). ctDNA extraction was performed using the QIAamp Circulating Nucleic Acid Kit (Qiagen).

Clinical and pathological variables were reported though descriptive analysis. Continuous variables were described though median and interquartile range (IQR), while categorical variables were reported as frequency. The associations between *ESR1* and PIK3CA status and CTCs enumeration and Clustered CTCs were explored through two-sided Fisher’s exact test.

Overall Survival (OS) defined as time from treatment start until death from any cause. The prognostic role of clustered CTCs was investigated in terms of OS through the Kaplan-Meier method with log-rank test for comparing different subgroups.

Statistical analysis was performed using STATA (StataCorp. (2019) Stata Statistical Software: Release 15.1. College Station, TX: StataCorp LP), JMP (SAS Institute Inc. (2019), version 15. Cary, NC), and R (R Core Team (2019), version 3.6.2. R Foundation for Statistical Computing, Vienna, Austria).

**Cell culture**

Different sources of genome-edited MCF7 and T47D *ESR1* mutant cell models were maintained as previously described (9-11). Briefly, individual clones were maintained in DMEM, supplemented with 10% FBS, 100μg/mL penicillin and 100mg/mL streptomycin, at 37°^C^ in a humidified incubator with 5% CO_2_. Mutation allele frequencies were confirmed using ddPCR. Hormone deprivation was performed for all experiments, unless otherwise stated, by washing and maintaining cells in phenol-red-free IMEM (Gibco, A10488) with 5% (Oesterreich MCF7 cell model) or 10% (other cell models) charcoal-stripped serum (CSS, Gemini, #100-119), twice a day for three days. For genome-edited models with multiple clones (Oesterreich and Gertz models), clones with the same genotypes were equally pooled for subsequent experiments. Other parental cell lines, ZR75-1 (CRL-1500), MDA-MB-134-VI (HTB-23), MDA-MB-330 (HTB-127) and MDA-MB-468 (HTB-132), were obtained from ATCC. BCK4 cells were developed as previously reported (12). Cell lines were maintained in the following media (Life Technologies) with 10% FBS: MDA-MB-468 in DMEM, MDAMB-134 and MDA-MB-330 in a 1:1 ratio of DMEM:L-15, ZR75-1 in RPMI and BCK4 in MEM with nonessential amino acids (Thermo Fisher, #11140050) and insulin (Sigma-Aldrich, #91077C).

**Generation of T47D stably overexpressing *ESR1* mutant cell models**

To generate *ESR1* mutant overexpressing cell models, *ESR1* WT and mutant plasmids, in a pcDNA3.1 backbone, were obtained from Addgene (*ESR1*-HA-WT #49498; *ESR1*-HA-Y537S #49499; *ESR1*-HA-D538G #49500; Empty vector #V790-20). T47D parental cells maintained in RPMI supplemented with 10% FBS were transfected with each of the plasmids and selected with 500μg/ml G418 (Thermo Fisher, #10131035) for 3 weeks. G418 treated media was changed every 3 days during the selection process. Overexpression of WT and mutant ER was further validated by immunoblot and ddPCR in pooled clones and used for further experimentation.

**Short term CTC cluster assessment**

4-week old female nu/nu athymic mice were ordered from The Jackson Laboratory (002019 NU/J) according to University of Pittsburgh IACUC approved protocol #19095822. MCF7 WT and mutant cells were stably labelled with RFP-luciferase by infection with the pLEX-TRC210/L2N-TurboRFP-c lentivirus plasmid. Labelled cells were hormone deprived and resuspended in PBS at a final concentration of 10^7^ cells/ml. 100μl of cell suspension was then injected into nude mice with 6 mice per group via an intracardiac left ventricle injection. Post-injected mice were immediately imaged using the IVIS200 in vivo imaging system (124262, PerkinElmer) after D-luciferin intraperitoneal injection to confirm successful cell delivery into the circulation system. All mice were euthanized after one hour of injection and their whole blood were extracted via cardiac puncture and collected into CellSave Preservative Tubes (#790005, CellSearch). Blood samples were mixed with 7ml of RPMI media and shipped to University of Minnesota for CTC enrichment. Specifically, 1mL of 10% formalin is added to the blood and the tube is rocked for 4 minutes. 5mL of blood is placed in an assembled faCTChecker filtration cassette and an additional 5mL of PBS is added. The blood is filtered with the manual, 10mL filtration setting. 10mL of PBS is added to the cassette and filtered to wash remaining blood cells from the filter. Additional washes are performed until blood is no longer visible on the filter. Filtration slide is removed from the cassette and the plastic backing and sticker are put in place beneath the filter. CTCs were stained with antibody against pan-cytokeratins (CK) conjugated Alex Fluor 488 (Thermo Fisher, #53-9003-82) and DAPI during mount using ProLong Gold (Thermo Fihser, P36935). Slides with stained CTCs were manually scanned in a blind manner and all visible single CTCs or clusters were imaged under 5X or 40X magnifications respectively. To set up criteria for identifying CTC clusters via images, we analyzed seven single CTCs with intact CK signal distribution and calculated the average nuclei-edge to membrane distance (x). Inter-nuclei-edge distance greater than 2x for any two CTCs were excluded in CTC cluster calling. All measurements were performed in a blind manner.

**qRT-PCR**

MCF7 and T47D cells were seeded in triplicates into 6-well plates with 120,000 and 90,000 cells per well respectively. After desired treatments, RNA was and cDNA was synthesized using iScript kit (#1708890, BioRad, Hercules, CA). qRT-PCR reactions were performed with SybrGreen Supermix (#1726275, BioRad), and the ΔΔCt method was used to analyze relative mRNA fold changes with *RPLP0* measurements serving as the internal control. All primer sequences can be found in Supplementary Table S10.

For ECM molecules, RT^2^ Profiler PCR array experiments were performed followed the provided manufacturing protocols from Qiagen (PAHS-013Z) with three biological replicates. Cq values were normalized against the GEO database mean of five house-keeping genes, and the expression fold change (FC) was calculated via comparison of *ESR1* mutant and *ESR1* WT expression levels. To select significantly altered genes, a student’s t test was performed to each pair of WT and mutant cells with p-values adjusted for multiple comparisons. Significant differentially expressed genes were filtered with a threshold of Cq<35 and a q-value<0.1 in at least one representative group of each cell line.

**Immunoblotting**

After desired treatments, cells were lysed with RIPA buffer spiked with a fresh protease and phosphatase cocktail (Thermo Scientific, #78442) and sonicated. Protein concentrations were quantified using the Pierce BCA assay kit (Thermo Fisher, #23225). 80-120µg of protein for each sample was loaded onto SDS-PAGE gels, and then transferred onto PVDF membranes. The blots were incubated with the following antibodies: desmocollin 1 (sc-398590), desmoglein 2 (sc-80663), plakophilin (sc-33636), connexin 26 (sc-7261) and cFOS (sc-52) from Santa Cruz; ER-α (#8644), HA (#3724), Non-phospho-β-catenin (#19807), Histone H3 (#4499), AIF (#5318), GSK3β (Ser9, #5558), phospho-GSK3α (Ser21, #9316), GSK3β (#12456) and GSK3α (#4337) from Cell Signaling Technology; β-catenin (#610154) from BD; Tubulin (T6557) and connexin 43 (C6219) from Sigma Aldrich; and *TIMP3* (ab39184) from Abcam.

**Immunofluorescence staining**

MCF7 cells were hormone deprived as previously described and seeded on coverslips. After attachment, cells were fixed with 4% paraformaldehyde and blocked with 3% BSA solution plus 0.1% tritonX-100. Primary antibody against desmoglein 2 (Santa Cruz, sc-80663) was applied to stain the cells followed by secondary FITC-conjugated antibody (Thermo Scientific, A16079) and Hoechst staining (Thermo Scientific, #62249). For F-actin filament staining, Alexa 488-conjugated Phalloidin was used (Thermo Scientific, #12379). Coverslips were mounted and images were taken using a fluorescence microscope (Olympus, CZX16) under the 20X objective. Quantification was performed by dividing the integrated intensity of FITC signals by the number of nuclei in each image. Data from 20 sections of images were combined for analysis, collected from a total of four independent experiments. The results reported are from all four independent experiments combined, with scoring performed blindly.

For the immunofluorescence staining of mice lung sections evaluating micro-metastases, 5-micron FFPE slides of each sample were first deparaffinized and rehydrated. The slides were then subjected to an antigen retrieval step by boiling the slides in a citrate buffer (16mM sodium citrate, 4mM citrate acid monohydrate, pH=6.0) using a high-pressure cooker for 20 minutes. Upon cooling, the lung sections were cycled using PAP pen and rinsed in PBST for 5 minutes. The slides were incubated in a 100mM glycine solution to reduce background staining as well as incubated in a blocking buffer (0.3% Triton X-100, 5% goat serum in PBS) for one hour. Primary antibodies were mixed in blocking buffer and applied on all slides overnight at 4°^C^ (Human CK19: Thermo Fisher 190-P1; CK8+18: Abcam #53280). Slides were further washed three times with PBST and incubated in secondary antibody for 1 hour at room temperature (Alexa Fluor 488: Thermo Fisher A32723; Alexa Fluor 568: Thermo Fisher A11011). Hoechst staining was applied for nuclei visualization, and all the slides were mounted and imaged using a fluorescence microscope (Olympus, CZX16). Two representative regions of each slide were selected for blind quantification.

**Cell growth assay**

3,000 MCF7 or 4,000 T47D cells were seeded into either flat bottom 96-well ultra-low attachment plates (for 3D growth) (Corning, #3474) or regular 96-well plates (for 2D growth) (Corning, #353072). Cell numbers were quantified after the desired growth time course with either the Celltiter Glo luminescent cell viability kit (Promega, G7573) or the FluoReporter Blue Fluorometric dsDNA quantification kit (Invitrogen, F2962). Fluorescent readouts were corrected to background measurements.

**Pan-MMP activity assay**

A FRET-based MMP activity assay was performed using the MMP activity assay kit (Abcam, ab112146), following the manufacturing protocols. In brief, 25µg of protein from whole cell lysates was pre-mixed with APMA. MMP substrates were then loaded and fluorescence intensities were monitored under the excitation and emission wavelength of 490/525nm after 0, 5, 10, 15, 20, 25, 30, 40, 50 and 60 minutes of incubation. Background emissions were corrected, and fold changes were calculated by normalizing to initial readout.

**Spheroid invasion and collective migration assay**

Spheroid invasion assay was performed as previously described (15). 3,000 MCF7 or 4,000 T47D cells were seeded into 96-well round bottom ULA plate for a 2-day spheroid formation. For the invasion assay, 1.3mg/ml type I collagen (Corning, #354236) supplemented with 1% NEAA (Thermo Fisher Scientific, #11140050) was directly added into each well. Images were taken at day 0 and day 6 (T47D) or day 4 (MCF7). Invasion areas were quantified by subtracting day 0 areas from day 6 or 4 areas using Adobe Photoshop. For the collective migration assay, spheroids were gently transferred into a collagen coated 96-well plate in the presence of 5µg/ml Mitomycin C. Spheroid migration images were taken at day 0 and day 4 and migratory distances were calculated using the mean values of spheroid weights and heights normalized to day 0.

**Quantitative microfluidic fluorescence microscope imaging system**

qMFM assays were performed as previously described (16). Cells labelled with calcein were seeded onto coverslips for 24 hours. The same cell types were labelled with DiD cell staining solution (Thermo Fisher Scientific, V22887) and loaded into the microfluidic system under the shear stress of 6dynes/cm to allow for attachment. Videos were recorded for 200 frames with 100ms/frame. Nikon Element Software was used for quantification, with a module calculating cell movement slower than 3µm/s per frame. Adhered cells from T0 were subtracted, and cell-cell interaction events were normalized to the total calcein signals of each video.

**Calcein dye transfer assay**

MCF7 WT and mutant cells were trypsinized and labelled with 1µM calcein for 30 minutes. 10,000 labelled and unlabeled cells were mixed 1:1 and incubated in a Corning round bottom tube (#352054) for 12 hours. Cells were then subjected to flow cytometry analysis gating on single cell populations. GFP positive cells were analyzed by setting the threshold with positive and negative controls. Exchanged dye ratios were calculated by doubling the GFP+ cell percentages and subtracting 50%.

**Top-Flash luciferase assay**

125,000 T47D cells were seeded into a 12-well plate. M50 Super 8X Top-Flash luciferase (Addgene, #12456) and Renilla plasmids were co-transfected into all groups as previous described (17). Cells were lysed after 24 hours of transfection and luciferase values were measured with the Dual-luciferase Reporter Assay system (Promega, E1910). Relative Wnt activity was calculated as the ratio of firefly luciferase activity over internal control Renilla luciferase activity.

**Boyden chamber transwell assay**

For the chemotaxis assay, the QCM chemotaxis cell migration kit (Millipore Sigma, ECM508) was used, following the manufacture’s instruction. Briefly, T47D cells were hormone deprived and starved in serum free medium for 24 hours. Cells were digested and diluted to 10^6^ cells/ml. 300μl of cell suspensions were loaded, in duplicate, in the inner chamber of 8μm cell inserts. 10% CSS or serum free medium was loaded in the outer chamber. After 72 hours, cells were stained with 0.1% crystal violet and cells in the inner chamber were wiped using cotton swabs. The inserts were imaged and any remaining cells were then dissolved in extraction buffer and subjected to colorimetric measurement under OD450.

**Chromatin-immunoprecipitation (ChIP)**

ChIP experimentation was performed as previously described (18). The immunoprecipitation was performed using ERα (sc543) and rabbit IgG (sc2027) antibodies (Santa Cruz Biotechnologies). Histone 3 acetylation at K27 site (ab4739) and Histone 3 di-methylation at K4 site (ab7766) and FOXA1 (ab23738) antibodies were obtained from Abcam.

For ER ChIP-seq, DNA samples were pooled from individual clones with the same genotype and at least 10ng of DNA from each sample was sent to McGill University Sequencing Core for library preparation using TruSeq ChIP Library Preparation Kit (Illumina IP-202) and sequencing using the Illumina Hiseq 2500 Platform (ER ChIP-seq). Over 16M single end 50bp reads were allocated for each sample.

For FOXA1 ChIP-seq, DNA samples were originally prepared from pooled cells in biological triplicates, and replicates of each group were further combined to ensure sufficient input amount for subsequent process. The pooled samples were sent to the Health Sciences Sequencing Core at Children’s Hospital of Pittsburgh for library preparation using TruSeq ChIP Library Preparation Kit (Illumina IP-202) and sequencing using Illumina NextSeq platform. Over 80M single end 150bp reads were allocated for each sample.

For ChIP-qPCR, DNA was diluted and subjected to qRT-PCR as described above. The fold enrichment method was used to quantify binding enrichment at selected sites. IgG-IP’d samples were used as negative controls. The primer sequences for targeted genomic regions are reported in Supplementary Table S8.

**ChIP-sequencing Analysis**

ChIP-seq reads were aligned to Hg19 reference genome assembly using Bowtie 2.0 (19), and peaks were called using MACS2.0 with a p-value<10^-5^ (ER ChIP-seq) or a q-value<0.05 (FOXA1 ChIP-seq) (20). We used the Diffbind package (21) to perform principle component analysis, identify differentially expressed binding sites and analyze intersection ratios with other datasets. Briefly, all BED files for each cell line were merged and binding intensity was estimated at each site based on the normalized read counts in the BAM files. Pairwise comparisons between WT and mutant samples were performed to calculate fold change (FC). Binding sites were sub-classified into three categories: gained sites (FC>2), lost sites (FC<-2), and not-changed sites (2<FC<2). Heatmaps and intensity plots for binding peaks were visualized by EaSeq (22). For gene annotation from FOXA1 binding sites, gained FOXA1 peaks were selected and annotated genes were inspected in a ± 200kb range of the FOXA1 peaks using ChIPseeker (23). For intersection analysis of the D538G-regulated non-canonical ligand-independent genes, broad differentially expressing genes were first called using a cutoff of |fold change|>2, FDR<0.005 between WT and D538G cells. Meanwhile, E2-regulated genes were called using the cutoff of |fold change|>1.5, FDR<0.01 between WT and WT+E2 groups. D538G ligand-independent genes which are also regulated in WT cells were excluded from the analysis.

**RNA-sequencing analysis**

Data generation and processing of the 54 ER+ tumors in the WCRC cohort was described above. For the MET500 cohort, RNA-seq fastq files from 91 metastatic breast cancer samples were downloaded from the Database of Genotypes and Phenotypes (dbGaP) with accession number phs000673.v2.p1. Transcript counts from all samples were quantified with Salmon v.0.8.2 and converted to gene-level counts with tximport. The gene-level counts from all studies were then normalized together using TMM with edgeR. Log2 transformed TMM-normalized counts per million [log_2_(TMM-CPM+1)] were used for analysis. To predict ER positivity based on *ESR1* expression, the TCGA cohort was used as a reference. Briefly, putative “ER+” (higher than a pre-defined cutoff) and “ER-” (lower than a pre-defined cutoff) statuses were predicted based on *ESR1* log_2_(CPM+1) values of 1045 primary tumors using each consecutive interval of 0.1 between 3 (first quartile of *ESR1* expression levels in MET500) and 8.8 (third quartile of *ESR1* expression levels in MET500). The predicted results were then compared to pathological ER status identification for each cutoff selection. Log_2_(CPM+1) values of 5.6 were determined as the final cutoff for ER status due to a highest concordance ratio towards pathological records (95.5%). 46 putative ER positive samples were then filtered in the MET500 cohort. *ESR1* mutation status was extracted using the MET500 portal (<https://met500.path.med.umich.edu>). For the DFCI cohort, raw counts data was obtained and normalized to log_2_(TMM-CPM+1) for further analysis. *ESR1* mutation status was called using separate whole exon sequencing data. For the POG570 cohort, raw count matrixes and mutation statuses were downloaded from the BCGSC portal (<https://www.bcgsc.ca/downloads/POG570/>). ER status of each patient was additionally requested from the cited original resources and only ER+ metastatic tumors were used for downstream analysis. For all three cohorts, *ESR1* mutations were further selected by experimentally validated variants for downstream analysis.

For all datasets, differential expression (DE) analysis was performed using the DESeq2 package (24). In brief, genes were prefiltered with a log_2_CPM>0 criteria across all samples. DE genes with a q-value below 0.1 and an absolute log_2_ fold change above 1.5 were used for Ingenuity Pathway Analysis (25). GSEA analysis was performed using the Broad GSEA Application (26). Gene set variation analyses were performed using the GSVA package (27). All gene sets used in this study are reported in Supplementary Table S5. Data visualizations were performed using “ggpubr” (28) and “VennDiagram” packages (29).

**Statistical Analysis**

GraphPad Prism software version 7 and R version 3.6.1 were used for statistical analysis. All experimental results included biological replicates and are shown as mean ± standard deviation, unless otherwise stated. Specific statistical tests are indicated in corresponding figure legends. All tests were conducted as two-tailed, with a p<0.05 considered statistically significant. Drug synergy was calculated based on the Bliss independence model using the SynergyFinder R package (30). Bliss synergy scores were used to determine synergistic effects.

**Data Availability**

The ER ChIP-seq data has been deposited onto the Gene Expression Omnibus database (GSE125117). Deposition of FOXA1 ChIP-seq has not been performed. All publicly available resources used in this study are summarized in Supplementary Table S11. All raw data and scripts are available upon request from the corresponding author.
